# The extracellular association of the bacterium “*Candidatus* Deianiraea vastatrix” with the ciliate *Paramecium* suggests an alternative scenario for the evolution of *Rickettsiales*

**DOI:** 10.1101/479196

**Authors:** M. Castelli, E. Sabaneyeva, O. Lanzoni, N. Lebedeva, A.M. Floriano, S. Gaiarsa, K. Benken, L. Modeo, C. Bandi, A. Potekhin, D. Sassera, G. Petroni

**Affiliations:** Centro Romeo ed Enrica Invernizzi Ricerca Pediatrica, Dipartimento di Bioscienze, Università degli studi di Milano, Milan, Italy; Department of Cytology and Histology, Faculty of Biology, Saint Petersburg State University, Saint-Petersburg, Russia; Dipartimento di Biologia, Università di Pisa, Pisa, Italy; Centre of Core Facilities “Culture Collections of Microorganisms”, Saint Petersburg State University, Saint Petersburg, Russia; Dipartimento di Biologia e Biotecnologie, Università degli studi di Pavia, Pavia, Italy; UOC Microbiologia e Virologia, Fondazione IRCCS Policlinico San Matteo, Pavia, Italy; Core Facility Center for Microscopy and Microanalysis, Saint Petersburg State University, Saint-Petersburg, Russia; Department of Microbiology, Faculty of Biology, Saint Petersburg State University, Saint-Petersburg, Russia

**Keywords:** Deianiraea vastatrix, Deianiraeaceae, evolution of intracellularity, Rickettsiales ancestor, epibiont, ectosymbiont, Rickettsiales phylogeny

## Abstract

*Rickettsiales* are a lineage of obligatorily intracellular *Alphaproteobacteria*, encompassing important human pathogens, manipulators of host reproduction, and mutualists. Here we report the discovery of a novel *Rickettsiales* bacterium associated with *Paramecium*, displaying a unique extracellular lifestyle, including the ability to replicate outside host cells. Genomic analyses show that the bacterium possesses a higher capability to synthesize amino acids, compared to all investigated *Rickettsiales*. Considering these observations, phylogenetic and phylogenomic reconstructions, and re-evaluating the different means of interaction of *Rickettsiales* bacteria with eukaryotic cells, we propose an alternative scenario for the evolution of intracellularity in *Rickettsiales*. According to our reconstruction, the *Rickettsiales* ancestor would have been an extracellular and metabolically versatile bacterium, while obligate intracellularity and genome reduction would have evolved later in parallel and independently in different sub-lineages. The proposed new scenario could impact on the open debate on the lifestyle of the last common ancestor of mitochondria within *Alphaproteobacteria*.

## Introduction

The “Bergey’s Manual of Systematics of Archaea and Bacteria” states that *Rickettsiales* (*Alphaproteobacteria*) “multiply only inside [eukaryotic] host cells” (Dumler and Walker 2015). Obligate intracellularity indeed represents the main feature of this bacterial order, considering that their host range, metabolic capabilities, and spectrum of effects on the hosts greatly vary (Perlman et al. 2006; Werren et al. 2008; Montagna et al. 2013; Castelli et al. 2016). As a matter of fact, *Rickettsiales* encompass highly diverse representatives, including human pathogens (Kocan et al. 2004; Parola et al. 2013; Weinert et al. 2009), reproductive parasites of arthropods (Werren et al. 2008), mutualists of nematodes (Taylor et al. 2005; Werren et al. 2008) or arthropods (Hosokawa et al. 2010), and several bacteria associated with unicellular eukaryotes, producing undisclosed consequences on their hosts (Castelli et al. 2016). Additionally, *Rickettsiales* are well known to evolutionary biologists, since several studies suggested them as the sister group of mitochondria (Andersson et al. 1998; Fitzpatrick et al. 2006; Wang and Wu 2015), although there is no full agreement on this point (Esser et al. 2004; Abhishek et al. 2011; Thiergart et al. 2012; Martijn et al. 2018).

A common trait of most *Rickettsiales* is the ability to perform horizontal transmission, testified by multiple lines of evidence: lack of congruence in host and symbiont phylogenies (Perlman et al. 2006; Epis et al. 2008; Weinert et al. 2009; Driscoll et al. 2013), several examples of closely related bacteria harbored by unrelated hosts (Kocan et al. 2004; Dantas-Torres et al. 2012; Schrallhammer et al. 2013; Senra et al. 2016; Matsuura et al. 2012), and even experimental host transfers (Braig et al. 1994; Caspi-Fluger et al. 2012; Schulz et al. 2016; Senra et al. 2016). Thus, *Rickettsiales* are “host-dependent”, but, with few exceptions (Casiraghi et al. 2001), not “host-restricted”, because they are not exclusively vertically transmitted.

Therefore, the life cycle of most *Rickettsiales* can be ideally subdivided into two phases. The main phase is intracellular, during which all life functions, including replication, are exerted. During the other phase, which is facultative, the bacteria survive temporarily in the extracellular environment and can enter new host cells (Walker and Ismail 2008; Rikihisa 2010; Schulz et al. 2016). According to our best knowledge, these “transmission forms”, sometimes displaying altered morphology (Philip & Casper 1981; Labruna et al. 2004; Sunyakumthorn et al. 2008), have never been reported to replicate.

Until recently, on the basis of 16S rRNA gene phylogenies, *Rickettsiales* included four families, namely *Rickettsiaceae*, *Anaplasmataceae*, “*Candidatus* (*Ca.*) Midichloriaceae” (from now on, *Midichloriaceae*), and *Holosporaceae*. This classification is still preferred by some authors (e.g Driscoll et al. 2013; Kang et al. 2014), who consider the whole clade monophyletic. However, in the latest years several studies, especially those employing multiple concatenated genes, provided alternative phylogenetic scenarios arguing against the monophyly of *Rickettsiales sensu lato* (i.e. including *Holosporaceae*) (Georgiades et al 2011; Ferla et al. 2013; Wang & Wu 2015; Hess et al. 2016; Parks et al. 2018). Thus, considering also the high sequence divergence, *Holosporaceae* were elevated to the rank of independent order, i.e. *Holosporales* (Szokoli et al. 2016a). In this work, the definition by Szokoli and co-authors (2016a) of *Rickettsiales sensu stricto* (*Rickettsiaceae*, *Anaplasmataceae*, and *Midichloriaceae*) is followed.

While *Rickettsiales sensu stricto* form a strongly supported monophyletic group (e.g. Driscoll et al. 2013; Wang & Wu 2015; Hess et al. 2016; Szokoli et al. 2016a; Floriano et al. 2018), the members of the three families can be distinguished by multiple features, including the host cell entrance mechanism and the presence or absence of a host-derived membrane envelope in the intracellular phase (Walker and Ismail 2008; Rikihisa 2010; Castelli et al. 2016).

Regardless of these differences, the transition to an obligate intracellular state has always been assumed to have occurred only once in *Rickettsiales* evolution, before the separation of the three families (Darby et al. 2007; Weinert et al. 2009; Sassera et al. 2011; Schulz et al. 2016). Accordingly, obligate intracellularity has always been considered an apomorphy of *Rickettsiales*, namely a shared character directly inherited from their common ancestor.

Here we present the first report of an extracellular *Rickettsiales* bacterium, “*Ca*. Deianiraea vastatrix”, displaying an unprecedented extracellular and putatively parasitic lifestyle in the interaction with the ciliate protist *Paramecium primaurelia*. According to the obtained novel morphological, phylogenetic and genomic data, we re-evaluated traditional hypotheses on the evolution of *Rickettsiales*, which always implied an “intracellularity early”, dating back to the common ancestor. Thus, based on the here presented findings, we propose an alternative scenario, named “intracellularity late”, under which we suggest that intracellularity could have evolved in parallel and independently along multiple *Rickettsiales* sub-lineages from an extracellular, yet host-associated, ancestor.

## Results

### Live and electron microscopy observations

A monoclonal culture of *P. primaurelia* cells (strain CyL4-1) was obtained starting from a sample of waste stream water collected in Larnaca – Cyprus. After one month of laboratory cultivation of the strain, many dead cells were registered in the culture. Microscopic observations revealed an abnormal phenotype of paramecia: shortened ovoid-rounded shape and immotility caused by a nearly complete loss of cilia, which in normal paramecia fully cover the cell and are employed for locomotion and feeding (Supplementary material 1). The deciliated areas of the cell surface were covered by a dense overlay, formed by one to multiple layers of tightly packed intermingled bacteria (Figure 1a; Figure 2a; Supplementary material 2), in a fashion somehow reminiscent of the arrangement of episymbiotic bacteria covering the surface of ciliates from oxygen-depleted environments (Modeo et al. 2013; Nitla et al 2018). The thickness of the overlay looked roughly proportional to the level of degeneration in the shape of the ciliates. These bacteria appeared uniform in their morphology, presenting the typical double-membrane cell structure of Gram-negatives, and were slightly curved rods, sized approximately 1.6-1.7×0.1-0.2 μm. In longitudinal sections, the bacterial cells appeared to be progressively narrowing on one end, with a sharp apical tip, which sometimes took directly contact with the external side of the *Paramecium* cell membrane (Figure 2b,c). The bacterial cytoplasm was electron-dense, containing numerous ribosomes. Within the bacterial population, a small portion of dividing cells was regularly observed (Figure 2d). Despite the loss of most cilia of the affected *Paramecium* cells, neither the intracellular ciliary basal bodies were altered in number and structure, nor the external cell membrane was damaged in its integrity. However, other features potentially linked to the pathological state were observed, such as numerous cytoplasmic lysosomes and ring-like nucleoli, the latter being a typical trait in starving cells (Supplementary material 2). Other organelles and subcellular elements, such as trichocysts, mitochondria, or food vacuoles, did not present any appreciable alterations. Moreover, no intracellular bacteria were ever observed aside from those in food vacuoles.

**Figure 1.**
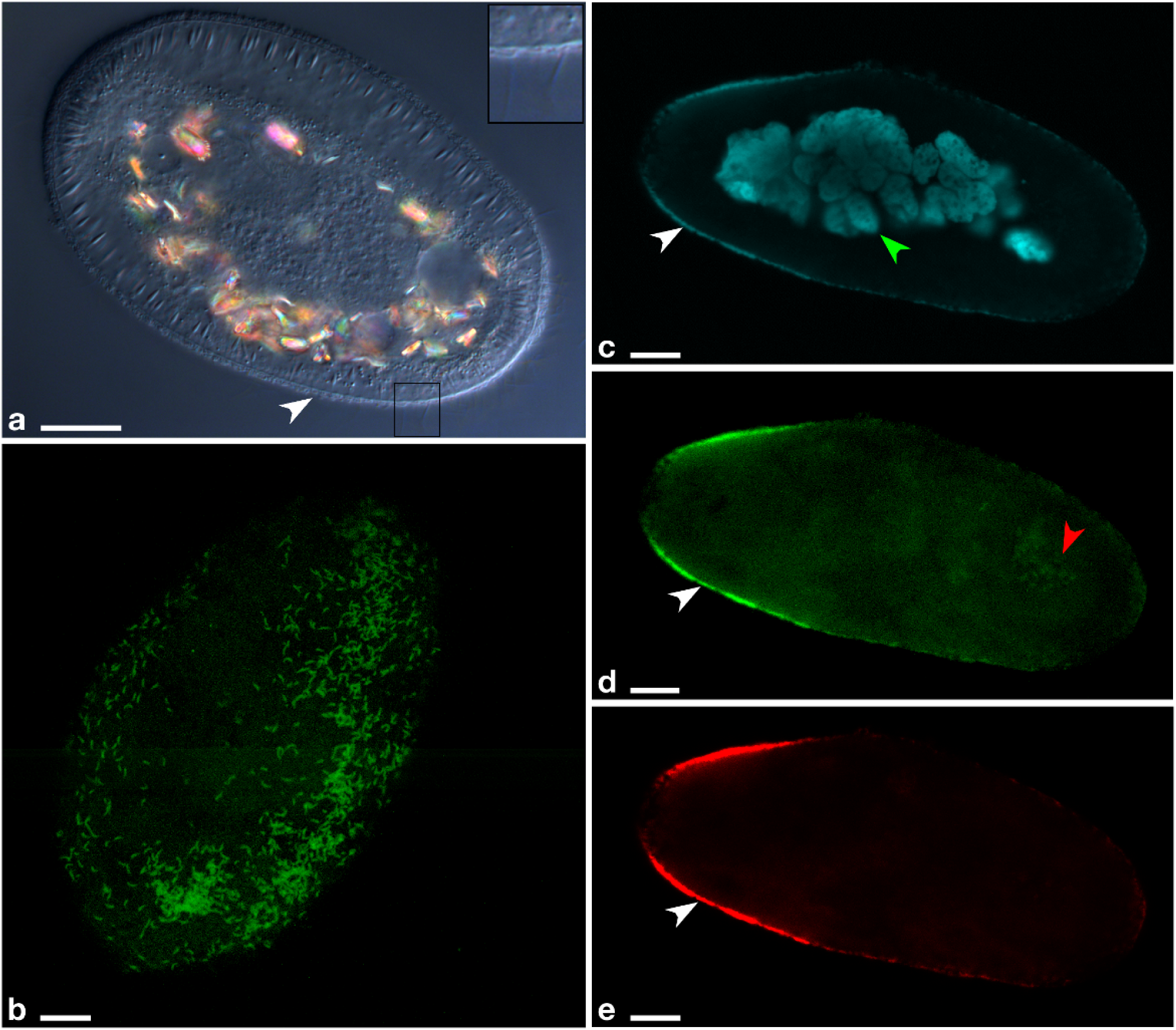
Microscopy images of *P. primaurelia* CyL4-1 covered by extracellular *Deianiraea* bacteria. **a,** Differential interference contrast (DIC) of a heavily infected CyL4-1 cell, displaying an atypical shortened and ovoid shape. The *Paramecium* cell surface is covered by a bacterial multilayer (white arrowhead). As shown in the enlarged inset on the right (corresponding to the square in the main picture) the ciliature is highly reduced, with only few remnants. **b,** Fluorescence *in situ* hybridization microscopy image of the surface of a *Deianiraea*-covered CyL4-1 cell. *Deianiraea* cells are visible by the green signal of the FITC (fluorescein isothiocyanate)-labelled universal bacterial probe EUB338 (Amann et al. 1990). **c-e,** Multichannel microscopy images showing a transversal plan from a FISH experiment on a CyL4-1 cell at an advanced stage of invasion by the putative parasite. In (**c**) the blue signal from DAPI (4′,6-diamidino-2-phenylindole) highlights the *Paramecium* fragmented macronucleus in autogamy (green arrowhead) and the bacterial cells. In (**d**) the green signal of FITC-labelled EUB338 is shown; in (**e**) the red signal of the *Deianiraea*-specific probe Deia_416 labelled with Cy3. In (**c-e**), the *Deianiraea* cells covering the external surface of the host cell on its outline are fluorescently stained (white arrowheads). In (**d**), food bacteria in a digestive vacuole are also visible (red arrowhead), without any corresponding signal from the specific probe in (**e**). Bars stand for 10 μm

**Figure 2.**
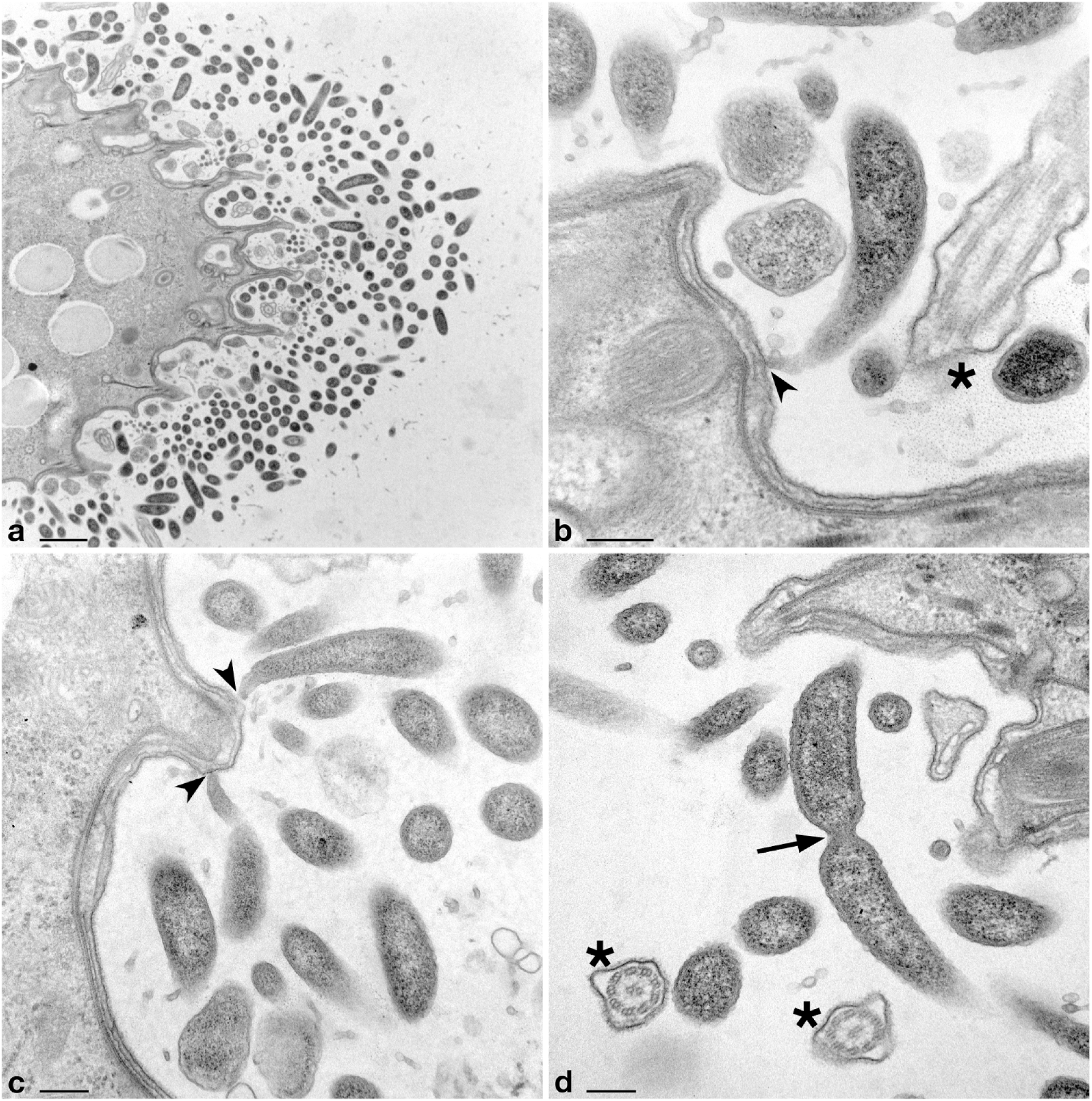
Transmission electron microscopy images of *Deianiraea* bacteria on the surface of *P. primaurelia* CyL4-1 cells. **a,** Portion of *Paramecium* cell surrounded by a huge number of extracellular *Deianiraea* bacteria, forming multiple layers. Bacterial cells appear electron dense, irregularly arranged, and sectioned in variably oriented planes. **b-d,** Details of *Deianiraea* cells at higher magnification. In (**b**) and (**c**) longitudinal sections of some *Deianiraea* cells are shown. The bacterial side proximal to the *Paramecium* cell progressively narrows, ending up in a sharp and slightly curved apical tip, which takes direct contact with the external side of the *Paramecium* cell membrane (black arrowheads). In (**d**) a dividing *Deianiraea* cell, with the division septum highlighted by a black arrow. The few residual *Paramecium* cilia are evidenced by black asterisks in (**b**, **d**) sectioned from variable angles. Scale bars stand for, respectively, 1 μm (**a**) and 200 nm (**b-d**).

The extracellular bacteria attached to the *Paramecium* surface propagated rapidly within the culture, spreading the abnormal phenotype to other ciliate cells. Survival of the ciliate population was possible only by operating a regular (every 2-3 days) dilution of the load of the associated bacteria, i.e. transferring *Paramecium* cells into fresh medium containing food bacteria. Untreated (i.e. non-transferred) host cells invariably died within maximum 4 days. The applied procedure allowed laboratory maintenance for nearly eleven months of sub-replicates of the original *Paramecium* cultures, harboring the bacteria. During such period of time, the *Paramecium* cells and associated bacteria were regularly monitored (approximately 100 cells every week), and always presented the above described traits. The observed behavior led us to consider the surface-attached bacteria as probable parasites of the *Paramecium*, although direct experimental tests would have been necessary to fully prove this inference. Consistently, a sub-replicate, in which the prolonged regular transferring allowed to completely remove the extracellular surface-attached bacteria, was successfully maintained in culture with no sign of the altered phenotype. This sub-strain, free from the putative parasite, was named CyL4-1*.

### The putative parasitic bacteria can colonize the extracellular surface of other paramecia

The strain CyL4-1* and six other laboratory *Paramecium* strains were tested for susceptibility to the putative parasitic bacteria from CyL4-1 in experimental shift assays, using as carriers highly infected, dying CyL4-1 cells. The putative parasitic bacteria successfully colonized the surface of CyL4-1*, and all *Paramecium* cells perished within the 4^th^ day of the experiment, showing the same phenotype as in the original CyL4-1 strain. Unexposed, control cells were viable and proliferated regularly.

The transfer of the putative parasites to other naïve strains of *Paramecium* was attempted by the same procedure. This experiment showed a variable level of resistance to the putative parasite, with several strains being fully untouched, while others were affected, albeit more slowly than CyL4-1* (Supplementary material 3).

### *Deianiraeaceae*, a new family of order *Rickettsiales*

A single almost full-length 16S rRNA gene sequence (1431 bp) was obtained from CyL4-1 strain by PCR with universal bacterial primers followed by direct sequencing, thus indicating the uniformity of the bacterial population. The sequence showed the highest similarities with uncultured sequences of *Rickettsiales*-related bacteria, with best Blastn hit against a bacterium from nephridia of the earthworm *Eiseniella neapolitana* (82.7% identity; JX644354; Davidson et al. 2013). The use of newly designed specific probes in fluorescence *in situ* hybridization (FISH) experiments demonstrated that the characterized sequence belonged to the novel putatively parasitic bacterium (named “*Ca*. Deianiraea vastatrix”, from now on *Deianiraea vastatrix*, or simply *Deianiraea*. Taxonomic description at the end of the Discussion) and clearly showed that it constituted the entire external layers around the *Paramecium* cells (Figure 1b-e; Supplementary material 4). Conversely, *Deianiraea* was absent in food vacuoles, andnever found inside the ciliate cells, confirming the extracellularity of this bacterium.

Phylogenetic analyses showed that *Deianiraea* is a member of *Rickettsiales sensu stricto,* and belongs to a novel divergent family-level clade with 12 other uncharacterized database sequences. This clade is fully supported in the analyses (100 ML; 1.00 BI; Figure 3; Supplementary material 5), and will be referred from now on as *Deianiraeaceae*. Within *Rickettsiales*, *Deianiraeaceae* formed a strongly supported group together with the families *Anaplasmataceae* and *Midichloriaceae* (97 ML; 1.00 BI). Although the precise relationships among those three lineages were unresolved, they all appeared equally supported and at the same rank.

**Figure 3.**
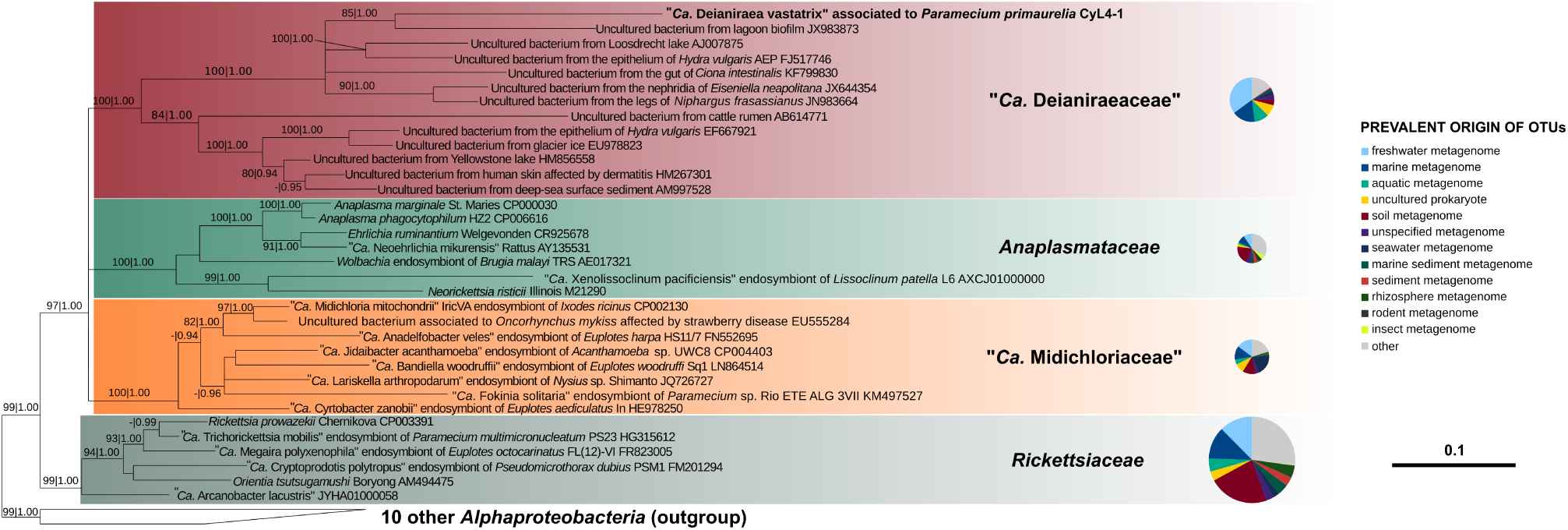
Bayesian phylogenetic tree on 1,398 16S rRNA gene positions on 34 bacteria of the order *Rickettsiales* with the GTR + I + G substitution model. Posterior probabilities (after 1,000,000 generations) and maximum likelihood bootstrap values (with 1,000 pseudo-replicates) of each internal tree nodes are shown on branches (values below 0.85|70 were omitted). The newly characterized *Deianiraea*, putative parasite of *P. primaurelia* CyL4-1 is evidenced in bold. The four *Rickettsiales* families (*Rickettsiaceae*, *Anaplasmataceae*, *Midichloriaceae*, and the herein newly proposed *Deianiraeaceae*) are highlighted by colored boxes. The outgroup, composed by other ten *Alphaproteobacteria*, is shown collapsed as a trapezoidal shape. The scale bar stands for estimated 0.10 sequence divergence. On the right side, for each family the pie chart shows the proportion of prevalent environmental origins (according to the IMNGS categories) of the OTUs obtained after IMNGS database queries with the representative sequence set of the family. The relative areas of the charts are proportional to the total amount of OTUs of the respective families. Only environmental origins that are the prevalent for at least 3% of total OTUs assigned to one family were displayed.

Overall, the 16S rRNA gene sequence identity within *Deianiraeaceae* and compared to other *Rickettsiales* are respectively 77.4-100% and 75.9-86.0% (Supplementary material 6). Two major subclades were found within *Deianiraeaceae* (Figure 3). The sequence identities within subclade 1 (including *Deianiraea*) are ≥81.5%, and within subclade 2 ≥81.1%, while between the two subclades they range between 77.4 and 81.8%.

The sequences of other representatives of *Deianiraeaceae* are from uncharacterized bacteria, mostly from aquatic environments, both freshwater and marine, in several cases associated with diverse eukaryotes, including cnidarians (*Hydra vulgaris*: subclades 1 and 2), ascidians (*Ciona intestinalis*), earthworms (*E. neapolitana*) and amphipods (*Niphargus frasassianus*) (subclade 1) (Fraune & Bosch 2007; Fraune et al. 2010; Bauermeister et al. 2012; Davidson et al. 2013; Dishaw et al. 2014). Other members of the clade were found in association to terrestrial animals, such as in cattle rumen, and, notably, on human skin of dermatitis-affected subjects (Kong et al. 2012; Shinkai et al. 2012) (subclade 2) (Figure 3).

In order to evaluate the distribution and abundance of members of the novel family, a screening was performed on the IMNGS platform (integrated microbial NGS) (Lagkouvardos et al. 2016). The family *Rickettsiaceae* displayed the highest total number of retrieved OTUs, followed by *Deianiraeaceae*, *Midichloriaceae*, and *Anaplasmataceae* (Figure 3; Supplementary material 7). In terms of relative environmental origin (according to IMNGS categories), *Rickettsiaceae* and *Anaplasmataceae* displayed similar patterns, with soil as the prevalent source (>20% OTUs), and several other OTUs from different water origins. On the contrary, *Midichloriaceae* and *Deianiraeaceae* OTUs derived mostly from aquatic environments at variable salinities, with the prevalence of seawater or marine for *Midichloriaceae* and freshwater for *Deianiraeaceae*.

### Genome features

The genome of *Deianiraea* is formed by a single circular replicon, ~1.2 Mbp long. Its size and GC content (32.9%) are consistent with the range of known *Rickettsiales.* The overall coding percentage is ~92.2%, by over two percentage points the highest among sequenced *Rickettsiales* (Supplementary material 8). The genome includes 1129 ORFs, as well as 34 tRNAs for all 20 amino acids, a single rRNA operon (split with 16S rRNA gene in one locus and 23S-5S rRNA genes in another, as in all *Rickettsiales*).

A relatively limited number of mobile elements or their remnants were found, namely 3 transposases (2 complete and 1 partial), two putative prophages (17 total phage genes), plus 13 other phage genes scattered in the genome (Supplementary material 9, 10). The respective best blastp hits on NCBI nr proteins were quite divergent (Supplementary material 9, 10), preventing a clear identification of their evolutionary relatedness to the mobile sequences from other organisms.

Phylogenomic analyses were performed on two previously determined sets of highly conserved protein coding genes, respectively 24 (Lang et al. 2013) and 120 genes (Parks et al. 2018), with different but equally representative taxon selections (respectively gene set 1: 23 taxa and gene set 2: 100 taxa). The retrieved results were consistent, and confirmed those of 16S rRNA gene analysis, in particular the affiliation of *Deianiraea* to *Rickettsiales sensu stricto* and the monophyly of the group composed by *Anaplasmataceae*, *Midichloriaceae* and *Deianiraea* (Figure 4; Supplementary material 11, 12). In addition, unlike the tritomy shown in the 16S phylogeny, these phylogenomic analyses indicate a sister relationship of *Deianiraea* to *Anaplasmataceae*. Moreover, none of the analysis retrieved a sister relationship of *Holosporales* respect to *Rickettsiales sensu stricto* and, in the phylogenomic tree of gene set 2 (including available *Holosporales*; Supplementary material 12), *Holosporales* did not appear monophyletic, in agreement with recent phylogenomic trees (Parks et al. 2018).

**Figure 4.**
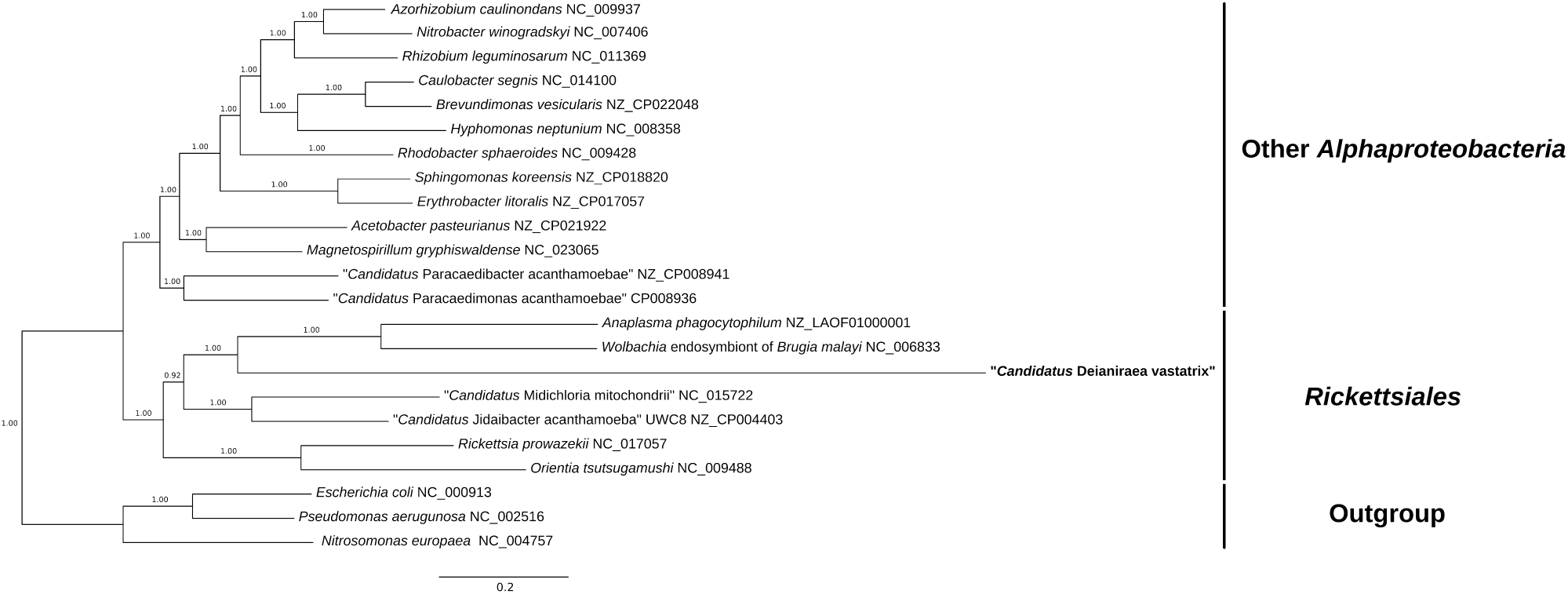
Bayesian phylogenomic tree of representative *Rickettsiales* and other *Alphaproteobacteria.* Inference was based on the concatenated protein alignment (4,038 characters) from 24 highly conserved orthologs (Gene set 1; Lang et al. 2013) with the LG+I+G evolutionary model. Posterior probabilities (after 1,000,000 generations) of each internal tree nodes are shown on branches (The maximum likelihood tree inferred on the same dataset is shown in Supplementary material 11). The newly characterized *Deianiraea* putative parasite of *P. primaurelia* CyL4-1 is evidenced in bold. The *Rickettsiales* (including representatives of each of the four families *Rickettsiaceae*, *Anaplasmataceae*, *Midichloriaceae,* and *Deianiraeaceae*), other *Alphaproteobacteria*, and the outgroup are evidenced on the right side. The scale bar stands for estimated 0.2 sequence divergence.

### Predicted *Deianiraea* metabolic features

The *Deianiraea* ORFs were assigned to a total of 749 distinct COGs (Supplementary material 13). Most of the predicted metabolic and cellular features, including number and proportion of different functional categories, are in compliance with other *Rickettsiales* (Supplementary material 14, 15). Slight differences can be found in stress response, namely the presence in *Deianiraea* of two peculiar azoreductases.

For what concerns DNA repair, the *Deianiraea* genome encodes the major systems, such as mismatch repair, nucleotide excision repair and homologous recombination after single-strand break, as well as DNA alkylation repair (AlkB and AlkD proteins).

In terms of carbohydrate and energetic metabolism, *Deianiraea* is predicted to perform complete gluconeogenesis and the non-oxidative pentose-phosphate pathway (except transaldolase), but key glycolytic enzymes are missing. The gene sets for pyruvate dehydrogenase, Krebs cycle and oxidative phosphorylation are present, including NADH dehydrogenase, succinate dehydrogenase, cytochrome *bd* terminal oxidase and ATP synthase, but, differently from other *Rickettsiales*, cytochrome *c* and its oxidase and reductase are missing. A *tlc* ATP/ADP translocase, homolog to those of other *Rickettsiales*, was identified as well.

*Deianiraea* is also equipped with biosynthetic pathways for the main structural components of cell inner membrane, cell wall and outer membrane, namely phospholipids, isoprenoids (via methylerythritol phosphate pathway), peptidoglycan, and lipopolysaccharide (LPS). These traits are shared with many other *Rickettsiales,* with the significant exception of *Anaplasmataceae* and *Orientia,* which lack LPS and, partially or completely, peptidoglycan. No capsule synthesis is present in *Deianiraea*, as compared to “*Ca*. Jidaibacter acanthamoeba” (Wang & Wu 2014; Schulz et al. 2016).

In compliance with other *Rickettsiales*, there is an overall paucity of other biosynthetic pathways in *Deianiraea.* In particular, the potential for nucleotide synthesis is scarce, and this bacterium likely relies on interconversion. Only some cofactors can be produced, namely biotin, hem, ubiquinone, folate (the synthetic pathway includes a two-component aminodeoxychorismate synthase) and Fe-S clusters. The gene sets for several different membrane transporters, including ABC and MFS types, were found. As in other *Rickettsiales*, their activity may complement the metabolic deficiencies of *Deianiraea*. Contrarily to other recently sequenced *Rickettsiales* genomes (Sassera et al. 2011; Martijn et al. 2015), no flagellar or chemotaxis genes were found.

A total of 84 COGs (~11.2%) resulted exclusive of *Deianiraea*, and absent in the other analyzed *Rickettsiales* (Supplementary material 16). Among those COGs, several genes involved in the biosynthesis of multiple amino acids and in cell adhesion were found, representing the prominent peculiarities of the novel genome. These proteins were thus analyzed in detail. Notably, *Deianiraea* is also the only *Rickettsiales* member possessing a bacterial cytoskeletal component, namely a bactofilin family protein possibly involved in asymmetrical cell division and cell polarity (Kühn et al. 2010), a morphological feature that is indeed shown by *Deianiraea* (see Figure 2).

### Amino acid metabolism

*Deianiraea* has the predicted capability to synthesize the vast majority (16 out of 20) of proteinogenic amino acids, a striking difference with respect to all known *Rickettsiales* (Figure 5; Supplementary materials 17, 18, 19). Moreover, most importantly, the biosynthesis of eight of those amino acids is a unique competence of *Deianiraea* among *Rickettsiales*, raising the total number of amino acids that can be synthesized by at least one *Rickettsiales* member from 10 to 18. *Deianiraea* is by far the richest *Rickettsiales* in term of amino acids biosynthesis, with a total of 56 genes devoted to amino acid production, while no other member of the order outreaches 30 genes (Figure 5; Sassera et al. 2011; Schulz et al. 2016; Dunning-Hotopp et al. 2006; Andersson et al. 1998; Martijn et al. 2015).

**Figure 5.**
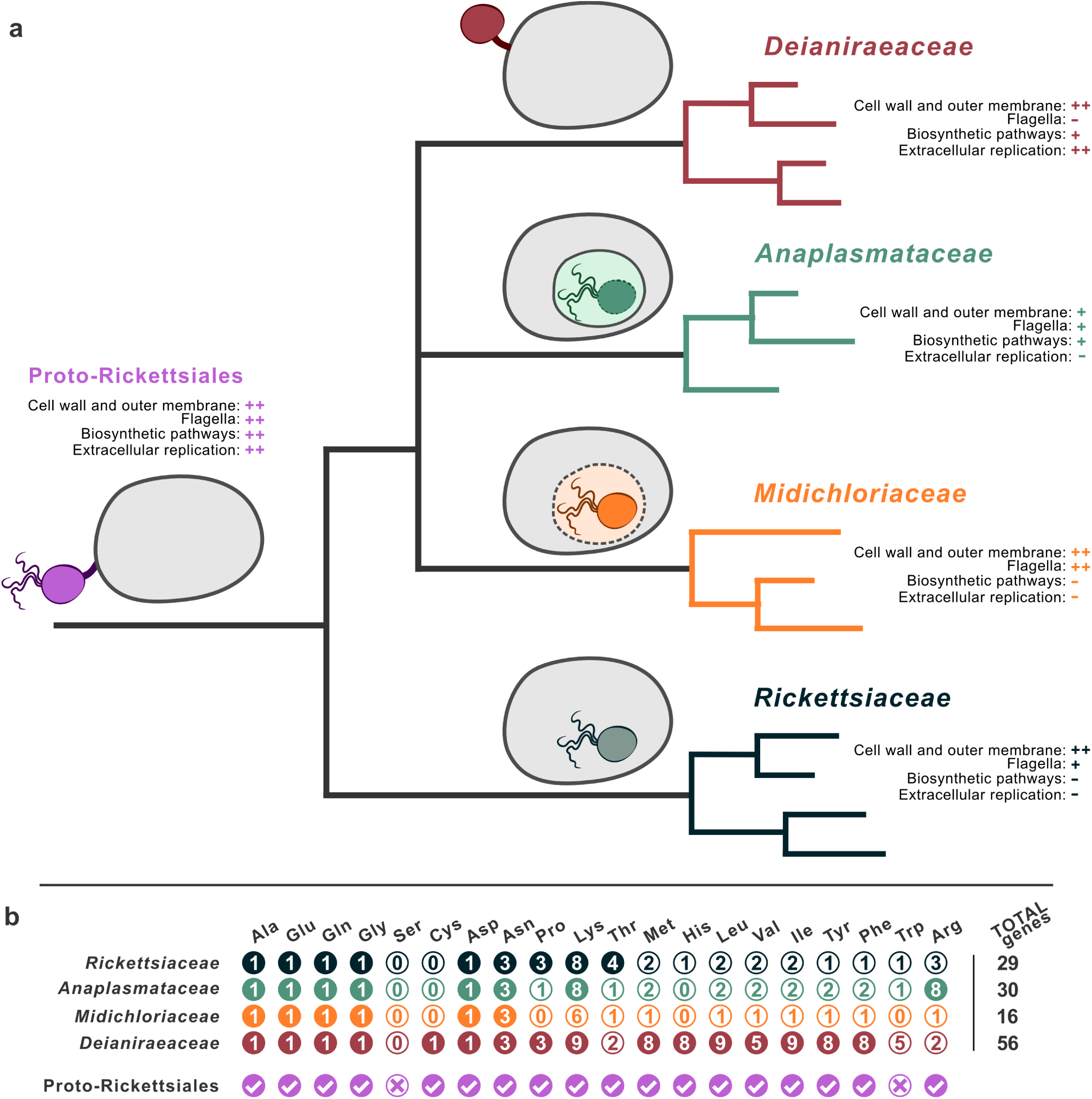
Reconstrution of ancestral *Rickettsiales* features according to the “intracellularity late” hypothesis. **a,** An idealized evolutionary tree of the four *Rickettsiales* families, is shown. The drawings and notes summarize selected features of each family on the right, and the inferred features of the ancestor at the root of the tree. In the drawings, flagella are shown if present in at least one member of the family, while vacuolar membrane is shown as a solid line if present for all members of the family (*Anaplasmataceae*), or a dashed line if present only for some members (*Midichloriaceae*). “Cell wall and outer membrane” and “Biosynthetic pathways” indicate the degree of completeness of, respectively, peptidoglycan and lipopolysaccharide biosynthesis or the synthesis of amino acids, cofactors, and nucleotides: “++” stands for complete in most known representatives, “+” for (at most) rich in most known representatives, “-” for (at most) scarce in most known representatives. Solid and dashed bacterial outlines in the drawing correspond to different degrees of “Cell wall and outer membrane”. “Flagella” and “Extracellular replication” indicate, respectively, the presence of flagella/flagellar genes or of areplication extracellular respect to the host cells (alike *Deianiraea*): “++” stands for present in most known representatives, “+” for present in some known representatives, “-” for absent in all known representatives. **b,** Summary of the amino acids biosynthetic abilities in the *Rickettsiales* lineages. Columns stand for single amino acids, rows for organisms or families. Filled circles indicate that the corresponding biosynthetic pathway is present in the respective organism or family (presence in a single family member is the sufficient condition). The number in each circle indicate the total number of distinct ortholog genes involved in the pathway found in the respective organism or family (Supplementary Material 18). The pathway presence in the Proto-Rickettsiales ancestor was inferred based on the presence in at least one current *Rickettsiales*. Total numbers of genes per organism or family are indicated on the right, and differ from the sum of the rows due to the presence of genes shared by multiple pathways (Supplementary Material 18).

### Interaction with *Paramecium*

*Deianiraea* is equipped with several secretion systems, which may mediate the interaction with *Paramecium*. These include most of the *Rickettsiales* typical set (Gillespie et al. 2015), namely Sec and Tat translocons, and *rvh* type IV secretion system. In compliance with the canonical composition in *Rickettsiales*, *Deianiraea* type IV secretion system includes single copy (VirB2, VirB3, VirB10, VirB11, VirD4), duplicated (VirB4, VirB8, VirB9), and multi-copy (VirB6: 3 copies) subunits, and is devoid of the extracellular subunit VirB5 (Gillespie et al. 2016). Differently from other *Rickettsiales*, no VirB7 homolog was found, possibly representing a false negative due to the short length of this gene and its high sequence divergence (Gillespie et al. 2009).

On the other side, components of secretion systems widely present in other *Rickettsiales*, namely the Apr/Hly type I secretion system proteins and Sca-like type V system autotransporter proteins (Gillespie et al. 2015), are absent.

The components of two distinctive secretion and interaction systems were found. Indeed, the core gene set for type II secretion system (Korotkov et al. 2012), including three pseudopilins, was identified, a structure having an equivalent putatively complete counterpart among *Rickettsiales* only in “*Ca*. Jidaibacter”. In addition, an unusual two-partner type V secretion system was detected. Two ORFs (595 and 1997 aa), arranged in a putative operon, were recognized as secreted effectors (COG3210), and include one or more hemagglutinin repeats, respectively (Supplementary material 13). Proteins of this family have a role in cell adhesion and toxicity towards eukaryotes or other bacteria, and frequently occur in operons, where shorter gene products may display a regulative activity (Guérin et al. 2017). Despite the recognizable structural motifs, the two proteins share extremely limited (<32%) sequence identities with homologs from other bacteria, preventing to clearly identify their evolutionarily closest relatives and thus more precisely infer their function. In close proximity (but reverse oriented), the gene for the corresponding outer-membrane transporter was found (COG2831, Supplementary material 16). No homologs of adhesins or invasins typical of other *Rickettsiales* were found.

Fifty-three ORFs were predicted to possess a signal peptide and no transmembrane helix, a group that may include secreted effectors (Supplementary material 20). Interestingly, some of these ORFs present eukaryotic-like domains potentially involved in interaction with *Paramecium*, such as ankyrin and tetratricopeptide repeats. Additionally, three MORN repeat proteins (COG4642) were identified in the *Deianiraea-*exclusive sets (Supplementary material 16). This domain is also potentially involved in host cell adhesion and invasion (McGuire et al. 2014).

### Horizontal gene transfer evaluation

The possibility that the 98 proteins assigned to the 84 *Deianiraea*-exclusive COGs were the result of horizontal gene transfer (HGT) was investigated by inspecting sequence identity of blastp hits across multiple taxonomic ranges, and, in selected cases, phylogenetic analyses (Supplementary material 16, 22). In 72 cases, the high sequence divergence allowed to exclude a recent HGT event from a known donor. For other 6 genes, phylogenetic analyses were applied, and did not produce supported topologies, preventing to robustly trace the recent evolutionary history of those *Deianiraea* genes (Supplementary material 21). On the other side, a some of the genes (21) were found to actually have putative homologs in some *Rickettsiales* (other than those employed in the COG classification).

The products of the remaining genes assigned to *Deianiraea-*exclusive COGs are involved in amino acids biosynthetic pathways. Considering that such pathways were the most distinctive genomic trait identified in *Deianiraea* with respect to other *Rickettsiales*, additional and more detailed analyses were employed on all the genes involved. These analyses included phylogenies on concatenated protein sequences, statistical tests based on GC content and CAI (codon adaptation index), and careful comparative analyses of the inferred pathways (Supplementary material 17-19). Overall, because of high sequence variation with respect to any other organism and compositional consistence with respect to the other *Deianiraea* ORFs, no clear recent HGT was evidenced (Supplementary material 19), further supporting the inference that these genes could have been vertically inherited from the *Rickettsiales* ancestor.

## Discussion

The order *Rickettsiales* encompasses a wide range of bacteria that have attracted the interest of numerous researchers through the years, due to their importance as human pathogens, mutualist symbionts, parasites and manipulators of the host reproduction (Werren et al. 2008; Weinert et al. 2009; Perlman et al. 2006; Hosokawa et al. 2010; Taylor et al. 2005). In this plethora of behaviors and lifestyles, the intracellular localization has always been marked as a constant. Indeed, all *Rickettsiales* show obligatory intracellular associations with host cells, which are necessary for their cell division. Parsimoniously, this trait has always been considered as inherited from the ancestor of the order (Darby et al. 2007; Weinert et al. 2009; Sassera et al. 2011; Schulz et al. 2016).

Herein, we present *Deianiraea vastatrix*, the first *Rickettsiales* bacterium with a completely extracellular lifestyle, albeit associated with a unicellular eukaryote. The bacterium has been observed in an extracellular and putatively parasitic interaction with the single-celled eukaryote *P. primaurelia*, showing the ability to attach to its external surface, but never entering the cell (Figure 1; Figure 2). *Deianiraea* replicates in this extracellular location, covering the *Paramecium* cells, being the apparent cause of observed morphological alterations and lethal effects, with variable degree of virulence depending on the genotype of the ciliate. It may be hypothesized that the massive loss of cilia could be the event triggering the degeneration of *Paramecium* cells. Indeed, loss of cilia caused by heavy metals has been shown to cause mortality in ciliates (Larsen and Nilsson 1983), and the effect caused by *Deianiraea* could be analogous. This kind of interaction between an extracellular bacterium and a unicellular eukaryote is reminiscent of the case of the phylogenetically unrelated *Vampirovibrio chlorellavorus* (*Melainabacteria*) and *Chlorella* algae (Soo et al. 2014). However, *Vampirovibrio* has a much larger genome (~3 Mb, including two plasmids) and a consistently wider metabolic repertoire, including lytic enzymes putatively involved in digestion of host cells. Interestingly, it can synthesize almost the same amino acids as *Deianiraea* (15 in total: comparatively able to synthesize tryptophan and unable to produce histidine or phenylalanine).

The molecular bases of the interaction between *Deianiraea* and its host are not disclosed, but several genes mined in the genome of this bacterium provide hints on the possible mechanisms used to attack the ciliate, namely the presence of peculiar secretion systems and adhesion molecules. In particular, a distinctive two-partner type V secretion system likely enables the bacterium to release effectors bearing hemagglutinin repeats, with potential roles both in cell adhesion and toxicity (Guérin et al. 2017). Moreover, considering that *Deianiraea* possesses a type II secretion system, rare in *Rickettsiales* (only “*Ca*. Jidaibacter presents an equivalent putatively functional set; Schulz et al. 2016), and that components of this system are homologous of pili (Mattick 2002), a role of this protein complex in adhesion or motility of *Deianiraea* may also be hypothesized. Additionally, while the overall metabolic repertoire of *Deianiraea* is limited, it probably acquires the necessary small precursor molecules directly from the *Paramecium* host, exploiting a relatively wide set of membrane transporters, including the ATP/ATP translocase, as in other *Rickettsiales.* Indeed, it can be speculated that such molecules could be made available for the bacteria in the extracellular proximity of *Paramecium* either by a *Deianiraea*-triggered mechanism, or simply released after death of infected host cells.

In synthesis, *Deianiraea* presents a unique extracellular lifestyle, distinct from all other *Rickettsiales*. This novel finding raises the question whether *Deianiraea* has reverted to extracellularity from an obligate intracellular ancestor or extracellularity was the ancestral state of the last common ancestor of the order.

As evidenced in our phylogenetic and phylogenomic analysis (Figure 3; Figure 4), *Deianiraea* is a member of a new family level clade of *Rickettsiales* (herein named *Deianiraeaceae*), which is not the earliest diverging clade, but is clearly phylogenetically derived. Screening on the IMNGS 16S rRNA amplicon database showed that the diversity of *Deianiraeaceae* is comparable to the other three families of *Rickettsiales*. No information is available on other *Deianiraeaceae* bacteria, however in some cases their origin, namely the skin of aquatic animals or humans (Kong et al. 2012; Fraune and Bosch 2007; Fraune et al. 2010; Bauermeister et al. 2012), allows to wonder whether they may be extracellular as well.

Moreover, genomic analyses showed that *Deianiraea* displays genes encoding for the biosynthetic pathways of 16 amino acids, a repertoire much larger than that of any other known *Rickettsiales* (Figure 5; Sassera et al. 2011; Schulz et al. 2016; Dunning-Hotopp et al. 2006; Andersson et al. 1998). Although available data do not offer a conclusive explanation on the functional relevance of such wider metabolic capabilities, these can be considered consistent with the unique extracellular lifestyle of *Deianiraea.* Due to the absence of the *Deianiraea*-exclusive amino acids biosynthesis gene sets in all other sequenced *Rickettsiales*, to the low level of identity with available orthologs and to the consequently limited phylogenetic signal, it is difficult to evaluate whether they are ancestral or the result of an ancient HGT event. In the future, the sequencing of additional phylogenetically related genomes may allow to obtain a more complete picture. At present, considering also the homogeneity of GC content and codon usage to the rest of the genome, we can assert that there is no evidence of recent HGT for any of the pathways analyzed (Supplementary material 19). This scenario is consistent with the expectations in the case they were indeed inherited vertically from the last common ancestor of *Rickettsiales* (Proto-Rickettsiales), which, according to such reconstruction, should have been more biosynthetically capable than previously thought (up to 18 amino acids) (Figure 5). On the other hand, in the context of potential HGT events involving *Deianiraea* genes in general, a role of mobile elements cannot be excluded, although the impossibility to trace the recent evolutionary history of these elements (Supplementary material 9, 10) prevents to clearly evaluate and discern such role.

Considering literature data, we can conclude that all known *Rickettsiales*, except *Deianiraea,* are intracellular, however most of them are able to perform horizontal transmission and host species switch through a temporary extracellular state. The process and molecular recognition machinery used to enter host cells, as well as the consequent intracellular behavior and location, are not conserved among the *Rickettsiales* lineages, and might differ even within the same family (Walker and Ismail 2008; Ge and Rikihisa 2011; Rikihisa 2010). As a matter of fact, *Rickettsiaceae* enter host cells by inducing phagocytosis, and typically escape the phagosomal membrane, residing freely in the host cytoplasm and exploiting host cytoskeleton for movement (Cardwell and Martinez 2011; Gouin et al. 2004; Walker and Ismail 2008; Renesto et al. 2009; Ge and Rikihisa 2011). *Anaplasmataceae* hijack the phagocytic process, modifying the host vacuole for their survival and proliferation, thus exploiting the intra-vacuolar niche (Rikihisa 2010; Kumar et al. 2013; Kahlon et al. 2013; Seidman et al. 2014). *Midichloriaceae* are more diverse, encompassing both bacteria enclosed in host-derived membranes and others dwelling free in the cytoplasm (Vannini et al. 2010; Sassera et al. 2011; Schulz et al. 2016). Despite direct evidence of their host transfer abilities (Schulz et al. 2016; Senra et al. 2016), the cell entering mechanisms of *Midichloriaceae* are poorly understood.

Taken together, the discovery of an extracellular *Rickettsiales* and the above detailed variability of *Rickettsiales*, both in general and in terms of interaction with hosts, allow to propose an alternative hypothesis on the evolution of the order. The classical hypothesis, which we may call “intracellularity early”, assumes that the common *Rickettsiales* ancestor was an obligate intracellular bacterium, considering that all known *Rickettsiales*, up to now, shared that trait. Here we propose the novel “intracellularity late” hypothesis, largely based on the new and unprecedented features observed in *Deianiraea*, but also on the diversity of “intracellularity” among all other *Rickettsiales*. Indeed, in our opinion, the traits that are specific to single *Rickettsiales* families, rather than different shades of a common mechanism, can be seen as indicative of multiple independent instances of evolution of host cell entrance and interaction mechanisms, leading to parallel and independent evolution of analogous intracellular lifestyles within *Rickettsiales*. However, while the single reversal of *Deianiraea* to extracellularity (necessary for the “intracellularity early” hypothesis) could be seen as more parsimonious, it is difficult to evaluate how many and which changes and gains-of-function would have been necessary for such a complex and, to our knowledge, unprecedented evolutionary process.

Following the “intracellularity late” hypothesis, we inferred the main features of the last common ancestor of *Rickettsiales* (Proto-Rickettsiales), based both on previous knowledge and on novel findings (Figure 5). According to this interpretation, the discovery of *Deianiraea* allows to interpret pre-existing observations on *Rickettsiales* from a different perspective, suggesting that the ancestor may have been extracellular, though probably displaying a pre-adaptation to the interaction with eukaryotic cells such as type IV secretion systems (Gillespie et al. 2009) or ATP/ADP translocases (Schmitz-Esser et al. 2004). In this scenario, such characteristics were instrumental in the parallel evolution of multiple strictly intracellular lineages within the order. Considering that most *Rickettsiales* (including *Deianiraea*) are found in aquatic environments (Castelli et al. 2016) (Figure 5), the Proto-Rickettsiales was probably an aquatic bacterium as well (Vannini et al. 2005; Ogata et al. 2006; Weinert et al. 2009; Driscoll et al. 2013; Schrallhammer et al. 2013; Kang et al. 2014). Combining multiple features present in different *Rickettsiales* lineages, it is possible to infer that such Proto-Rickettsiales bacterium probably had a rather high metabolic and functional complexity, consistent with the extracellular and likely eukaryotic-associated lifestyle (Figure 5). In details, in terms of metabolic capabilities, it could have been equipped with the core biosynthetic pathways, in particular for amino acids (present work), but also for cofactors and nucleotides (as in multiple current *Anaplasmataceae*, Dunning Hotopp et al. 2006), and was possibly capable to survive in low-oxygen conditions (Sassera et al. 2011). From the morpho-functional side, it could have been provided with a robust structure made by functional peptidoglycan and LPS (Driscoll et al. 2017; Sassera et al. 2011; Schulz et al. 2016, present study), could synthesize capsule (Schulz et al. 2016), possessed flagella (Sassera et al. 2011; Vannini et al. 2014; Schulz et al. 2016; Kwan and Schmidt 2013; Martijn et al. 2015) and pili (present work; Schulz et al. 2016), and was able to perform chemotaxis (Martijn et al. 2015). During the course of evolution, the association of its descendants to eukaryotic cells would have then become tighter and tighter, with a reduction in genome size and complexity, finally leading to the parallel depletion of most metabolic and functional features, including, in most known lineages, the capability to reproduce extracellularly, and a consequent adaptation to the obligate intracellular state. Therefore, the diversity in host range, mechanisms of interaction, and functional and metabolic features of current *Rickettsiales*, as well as their generally conserved ability to colonize new hosts, could be seen as a reflection of the much higher versatility of their ancestor. Such “intracellularity late” scenario would have additional consequences on the inference of the even more ancient evolution of mitochondria from *Alphaproteobacteria*. Indeed, there is an open debate on whether mitochondria and *Rickettsiales* are sister groups (Andersson et al. 1998; Fitzpatrick et al. 2006; Wang and Wu 2015), or they are not phylogenetically related (Esser et al. 2004; Thiergart et al. 2012; Martijn et al. 2018). Assuming that the Proto-Rickettsiales was in fact an extracellular bacterium, even in case mitochondria and *Rickettsiales* truly shared a common ancestor, it would become evident that such ancestor should have been extracellular as well. Consequently, their intracellular lifestyles would have been acquired independently, much similarly to what would have happened according to the alternative hypotheses not implying a close phylogenetic relatedness of mitochondria and *Rickettsiales* (Martijn et al. 2018). The main difference would be that a hypothetical common ancestor of mitochondria and *Rickettsiales* could be envisioned as a bacterium with a degree of metabolic independence, and possibly equipped with the machinery to interact with other microorganisms. Such “interaction-prone” ancestor could explain the capability of evolving intracellularity multiple times, within *Rickettsiales* and, possibly, by the ancestor of mitochondria.

In recent years, an increasing amount of research was produced on neglected members of the order *Rickettsiales* (e.g. Wang & Wu 2014; Martijn et al. 2015; Schulz et al. 2016; Szokoli et al. 2016b; Castelli et al. 2018; Floriano et al. 2018; Yurchenko et al. 2018), which, contrarily to those thoroughly studied in the past, are not of direct medical or veterinary concern. Such investigations represented a major contribution to understand the phylogenetic and genomic diversity of *Rickettsiales*, and are gradually reshaping our interpretation of their evolution. In this framework, the discovery of *Deianiraea* has the potential to represent a tipping point for the evolutionary views on *Rickettsiales*, providing the first supporting evidence for the novel idea that their ancestor could have been an extracellular bacterium, and that, accordingly, obligate intracellularity in this order would have evolved multiple times in parallel and independently. Obviously, with the available data it is still premature to conclude which hypothesis is preferable between the “intracellularity early” and “intracellularity late” in *Rickettsiales*. In this regard, future research on other under-investigated lineages of *Rickettsiales*, in particular *Deianiraeaceae*, may definitely lead to a more complete understanding of the processes that have led to the evolution and diversification of extant *Rickettsiales*, and may thus support one of the two hypotheses.

### Description of “*Candidatus* Deianiraea vastatrix” gen. nov. sp. nov

*“Candidatus* De.i.a.ni.rae’a va.sta’trix” (De.ia.ni.rae’a; N.L. fem. n., dedicated to Deianira, in reference of the Greek myth in which Heracles was killed by his wife Deianira, who covered him with a tunic poisoned by the blood of the centaur Nessus; N.L. adj. from *vastare*, to devastate, to destroy, *vastatrix,* the destroyer).

Bacterium found in an extracellular association with the ciliate *Paramecium primaurelia* CyL4-1. Slightly curved rod-like shape (1.6 to 1.7 μm by 0.1 to 0.2 μm in size), frequently displaying a narrowed side ending up with a sharp apical tip, located in close proximity of or even direct contact with *P. primaurelia* cell membrane from the outside. Electron-dense cytoplasm. Devoid of flagella. Basis of assignment: SSU rRNA gene sequence (accession number: MH197138) and positive match with the species-specific FISH oligonucleotide probes Deia_416 (5′-GAGTTTTACAATCTTTCG-3′) and Deia_538 (5′-AGTAACGCTTGGACTCCA-3′). The complete genome sequence of the bacterium was deposited under the accession CP028925.

### Description of “*Candidatus* Deianiraeaceae” fam. nov

*“Candidatus* Deianiraeaceae” (De.i.a.ni.rae.a′ce.ae, N.L. fem. n. “*Candidatus* Deianiraea” type genus of the family; suff. -*aceae* ending to denote a family; N.L. fem. pl. n. “*Candidatus* Deianiraeaceae” the family of genus “*Candidatus* Deianiraea”)

The family “*Candidatus* Deianiraeaceae” is defined based on phylogenetic analysis of 16S rRNA gene sequences of the type genus and uncultured representatives from various origins; several sequences derive from freshwater environments, and were frequently retrieved in association with eukaryotic organisms. The family belongs to the order *Rickettsiales*, and currently contains one genus, “*Candidatus* Deianiraea”.

### Emended description of the order *Rickettsiales* Gieszczykiewicz (1939) Dumler, Barbet, Bekker, Dasch, Palmer, Ray, Rikihisa and Rurangirwa 2001

*Rickettsiales* (Rick.ett.si.a’les. N.L. fem. n. *Rickettsia* type genus of the order; -*ales* ending to denote order; N.L. fem. pl. n. *Rickettsiales* the *Rickettsia* order)

The description is identical to the one provided by Dumler & Walker (2015), with the following amendments (**evidenced in bold**):

“Rod-shaped, coccoid or irregularly shaped bacteria with typical Gram-negative cell walls. **Sometimes harbouring flagella. In most cases**, multiply only inside host cells, **but at least one exception is known”**

and

“The bacteria are parasitic forms associated with host cells of the mononuclear phagocyte system, the hematopoietic system, or the vascular endothelium of vertebrates; with various organs and tissues of helminths; with tissues of arthropods, which may act as vectors or primary hosts; **or with unicellular eukaryotes, either intracellularly or extracellularly”**

## Materials and Methods

### *Paramecium* strain origin and maintenance

The monoclonal *Paramecium* strain CyL4-1 was isolated on 29^th^ September 2014 from a water sample (9‰ salinity) collected from a waste water stream in Larnaca, Cyprus (34.91° N, 33.60° E). No specific permission was required for sampling, since the location was not a protected area, and ciliate and bacterial species are not considered endangered/protected.

CyL4-1 was initially maintained in a lettuce infusion inoculated with *Enterobacter aerogenes* (initially 5‰ salinity, then adapted to 0‰). In order to maintain for an extended period of time in the laboratory the *Paramecium* strain harboring *Deianiraea* bacteria, a transfer procedure was performed every 2-3 days. In detail, single viable cells of *Paramecium* were picked up and transferred into a Petri dish containing 250 μl of fresh lettuce infusion inoculated with *E. aerogenes* and supplemented with 0.8 μg/ml β-sitosterol (Merck, Darmstadt, Germany).

### Live cell observations

For living cell observations paramecia were analyzed using differential interference contrast (DIC) with a Leica 6000 microscope (Leica Microsystems, GmbH, Wetzlar, Germany) equipped with a digital camera DFC 500.

### Transmission Electron Microscopy

*Paramecium* cells were fixed with a phosphate fixative, containing 2.5% glutaraldehyde, 1.6% paraformaldehyde in 0.04 M phosphate buffer (pH 7.2-7.4), for 1.5 h, washed with the same buffer containing 12.5% sucrose (two steps, 30 min each), and postfixed with 1.5-2% osmium tetroxide for 1 h. Then, the material was processed as described earlier (Szokoli et al. 2016b). After dehydration, the cells were embedded in Epoxy embedding medium (Fluka, BioChemika) according to the manufacturer’s protocol. The blocks were sectioned with a Leica EM UC6 ultracut, stained with 1% aqueous uranyl acetate and 1% aqueous lead citrate and examined with a Jeol JM 1400 (Jeol, Ltd., Tokyo, Japan) electron microscope at a voltage of 90 kV.

### Atomic Force Microscopy

Living cells were placed on a cover slip and air dried. The surface topography was registered by means of NTEGRA AURA microscope in a semi-contact mode.

### *Deianiraea* shift assays

In order to check the ability of *Deianiraea* to colonize the cell surface of other *Paramecium* strains and affect them, cells of the strain CyL4-1, preliminary confirmed to be abnormally-shaped, unable to move, and densely covered by *Deianiraea* bacteria, were placed in Petri dishes containing 250 μl of lettuce infusion inoculated with food bacteria, ~50 *Paramecium* cells for each dish. The *Deianiraea*-free CyL4-1* sub-replicate of the original strain plus six additional naïve strains were then selected (Supplementary material 3), and each added in duplicate to these Petri dishes, 25-30 cells per dish. As a control, the target cells were transferred in dishes containing only lettuce infusion inoculated with food bacteria. Cells were fed once a week with 50 μl of fresh medium inoculated with food bacteria. The dishes were checked every 4 days for the presence of living motile cells and for the signs of the developing colonization by *Deianiraea* for a period of 40 days.

### Molecular characterization of *Paramecium* and *Deianiraea*

About 80-100 CyL4-1 cells were individually picked up, washed through several passages in sterile water and fixed with 70% ethanol. DNA was extracted with the Nucleospin Plant II kit (Macherey-Nagel), following the protocol for mycelium. All PCR reactions were performed with the TaKaRa ExTaq (TaKaRa Bio Inc), as previously described (Lanzoni et al. 2016). For the molecular characterization of *Paramecium* 18S rRNA gene, ITS and COI gene were amplified and directly sequenced (Lanzoni et al. 2016) (Supplementary material 22).

Characterization of *Deianiraea* was performed by PCR amplification of the almost complete 16S rRNA gene with universal bacterial primers (modified from Lane (Lane 1991)), as previously described (Boscaro et al. 2013), followed by sequencing with internal primers (Vannini et al. 2004).

### Fluorescence *in situ* hybridization

*Paramecium* cells were fixed in a depression slide with 4% paraformaldehyde in PBS for 30 min, transferred onto a SuperFrostPlus slide (Menzel-Glaser, Germany), and washed in PBS. The excess liquid was removed, and the cells were postfixed with ice-cold 70% methanol. The slides were washed in PBS, and a drop of hybridization buffer containing 10 ng/mL of each oligonucleotide probe employed and 15% formamide was added. The slides were covered with parafilm and placed in a wet chamber. The hybridization was performed for 2 h at 46°C, followed by two subsequent incubations with a washing buffer at the same formamide concentration, 30 min each, at 48°C (Manz et al. 1992). The hybridization buffer contained 10 ng/mL of each of the oligonucleotide probes and 15% formamide. Specific probes for *Deianiraea* were newly designed, and their specificity was assessed by search on the RDP (Ribosomal Database Project) (Cole et al. 2014) and on SILVA (Quast et al. 2013) (Supplementary material 23). Finally, the slides were mounted in Mowiol (Mowiol 4.88, Calbiochem) diluted in glycerol containing *p*-phenylenediamine (PPD) and DAPI (4′,6-diamidino-2-phenylindole) according to manufacturer protocol. The slides were analyzed with a Leica TCS SPE2 Confocal Laser Scanning Microscope. The images were further processed with Fiji-win32 open access software.

### 16S rRNA phylogenetic analysis

The partial 16S rRNA gene sequence of *Deianiraea* was aligned on the SSU Ref NR 99 Silva database (Quast et al. 2013) (together with the addition of selected closely related sequences retrieved on NCBI nucleotide) with the ARB software package (Westram et al. 2011), employing the automatic aligner followed by manual inspection to optimize base-pairing in the predicted rRNA structure. After preliminary phylogenetic analyses (data not shown) a total of 33 other *Rickettsiales sensu stricto* (Szokoli et al. 2016a) sequences were selected, plus 10 other *Alphaproteobacteria* as outgroup. The alignment was trimmed at both ends to the length of the shortest sequence, resulting in 1,398 characters (Supplementary material 24). Maximum likelihood (ML) and Bayesian inference phylogenetic analysis were performed as previously described (Sabaneyeva et al. 2018), employing respectively PhyML (Guindon and Gascuel 2003) and MrBayes (Ronquist et al. 2012), after selection of the best substitution model with jModelTest (Darriba et al. 2012). 16S rRNA gene identity values were calculated on the alignment employed for phylogeny.

### 16S rRNA gene amplicon database screening

With ARB, the whole set of sequences belonging to each of the four *Rickettsiales* families *sensu stricto* were selected from the SSURef NR99 128 Silva database. Chimeras were filtered out with online DECIPHER (Wright et al. 2012) and manual inspection of blastn results, and remaining sequences were grouped in clusters with ≥ 93% identity. The final representative set of *Rickettsiales* included a member for each cluster (the sequence with the lower average divergence from the same cluster) (Supplementary material 25). Thus, the sequences from this set were queried on the IMNGS (integrated microbial NGS) platform (Lagkouvardos et al. 2016), selecting hits with 90% minimal identity and 200 bp minimal length. For each family, the retrieved hits were merged, and ambiguous sequences (hit by members of two or more families) were removed. Operational taxonomic units (OTUs) for each family were retrieved with UCLUST (Edgar 2010) with 97% identity. Each OTU was then assigned to its most prevalent environmental origin according to IMNGS categories.

### Genome sequencing and assembly

Full details on the sequencing and assembly procedure can be found in the Supplementary material (Supplementary material 26, 27, 28, 29, 30) Approximately 100 cells of *Paramecium* covered by *Deianiraea* were individually picked up from the cultures, washed through several passages in sterile water and fixed in 70% ethanol. Thus, DNA was extracted with Nucleospin Plant II kit, using the protocol for mycelium. In order to reach sufficient DNA quantity for sequencing, the extract was subjected to whole-genome amplification (WGA) with the REPLI-g Single Cell Kit (Qiagen), following the instructions for DNA extracts. The WGA product was processed through a Nextera XT library, and sequenced on an Illumina HiSeq X machine by Admera Health LC (South Plainfield, NJ, USA), producing 37,100,964 pairs of 150 bp reads.

After quality check with FastQC (Andrews 2010), the reads were preliminary assembled using SPAdes 3.6 (Bankevich et al. 2012). The blobology pipeline was then applied (Kumar et al. 2013), in order to select only those contigs putatively belonging to *Deianiraea* from *Paramecium* and other free-living bacteria, namely, contigs with coverage ≥ 100x and best megablast hit with *Bacteria* or no hit were selected. This set of contigs was manually checked and revised. Thus, the reads mapping on the final selection of contigs (Langmead and Salzberg 2012) were reassembled separately, obtaining a total of 12 contigs. Assembly graph was visualized with Bandage (Wick et al. 2015), in order to detect putative connections among contigs. Consequently, regular and two-step walking PCR (Pilhofer et al. 2007) assays were designed and applied, allowing to join all the contigs (Supplementary material 30) into a single circular chromosome (sized 1,205,153 bp).

### Genomic analyses

The *Deianiraea* genome was annotated with predicted CDSs and non-coding RNAs by using Prokka (Seamann 2014), and functional annotation was manually curated. In details, all the predicted CDSs were queried by blastp on NCBI nr proteins, Swissprot, and *Rickettsiales*-only sequences from nr. For each CDS, the results were carefully examined manually, taking into account also protein lengths, and employing NCBI conserved domain searches (Marchler-Bauer et al. 2015). The Prokka annotation was thus manually modified accordingly. Predicted secreted proteins and transmembrane domains were predicted with SignalP (Emanuelsson et al. 2007) and TMHMM (Krogh et al. 2001).

For gene number and coding density comparisons, annotated genomes of all *Rickettsiales* bacteria (manually excluding metagenomic assemblies) were downloaded from NCBI.

Presence of insertion sequences and prophages were predicted with IS finder (Siguier et al. 2006) and PHAST (Zhou et al. 2011), respectively, and comparing the obtained results with the manually curated annotation. The origin of the identified mobile elements was investigated by inspecting blastp hits on the whole NCBI nr protein database, and on selected subsections of nr based on taxonomy.

### Phylogenomic analyses

For phylogenomic analysis, two curated datasets of highly conserved orthologs were employed, namely one with 24 genes (gene set 1; Lang et al. 2013), and one with 120 genes (gene set 2; Parks et al. 2018), with representative taxon selections. In details for gene set 1, 23 organisms were selected, namely seven *Rickettsiales sensu stricto* (including *Deianiraea*), 13 other *Alphaproteobacteria* (including 2 *Holosporales*), plus three *Betaproteobacteria* and *Gammaproteobacteria* as outgroup. The amino acidic sequences of *Deianiraea* orthologs were identified by reciprocal blastp hits. Single orthologs were aligned with MUSCLE (Edgar 2004), polished with Gblocks (Talavera and Castresana 2007), and concatenated together, resulting in 4,038 characters. For gene set 2, 100 organisms were selected, namely 34 *Rickettsiales sensu stricto* (including *Deianiraea*), 49 other *Alphaproteobacteria* (including 11 *Holosporales*), plus 17 *Betaproteobacteria* and *Gammaproteobacteria* as outgroup. For this gene set, alternative organism sub-selections were tested, to account for possible long-branch attraction events in the full selection (Supplementary material 12), The alignment by Parks and co-authors (2018) was used. The amino acidic sequence of orthologs in *Deianiraea* and selected organisms not present in the initial dataset were selected by reciprocal blast hits, and added to the existing alignment with MAFFT (Nakamura et al. 2018), keeping the original alignment length. Thus, genes were concatenated, and 34,747 positions were selected according to Parks and co-authors (2018).

The best substitution model for each concatenated alignment was estimated with ProtTest (Darriba et al. 2011). Thus, ML and BI phylogenies were estimated respectively with PhyML (gene set 1) or RAxML (gene set 2; Stamatakis et al. 2015) and MrBayes.

### Metabolic prediction and comparison with other *Rickettsiales*

Metabolic pathways reconstruction was performed employing BioCyc (Caspi et al. 2016), Pathway tools (Karp et al. 2015), and KEGG (Kanehisa et al. 2016), followed by manual inspection. Cluster of orthology groups (COGs) were identified by the NCBI pipeline (Galperin et al. 2015) for *Deianiraea* and other 11 representative *Rickettsiales sensu stricto.* The repertoire of COGs and functional categories was directly compared among those organisms, to identify *Deianiraea* peculiar traits.

In particular, the biosynthetic pathways for amino acids of *Deianiraea* were reconstructed in detail by manual inspection of blastp results against other *Rickettsiales,* other bacteria and conserved domain search results (Marchler-Bauer et al. 2015) (Supplementary material 17, 18, 19).

### Horizontal gene transfer tests

The possibility of HGTof the genes assigned to the COGs present in *Deianiraea* and absent in all other *Rickettsiales* was evaluated. First, blastp hits on the whole NCBI nr protein database, and on selected subsections of nr based on taxonomy, were inspected. Only phylogenetically informative HGT-candidate genes (i.e. those with identity with best hits higher than ≥50%, and with at least 5 percentage points difference in the identities with the best hits on whole nr and on *Alphaproteobacteria*-only) were selected for phylogenetic analyses. Briefly, orthologs were identified on selected genomes, aligned and polished with MUSCLE and Gblocks. For each gene, ML phylogenetic analysis was performed with RAxML, after selection of the best model using ProtTest.

For the amino acid biosynthetic pathways exclusive in *Deianiraea*, detailed analyses were performed. After ortholog selection, alignment and polishing (as above detailed), for each biosynthetic pathway a concatenated alignment was produced. For each alignment, ML phylogenetic analyses were performed with RAxML, after selection of the best model using ProtTest, and compared with the respective organismal phylogeny. Additionally, the statistical deviation of GC content and CAI (codon adaptation index) (Puigbo et al. 2008) respect to the whole *Deianiraea* gene set were tested as described previously (Sassera et al. 2011).

## Data availability

Data supporting the findings of this study have been deposited in GenBank with the accession codes: *P. primaurelia* CyL4-1 partial 18S-ITS1-5.8S-ITS2-28S - MH185950, *P. primaurelia* CyL4-1 partial cytochrome oxidase subunit I gene - MH188082, *Deianiraea vastatrix* CyL4-1 partial 16S rRNA gene – MH197138; *Deianiraea vastatrix* CyL4-1 genome – CP028925 (genome sequence and annotation file uploaded together with manuscript submission for review use).

## Supporting information

## Acknowledgments

This work was funded by the European Commission FP7-PEOPLE-2009-IRSES grant 247658 (project CINAR-PATHOBACTER) to GP, the Human Frontier Science Program Grant RGY0075/2017 to DS, Italian Ministry of Education, University and Research (MIUR): Dipartimenti di Eccellenza Program (2018–2022) - Dept. of Biology and Biotechnology “L. Spallanzani”, University of Pavia to DS, and, for the work of the group from St Petersburg, by the RSF grant 16-14-10157 to AP. We would like to thank A. Oren for advice in bacterial nomenclature and species description. S. Gabrielli and S. Lometto are acknowledged for assistance in graphical artwork and in phylogenomic analyses, respectively. Cultures were maintained in Culture Collection of Microorganisms (RC CCM), Saint Petersburg State University.

## Author contributions

N.L. isolated the host cells and performed live experiments. E.S. performed TEM and fluorescence microscopy. K.B. performed AFM. O.L., A.P. and G.P. performed molecular experiments and probe design. M.C., O.L. and G.P. performed phylogeny. M.C. and O.L. performed IMNGS analyses. M.C., A.M.F. and D.S. performed genome assembly and analyses. M.C., A.M.F., S.G. and D.S. performed metabolic and evolutionary analyses. General interpretation of the results and manuscript writing were performed by M.C., D.S. and G.P., and contributed by C.B., L.M., E.S. and A.P. All the authors contributed to the interpretation and writing of the specific results, and to the discussions. G.P. planned and coordinated the whole project, with D.S. planning and coordinating the genomic analyses, and E.S. coordinating the microscopy.

