## Supplementary material for "The extracellular association of the bacterium “*Candidatus* Deianiraea vastatrix” with the ciliate *Paramecium* suggests an alternative scenario for the evolution of *Rickettsiales*"

**Supplementary material legends**

S**upplementary material 1:** Differential Interference Contrast (DIC) (**a**) and Atomic-Force Microscopy (AFM) (**b**) images of healthy *Paramecium*-cells, non-infected by *Deianiraea.* In (**a**), numerous cilia (white arrowhead) are visible all around the cell. In the AFM (**b**), the brightness is proportional to the tridimensional height of the structure (i.e. the more exposed, the brighter). In (**b**), cortical alveoli (each bearing a cilium) are visible (black arrowhead). Scale bars: 10 μm (**a**), shown in figure (**b**).

**Supplementary material 2:** TEM (**a, b**) and AFM (**c**) of CyL4-1 cells infected by *Deianiraea*. Numerous cytoplasmic lysosomes (red arrowhead) are visible in (**a**), while in (**b**) ring-like nucleoli are shown (red arrow). In the AFM (**c**), the brightness is proportional to the tridimensional height of the structure (i.e. the more exposed, the brighter). In (**c**) a detail of the surface of a CyL4-1 covered by a dense overlay of *Deianiraea* bacteria (black arrow) is presented. Scale bars stand for 200 nm (**a**) and 1 μm (**b**), respectively, while in (**c**) scale is shown in figure.

**Supplementary material 3**: List of the strain employed as recipients in the *Deianiraea* transfer experiment, their environmental origin, species, culture collection, and their susceptibility in the experiment.

**Supplementary material 4:** FISH pictures of CyL4-1 cells infected by *Deianiraea*. (**a-c**) The same cell shown in Main Figure 1, observed from a superficial plane; (**d-e**) A different cell shown from a transversal plane. (**a,d**) The signal of DAPI. (**b,e**) FITC-conjugated EUB338 probe; (**c**) the Cy3-conjugated specific probe Deia_416; (**f**) the alternative Cy3-conjugated specific probe Deia_536. Scale bars stand for 10 μm.

**Supplementary material 5:** Full maximum likelihood (**a**) and Bayesian inference (**b**) trees inferred on the 16S rRNA gene alignment of *Rickettsiales* plus outgroup (condensed Bayesian shape with support values shown in main Figure 3). Numbers on branches report bootstrap values (**a**) and posterior probabilities (**b**). The four families and the two sub-clades of family *Deianiraeaceae* are evidenced. Scale bars stand for estimated sequence divergence of 10%.

**Supplementary material 6:** Identity table on the same 16S rRNA gene alignment used for phylogeny. The newly characterized sequence, as well as all self-identities, are reported in bold. The black square outline encloses all *Rickettsiales* and their reciprocal identities, while the members of each family and their reciprocal identities are highlighted in different colours. The red square outlines enclose the two *Deianiraeaceae* sub-clades.

**Supplementary material 7:** Table showing, for each *Rickettsiales* family, the number of OTUs reported according to the prevalent environmental origin according to IMNGS categories (non-integers result from tied environmental origins), and the proportion relative to the total OTUs of the respective family.

**Supplementary material 8:** General features of all *Rickettsiales* genomes (excluding metagenomic samples) downloaded from NCBI. The genomes are sorted according to the descending coding density. Values relative to *Deianiraea* are reported in bold. The highest and lowest value for each column are reported in bold and underlined. Italicized values refer to the genome of "*Candidatus* Arcanobacter lacustris", estimated to be incompletely sequenced.

**Supplementary material 9:** List of phage-related sequences found in the genome of *Deianiraea*. The two putative prophage regions retrieved by PHAST are reported, with the putative phage genes (bold), phage attachment sites (italics), on the basis of which the putative prophage regions were delineated by PHAST. All other intervening genes, i.e. any other genes internal to the prophage regions, are shown in regular typeface. A list of additional putative phage genes identified by manual annotation in the *Deianiraea* genome follows. For all putative phage genes, best blast hit on NCBI nr is reported.

**Supplementary material 10:** Putative transposases and possible accessory genes (potentially involved in transposition) identified by IS finder. Start and end position refer to the combined largest segment covered by significant hits in the same genome region. The e-value in the IS finder search of the best hit only is shown. In addition, for each putative element identified by IS finder, the corresponding manually annotated gene and the best blast hit on NCBI nr protein (including % identity and coverage) are reported.

**Supplementary material 11:** Full maximum likelihood (**a**) and Bayesian inference (**b**, same tree shown in main figure 4, here reported for comparison) trees of *Alphaproteobacteria* inferred with the LG+I+G model on the concatenated amino acidic alignment of 24 highly conserved genes (gene set 1; Lang et al. 2013). Numbers on branches indicate bootstrap values with 1,000 pseudo-replicates (**a**) and posterior probabilities after 1,000,000 iterations (**b**). The members of the order *Rickettsiales*, those of the order *Holosporales*, the other *Alphaproteobacteria*, and the outgroup are evidenced. Scale bars stand for estimated sequence divergence.

**Supplementary material 12:** Full maximum likelihood (**a, c, e**) and Bayesian inference (**b**, **d**, **f**) trees of *Alphaproteobacteria* inferred with the LG+I+G+F model on the concatenated amino acidic alignment of 120 highly conserved genes (gene set 2; Parks et al. 2018) on three different taxa datasets. Numbers on branches report bootstrap values with 1,000 pseudo-replicates (**a, c, e**), and posterior probabilities after 100,000 iterations (**b, d, f**); values of 100 (**a, c, e**) or 1.00 (**b, d, f**) were omitted. (**a**, **b**) include the whole taxonomic selection (100 organisms), (**c**, **d**) the same selection excluding “*Ca*. Fokinia solitaria”, and (**e**, **f**) the same selection as in (**a**, **b**) but excluding *Deianiraea*. The members of the order *Rickettsiales*, those of the order *Holosporales*, the other *Alphaproteobacteria*, and the outgroup are evidenced in black, members of each *Rickettsiales* family in red. While in (**a**, **b**), *Midichloriaceae* are non-monophyletic, in (**c**-**f**) all the *Rickettsiales* families are monophyletic, consistently with 16S rRNA phylogenies (main figure 3, Supplementary material 5). These analyses indicate a long-branch attraction phenomenon between the two fastest evolving sequences among *Rickettsiales* in (**a**, **b**), i.e. *Deianiraea* and “*Ca*. Fokinia solitaria”, which leads to an artefactual reconstruction in (**a**, **b**). Scale bars stand for estimated sequence divergence.

**Supplementary material 13:** Complete list of all COGs of *Deianiraea* sorted by functional category, obtained using NCBI COG pipeline, and the respective annotated genes.

**Supplementary material 14:** Number of COGs assigned to each functional category for *Deianiraea* and other selected *Rickettsiales*, including the percentage over the total for each organism.

**Supplementary material 15:** COG category comparison among *Rickettsiales* families. Each row stands for the exclusive COG repertoire of a *Rickettsiales* family, or the COGs shared by two or more families. The presence in a single family member is sufficient condition for inclusion. COG numbers for each row are reported subdivided by functional category, including the percentage over the row total. The total and exclusive *Deianiraea* COGs are highlighted in bold and red, respectively. The COG numbers of functional category E (Amino acid transport and metabolism) are evidenced in yellow.

**Supplementary material 16:** Complete list of all COGs found exclusively in *Deianiraea* respect to other *Rickettsiales*, sorted by functional category, and the respective annotated genes. The best hits on NCBI nr proteins and on selected taxonomic subsections (*Proteobacteria*, *Alphaproteobacteria*, *Rickettsiales*) are shown for each gene (except those involved in amino acids biosynthesis, detailed in Supplementary material 19). Genes selected for phylogenetic tests (Supplementary material 21) are italicized.

**Supplementary material 17:** Detailed description of analysis on amino acid biosynthetic pathways, including phylogenetic trees testing potential horizontal gene transfer.

**Supplementary material 18:** Schematic drawing of the reconstructed amino acid biosynthetic pathways in *Deianiraea*. Compounds are shown in regular typeface, enzymes are italicized, with, when present, gene products of *Deianiraea* inferred to perform the reaction (multimeric enzymes are indicated, while slashed genes indicate functional analogs). Precursor molecules are shown in blue, “trusted” reactions (and their products) exerted by *Deianiraea* are shown in green, “putative” reactions in orange, while non-exerted reaction (and their products) in red. Reactions performed exclusively by *Deianiraea* among *Rickettsiales* are enclosed in dotted boxes. Final amino acidic products are shown in bold, and the eight amino acids produced exclusively by *Deianiraea* among *Rickettsiales* are encircled by green outlines.

**Supplementary material 19:** Table listing all genes for amino acid metabolism described in Supplementary material 17 according to their respective pathway. Note that all but one pathway are not encoded in operons, thus are less likely to be the the result of a recent Horizontal Gene Transfer (HGT). The best blastp hits are also shown on the entire nr database, *Proteobacteria* only, *Alphaproteobacteria* only, and *Rickettsiales* only (red lettering indicates hits identified as non-orthologs). The genes used for the concatenated phylogenetic analyses are highlighted in bold. The results of statistical tests on GC content and Codon Adaptation Index (CAI) deviation are presented, together with a summary of the outcome of the phylogeny-based HGT test.

**Supplementary material 20:** List of all *Deianiraea* ORFs for which SignalP identified signal peptides, including the outcome of the TMHMM analysis, and the consequent inference as 55 “putative secreted effectors” and four “transmembrane proteins”.

**Supplementary material 21:** Maximum likelihood phylogenetic trees of selected *Deianiraea*-exclusive genes (see Supplementary material 16) inferred with RaxML, with 1,000 pseudo-replicates (bootstrap support reported on branches), and the Prottest-selected substitution model: (**a**) three SpoVG-like proteins: LG+I+G+F model; (**b**) Putative ImmA/IrrE family metallo-endopeptidase: BLOSUM62+I+G+F model; (**c**) DNA mismatch repair protein MutH: LG+I+G model; (**d)** Putative type III restriction-modification DNA methylase**:** LG+I+G model. The *Deianiraea* genes are evidenced in bold. Scale bars stand for sequence divergence (0.1 unless specified directly). In all cases, the low phylogenetic signal, clearly shown by the low support of most branches, did not allow to confidently resolve the tree structure.

**Supplementary material 22:** List of best blastn hits and respective identities for the three molecular markers used to characterize the *Paramecium* hosting *Deianiraea.*

**Supplementary material 23:** Number of the aspecific hits retrieved on RDP and SILVA searches for the two *Deianiraea*-specific 16S rRNA-targeted FISH probes, namely Deia_416 (5'-GAG TTT TAC AAT CTT TCG-3') and Deia_538 (5'-AGT AAC GCT TGG ACT CCA-3').

**Supplementary material 24:** Alignment in fasta format of the 1,398 nucleotide positions employed in the 16S rRNA gene phylogenies (main figure 3; Supplementary material 5).

**Supplementary material 25:** List of the 16S rRNA gene-based similarity clusters obtained for each *Rickettsiales* family on SILVA database. The total number of sequences included in each cluster is reported, as well as the SILVA “name” and NCBI accession of the sequence chosen as respective representative. The representative sequences were used to query the IMNGS database.

**Supplementary material 26:** Detailed description of the sequencing and assembly procedures of the *Deianiraea* genome.

**Supplementary material 27:** List of the contigs of the preliminary assembly in which SSU rRNA genes were identified with barrnap. For each contig, Blobology parameters are reported (see Supplementary material 28), as well as the positions in which the SSU rRNA gene was inferred, and the best blast hit of this gene sequence. Contigs are sorted by organismal assignment (green: *Deianiraea*; blue: *Paramecium*; red: other bacteria; grey: other eukaryotes), and, within each group, by descending sequencing coverage (upper and lower values for *Deianiraea* are highlighted in red).

**Supplementary material 28:** Classification of the contigs from the preliminary assembly according to the Blobology pipeline, i.e. length, GC content, sequencing coverage, and taxonomy of the best megablast hit on NCBI Nucleotide. The contigs are split into two parts, part 1 includes all the contigs of the “Final selection from preliminary assembly” (highlighted in yellow the 3 contigs added respect to the “Initial selection from preliminary assembly”), part 2 includes all the remaining contigs of the preliminary assembly.

**Supplementary material 29:** List of the contigs of the “First reassembly” in which SSU rRNA genes were identified with barrnap. For each contig, the positions in which the SSU rRNA gene was inferred, and the best blast hit of this gene sequence, are reported. Contigs are sorted by organismal assignment (green: *Deianiraea*; blue: *Paramecium*; red: other bacteria).

**Supplementary material 30:** List of primers used for genome closing of *Deianiraea*, as well as the respective PCR protocols.
