## Supplementary material for "The extracellular association of the bacterium “*Candidatus* Deianiraea vastatrix” with the ciliate *Paramecium* suggests an alternative scenario for the evolution of *Rickettsiales*": Suppl_17_new_Aminoacid_biosynthesis.docx

**Supplementary material: biosynthetic pathways of amino acids**

This section supplements the main text describing the procedures and results regarding the reconstruction of the biosynthetic pathways for amino acids in *Deianiraea*, and the phylogenetic analyses of the genes involved.

The prediction of the function of each gene putatively involved in the pathways was verified in detail by manual inspection of blastp results against NCBI databases, and conserved domain search results (Marchler-Bauer et al. 2015). The pathways were reconstructed manually with the reference of Biocyc (Caspi et al. 2016), Pathway tools (Karp et al. 2015), and KEGG (Kanehisa et al. 2016), which were also used for comparing with other bacteria. In particular, the presence in other *Rickettsiales* (excluding samples coming from metagenomic projects) of all the genes involved in amino acid biosynthesis (both those present in *Deianiraea* and those absent) was further directly evaluated by taxon-specific blastp searches. In order to consider a biosynthetic pathway present in a given organism, two main criteria were applied: the gene coding to the last enzymatic step, giving the final compound, must be present, and half (or more) of the genes specific of the pathway should be present.

Further analyses were conducted on the five biosynthetic pathways, responsible for the synthesis of eight amino acids, which were identified as exclusive of *Deianiraea* respect to all known *Rickettsiales*. In particular, phylogenetic analyses were employed to evaluate the evolutionary origin of those genes and the possibility of horizontal gene transfer (HGT). In order to identify the orthologs, the predicted proteins from 2644 complete bacterial genomes were downloaded from NCBI. For each *Deianiraea* protein analysed, the respective best blastp hit on each organismal protein set was identified. In order to filter out paralogs, multiple strategies were employed. First, preliminary screening of the best hits from the NCBI annotation was performed, followed by examination of preliminary single gene trees (data not shown). Additionally, in the cases when *Deianiraea* itself possessed two (or more) paralogous genes, the respective best hits on each organism were compared, and only the hit with highest identity was retained. Finally, all the *Deianiraea* genes that were found to have orthologs in an insufficient and not taxonomically representative set of organisms (in particular for what concerned the presence in other *Alphaproteobacteria*) were discarded from the analysis. Subsequently, the retained genes of each pathway were treated separately. For each pathway, a representative selection of proteobacterial organisms plus an outgroup was selected for the analysis. Then, each ortholog was aligned with MUSCLE and polished with Gblocks, and a concatenated alignment of all orthologs was produced. For each alignment, ML (maximum likelihood) phylogenetic analyses were performed with RAxML, after selection of the best model using ProtTest. For comparison, the respective organismal phylogenies were inferred, by selecting single copy orthogroups with OrthoFinder (Emms and Kelly 2015), which were aligned, polished and concatenated before ML and BI phylogenies, as described above. The Shimodaira-Hasegawa (SH) test was performed with RAxML, in order to compare the phylogeny of each pathway with the respective organismal phylogeny.

In the following sections, a detailed report is presented on the inferred reconstruction of the five metabolic pathways exclusive of *Deianiraea* among *Rickettsiales,* followed by the respective phylogenetic results.

**Aromatic amino acids synthesis**

Biosynthesis of aromatic amino acids (tryptophan, phenylalanine, tyrosine) in bacteria initially involves the shikimate pathway. This pathway, starting from erythrose 4-phosphate (an intermediate of the pentose-phosphate pathway) and phosphoenolpyruvate (and requiring another phosphoenolpyruvate at the 6^th^ step), leads to chorismate through 7 enzymatic steps (Pittard and Yang 2008; Bender 2012). In turn, chorismate leads to the synthesis of the aromatic amino acids and other aromatic compounds. The gene sequence of the enzyme catalysing the first step of the shikimate pathway (3-deoxy-7-phosphoheptulonate synthase) is atypical respect to all *Proteobacteria*. Only a limited number of significant blastp hits was found, and always with very low identity (Supplementary material 16.2), with sequences assigned to rare bacterial phyla retrieved in metagenomic samples (Anantharam et al. 2016; Lawson et al. 2017; Probst et al. 2018; Brown C.T et al. unpublished: [KKT47881.1]).

Additionally, a peculiar gene fusion of enzymatic activities was retrieved (Deia_00424), namely for the genes encoding in the order the enzymes of the 5^th^, 4^th^ and 3^rd^ of the shikimate pathway (respectively, shikimate kinase AroL, shikimate dehydrogenase AroE, and 3-dehydroquinate dehydratase AroD). As a matter of fact, several different combinations of gene fusions involved in the shikimate pathway are common in different evolutionary lineages (Bender 2012). Interestingly, the 3-dehydroquinate dehydratase is more similar to type I enzymes found in *Gammaproteobacteria* and *Betaproteobacteria*, rather than to type II, typical in other *Alphaproteobacteria*. The gene for the 6^th^ step of the pathway (3-phosphoshikimate 1-carboxyvinyltransferase) is present also in two *Anaplasmataceae*, namely “*Ca*. Neoehrlichia lotoris” (Daugherty, S.C et al. unpublished: [LANX01000001]) and “*Ca*. Xenolissoclinum pacificiensis” (Kwan and Schmidt 2013).

The biosynthetic pathway for tryptophan immediately deviates after chorismate (five genes for six enzymatic steps in *Escherichia coli*). This pathway is absent in *Deianiraea* and in all other *Rickettsiales*: only a single gene (anthranilate phosphoribosyltransferase TrpD) was found in the *Rickettsiales* bacterium Ac37b (Felsheim, R.F et al. unpublished. [NZ_CP009217]).

The synthesis of both tyrosine and phenylalanine involves the conversion of chorismate into prephenate by a chorismate mutase. The following step is specific for each amino acid, involving prephenate dehydratase for phenylalanine, and prephenate/cyclohexenyl dehydrogenase for tyrosine. Similarly to other *Alphaproteobacteria*, in *Deianiraea* these three enzymatic activities are encoded by three distinct genes, differently from the other *Proteobacteria* classes, presenting two genes, in which different isoforms of chorismate mutase are fused with the each of the other two enzymes. The last step is the amination of both tyrosine and phenylalanine, performed by a generalist amino acid aminotransferase (e.g. Deia_00080, Deia_00820, Deia_00902).

**Phylogenetic analyses**: Used eight genes (operationally subdividing Deia_00424 into three distinct genes according to the homology-predicted enzymatic activities) and 71 organisms, 1,250 sites. For the phylogenomics, used 59 ortholog genes and 10,126 sites.

**Figure 16.1** Maximum likelihood tree of the eight concatenated genes involved in the aromatic amino acids biosynthesis with the LG+I+G+F substitution model with 100 bootstrap pseudo-replicates. The position of *Deianiraea* genes is unsupported. Scale bar stands for estimated sequence divergence. Number on branches stand for bootstrap values.

**Figure 16.2** Reference maximum likelihood phylogenomic tree of the organisms employed for phylogenesis in **16.1** with the LG+I+G substitution model with 100 bootstrap pseudo-replicates. The five proteobacterial classes (as indicated in the figure) are all monophyletic and highly supported, including the positioning of *Deianiraea* within *Alphaproteobacteria*, which is consistent with the analyses shown in **Supplementary material 11**. Scale bar stands for estimated sequence divergence. Number on branches stand for bootstrap values.

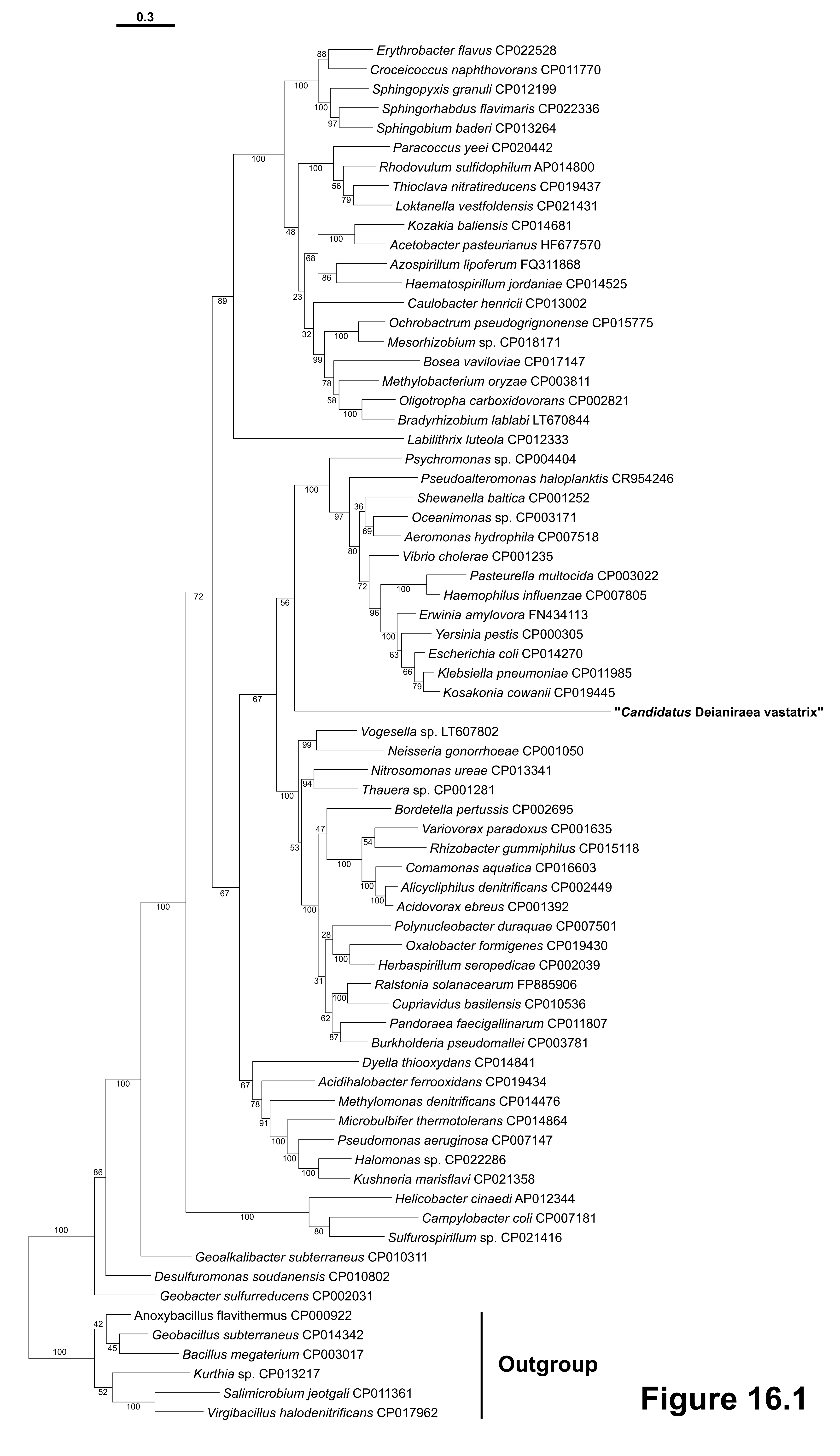

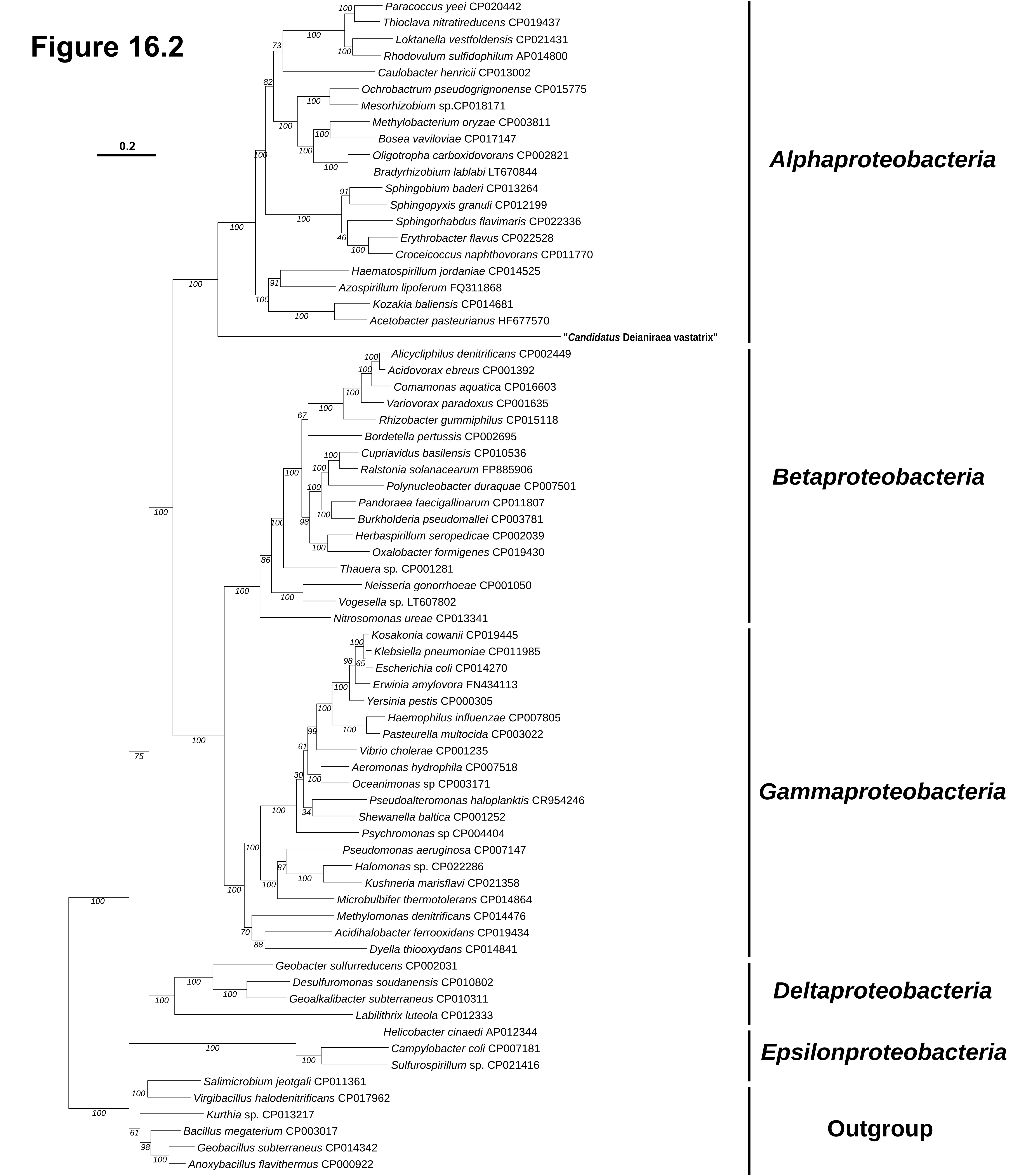

**Branched chain amino acids synthesis**

The biosynthesis of the three branched chain amino acids (valine, leucine, isoleucine) involves a number of shared enzymes (Salmon et al. 2006). Typically, 3-methyloxybutanoate, common intermediate of leucine and valine, is obtained in three enzymatic steps from two pyruvate molecules. Two isozymes working in the first step were found in *Deianiraea* (Deia_00067 acetolactate/acetohydroxybutanoate synthase large subunit; Deia_01102 acetohydroxybutanoate/acetolactate synthase large subunit). A putative homologue is present in “*Occidentia massiliensis*” (*Rickettsiaceae*) (Mediannikov et al. 2014). Another ORF (Deia_1007) with similarities to the accessory regulatory subunit of acetolactate synthase was identified, but, since the alignment involved only partial length of database reference sequences (data not shown), it was judged non-functional and not further considered. The gene for the second step of the pathway (ketol-acid reductoisomerase) is also present in few *Anaplasmataceae*, namely “*Ca*. Neoehrlichia lotoris” (Daugherty, S.C et al. unpublished [LANX01000001]) and *Anaplasma* spp. (Dark et al. 2009).

For the synthesis of valine from 3-methyloxybutanoate, a single additional step is needed, exerted by the branched-chain aminotransferase. Contrarily to other genes of the pathway, this is present in several *Rickettsiales*, having a likely a broader generalist amino acid aminotransferase activity, or possibly acting on the biosynthetic intermediates derived from hosts.

For leucine biosynthesis from 3-methyloxybutanoate, three specific enzymatic reactions are necessary, involving four proteins (two enzymes and a dimeric complex formed by the other two proteins), plus a fourth terminal amination step performed by the same branched-chain aminotransferase as for valine.

The most common bacterial biosynthetic pathway for isoleucine starts from the deamination of threonine, to produce 2-oxobutanoate. This compound is then processed by the same sequential four enzymes as for valine (including terminal aminotransferase), to obtain isoleucine. The specific gene for the initial step (threonine deaminase IlvA) is not present in *Deianiraea*, differently from other *Alphaproteobacteria*. A detailed inspection allowed to determine that ORF Deia_00356, initially flagged as an additional homologue of the leucine synthesis protein 2-isopropylmalate synthase, displays higher homology and similar domain architecture to citramalate synthase (data not shown). Citramalate synthase is involved in an alternative pathway for the synthesis of isoleucine found in a growing number of bacteria (Xu et al. 2004; Risso et al. 2008), including *Alphaproteobacteria* (Tang et al. 2009; McKinlay et al. 2010). Starting from pyruvate and acetyl-CoA, citramalate is formed. This compound is then processed by the two terminal leucine synthesis enzymes (excluding the aminotransferase) to obtain 2-oxobutanoate, which can then enter the canonical pathway. As possible additional alternative, methionine can also be a source of 2-oxobutyrate, through the hydrolytic action of methionine gamma-synthase/lyase.

**Phylogenetic analysis:** Used eight genes and 73 organisms, 2,674 sites. For the phylogenomics, used 42 ortholog genes and 6,630 sites.

**Figure 16.3** Maximum likelihood tree of the eight concatenated genes involved in the branched amino acids biosynthesis with the LG+I+G substitution model with 100 bootstrap pseudo-replicates. The position of *Deianiraea* genes is consistent with organismal phylogeny in **16.7** with high support. Scale bar stands for estimated sequence divergence. Number on branches stand for bootstrap values.

**Figure 16.4** Reference maximum likelihood phylogenomic tree of the organisms employed for phylogenesis in **16.3** with the LG+I+G substitution model with 100 bootstrap pseudo-replicates. The five proteobacterial classes (as indicated in the figure) are all monophyletic and supported, including the positioning of *Deianiraea* within *Alphaproteobacteria*, which is consistent with the analyses shown in **Supplementary material 11**. Scale bar stands for estimated sequence divergence. Number on branches stand for bootstrap values**.**

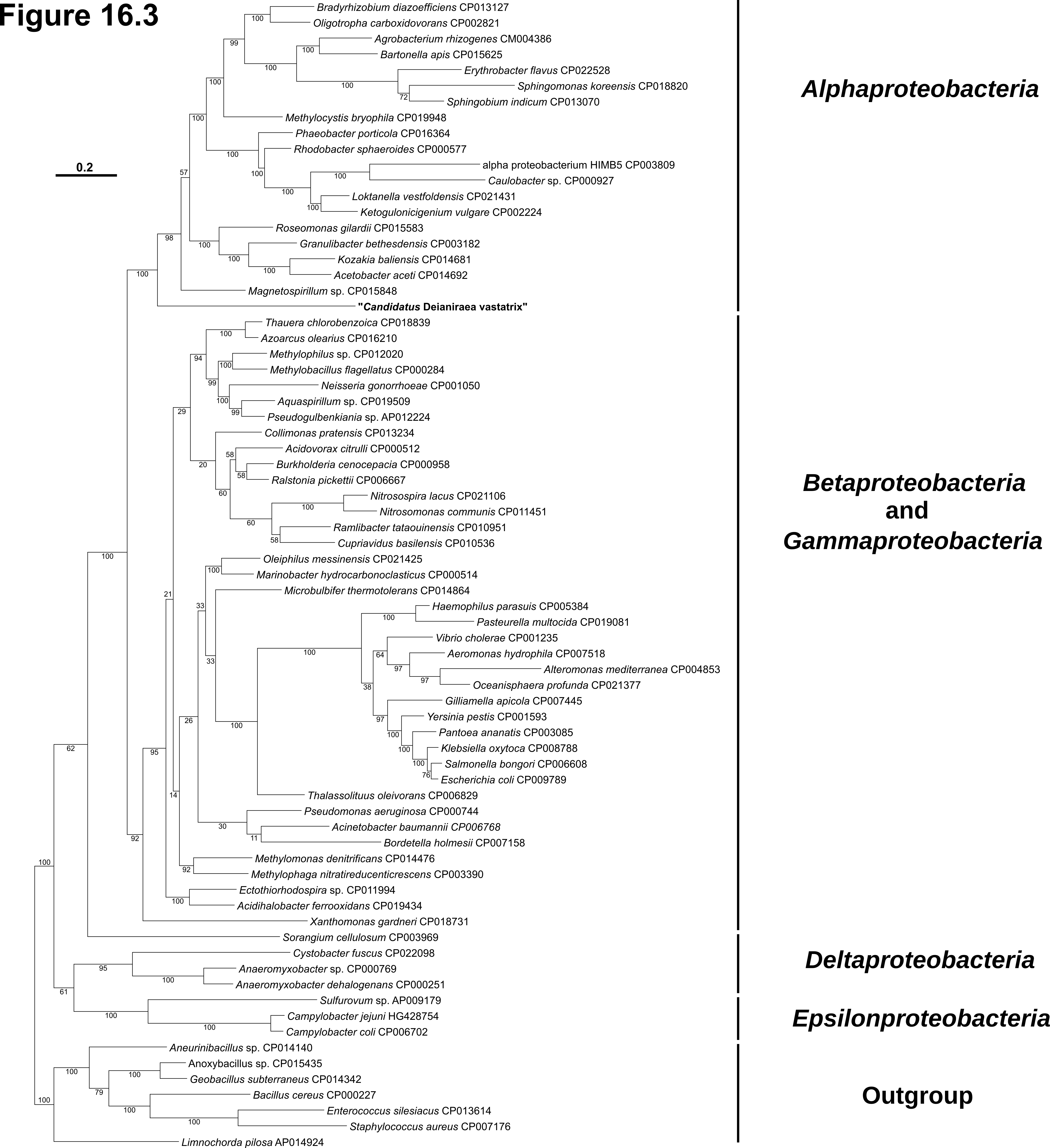

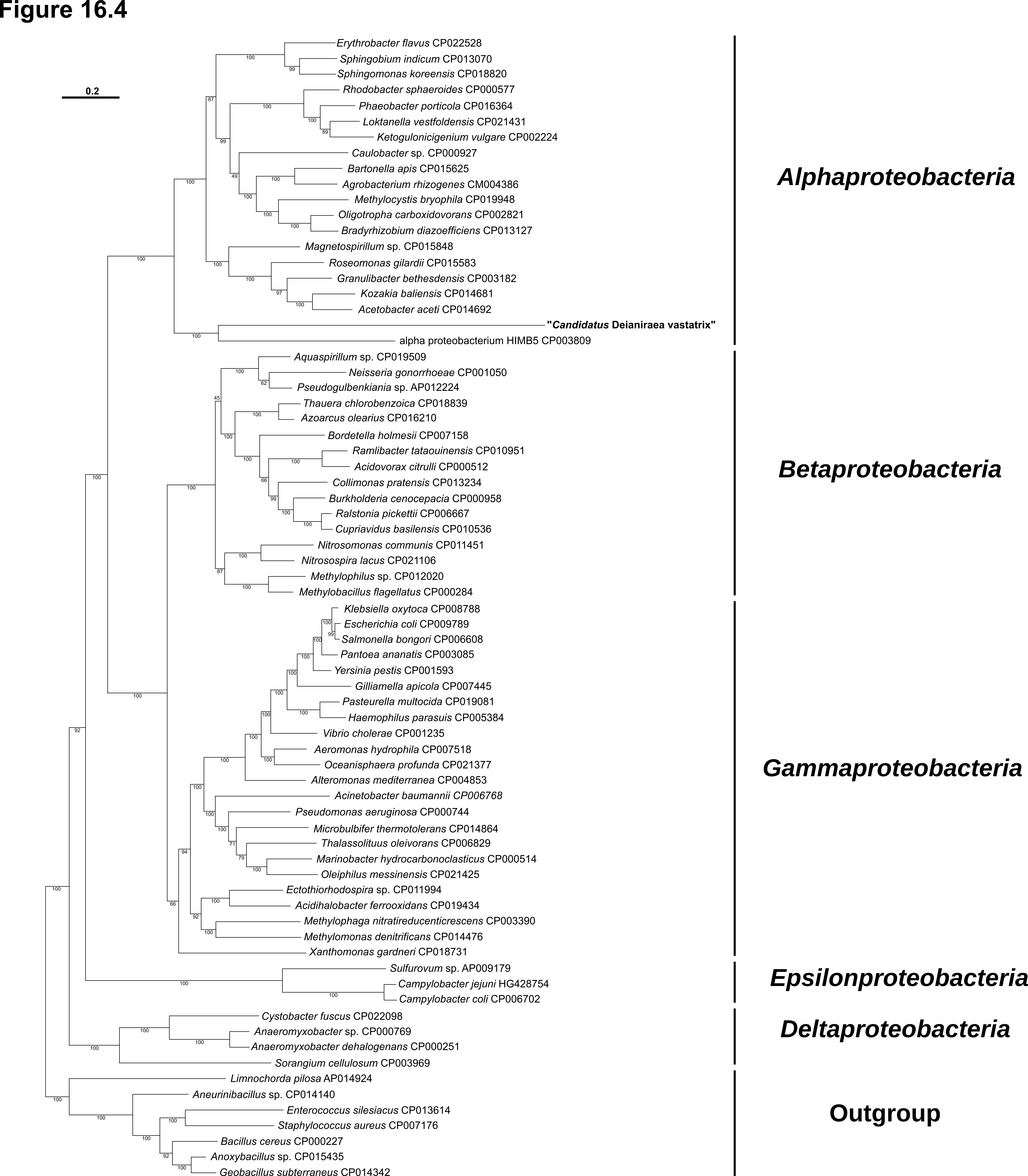

**Cysteine synthesis**

In most bacteria cysteine biosynthesis is exerted by a two-step pathway, starting from serine. In the *Deianiraea* genome only the second enzyme is encoded (Deia_00050: O-acetylserine sulfhydrylase/cysteine synthase A CysK). Thus, its substrate(s) acetyl-serine (or phosphoserine) should be either directly imported from *Paramecium*, or produced by an unidentified acetyl-(or phosho)-transferase acting on serine (instead of the canonical serine acetyltransferase/kinase, absent in *Deianiraea*). Alternatively, cysteine might also be produced from methionine via the reversed trans-sulfuration pathway (**methionine synthesis** pathway) (Bender 2012; Makino et al. 2016)

**Phylogenetic analyses**: Used one gene and 69 organisms, 261 sites. For the phylogenomics, used 49 ortholog genes and 6,009 sites.

**Figure 16.5** Maximum likelihood tree of the gene involved in the cysteine biosynthesis with the LG+I+G substitution model with 100 bootstrap pseudo-replicates. The position of *Deianiraea* genes and the main proteobacterial classes are unsupported. Scale bar stands for estimated sequence divergence. Number on branches stand for bootstrap values.

**Figure 16.6** Reference maximum likelihood phylogenomic tree of the organisms employed for phylogenesis in **16.5** and with the LG+I+G substitution model with 100 bootstrap pseudo-replicates. The five proteobacterial classes (as indicated in the figure) are all monophyletic and highly supported, including the positioning of *Deianiraea* within *Alphaproteobacteria*, which is consistent with the analyses shown in **Supplementary material 11**. Scale bar stands for estimated sequence divergence. Number on branches stand for bootstrap values.

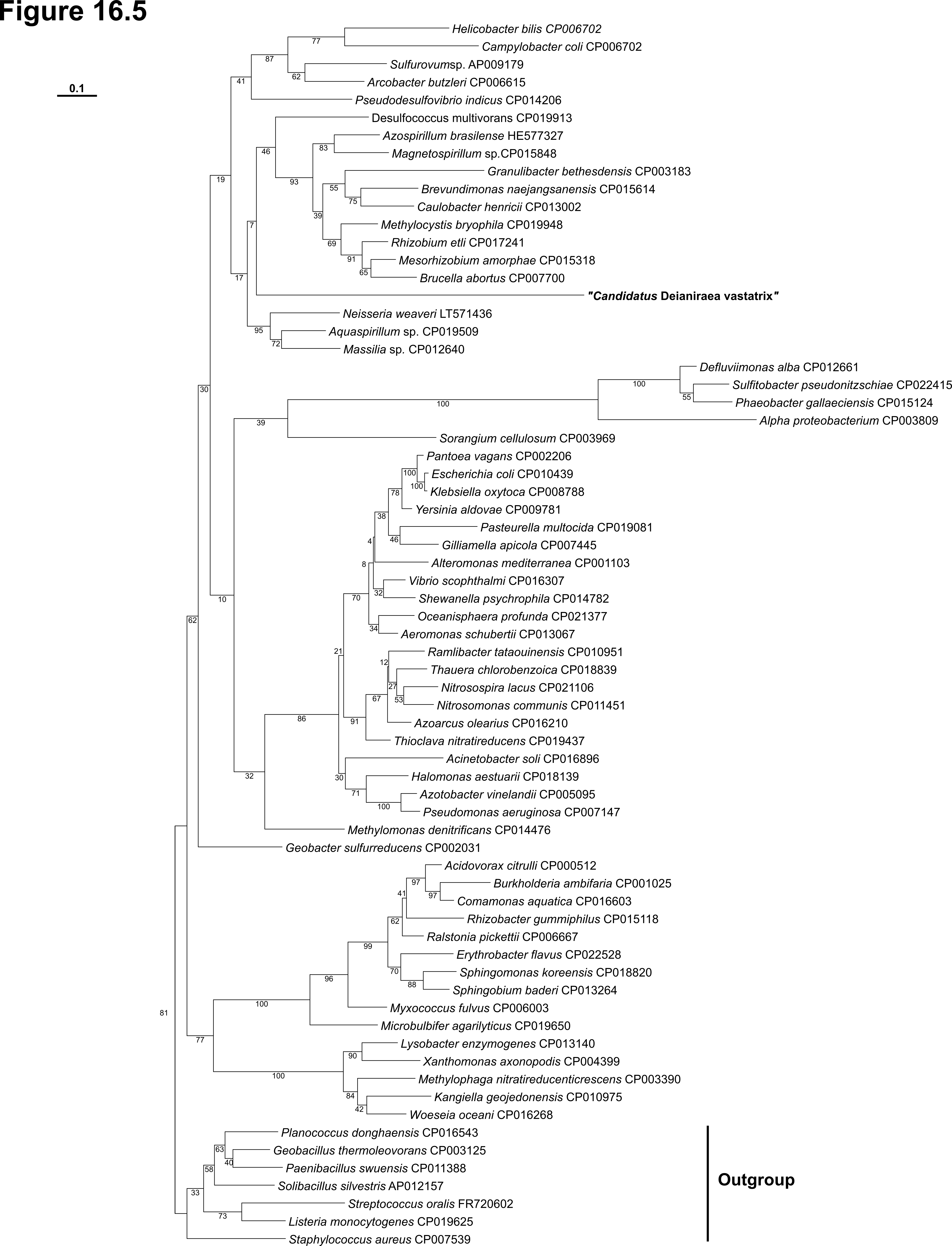

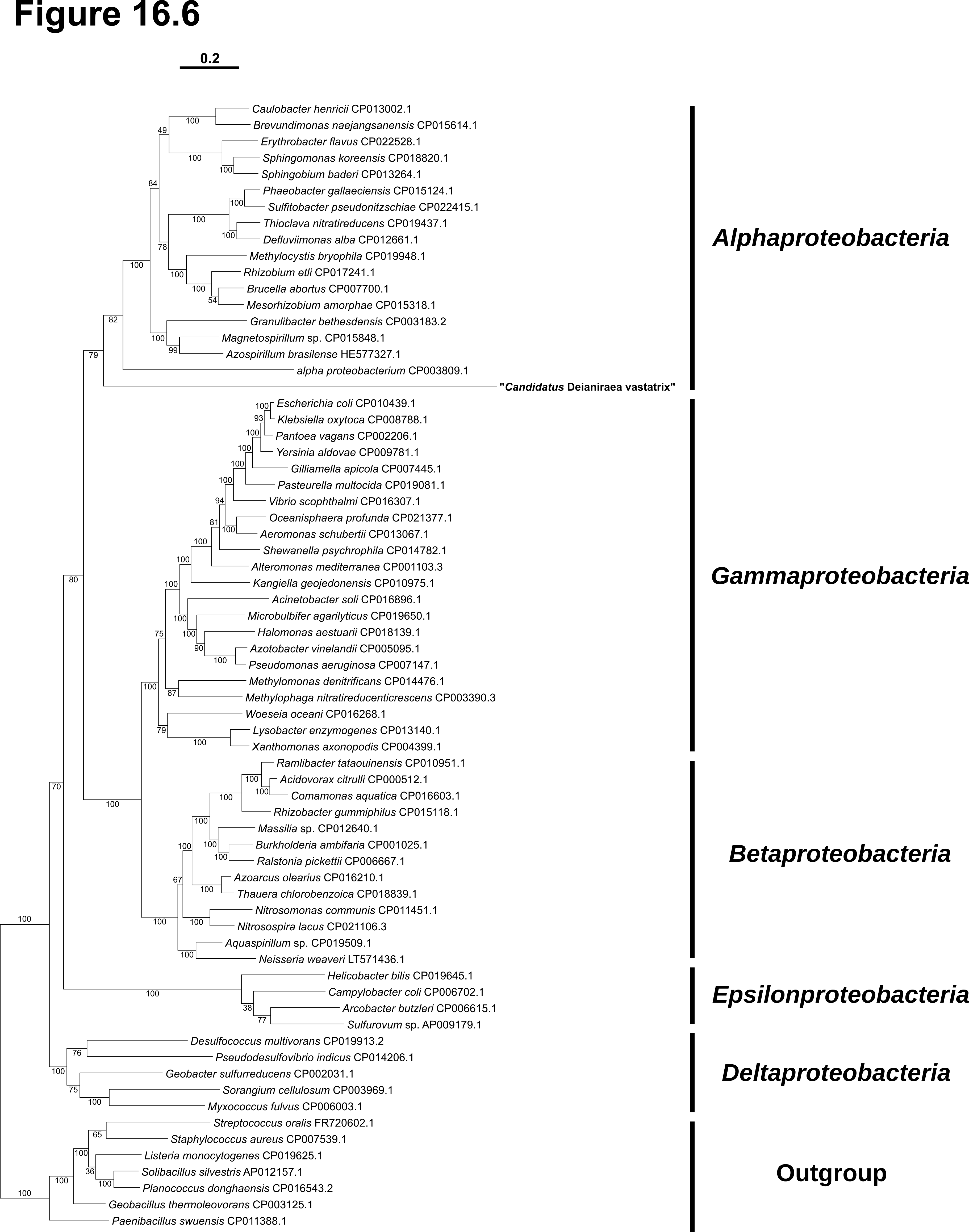

**Histidine synthesis**

Biosynthesis of histidine in bacteria is a multi-step pathway, involving ten enzymatic reactions, starting from 5-phosphoribosyl diphosphate, in turn obtained from the pentose phosphate intermediate ribose-5-phosphate via the ribose-phosphate diphosphokinase (Bender 2012; Winkler and Ramos-Montañez 2009). Depending on the organism, the number of genes involved in the pathway typically varies from eight to ten, given that alternative combinations of fused or separated of enzymatic activities for two pair of genes (regarding steps 2^nd^+3^rd^ and 6^th^+8^th^) might occur. *Deianiraea* possesses eight genes, with the two fused combinations. In particular, the presence of the bifunctional fused imidazoleglycerol-phosphate dehydratase/histidinol-phosphatase HisB (6^th^+8^th^ steps) is peculiar. Indeed, almost all *Alphaproteobacteria* possess separated genes for those enzymatic activities (with possible exceptions (Tully et al. 2017)), while the fused HisB is more common in other proteobacterial lineages, such as *Gammaproteobacteria* and *Epsilonproteobacteria*. Among *Rickettsiales*, the gene for the enzyme catalysing the 7^th^ step (Deia_00820 Histidinol-phosphate aminotransferase) finds a counterpart in *Rickettsiales* bacterium Ac37b (Felsheim, R.F et al. unpublished. [NZ_CP009217]), and possibly in “*Candidatus* Rickettsia asemboensis” (Jima et al. 2015).

**Phylogenetic analyses** Used nine genes (operationally subdividing HisB gene into two distinct genes according to the homology-predicted enzymatic activities) and 72 organisms, 1,123 sites. For the phylogenomics, used 56 ortholog genes and 7,917 sites.

**Figure 16.7** Maximum likelihood tree of the nine concatenated genes involved in the histidine biosynthesis with the LG+I+G+F substitution model with 100 bootstrap pseudo-replicates. The *Deianiraea* genes are very far-related to any other organism. Scale bar stands for estimated sequence divergence. Number on branches stand for bootstrap values

**Figure 16.8** Reference maximum likelihood phylogenomic tree of the organisms employed for phylogenesis in **16.7** with the LG+I+G substitution model with 100 bootstrap pseudo-replicates. The five proteobacterial classes (as indicated in the figure) are all monophyletic and highly supported, including the positioning of *Deianiraea* within *Alphaproteobacteria*, which is consistent with the analyses shown in **Supplementary material 11**. Scale bar stands for estimated sequence divergence. Number on branches stand for bootstrap values.

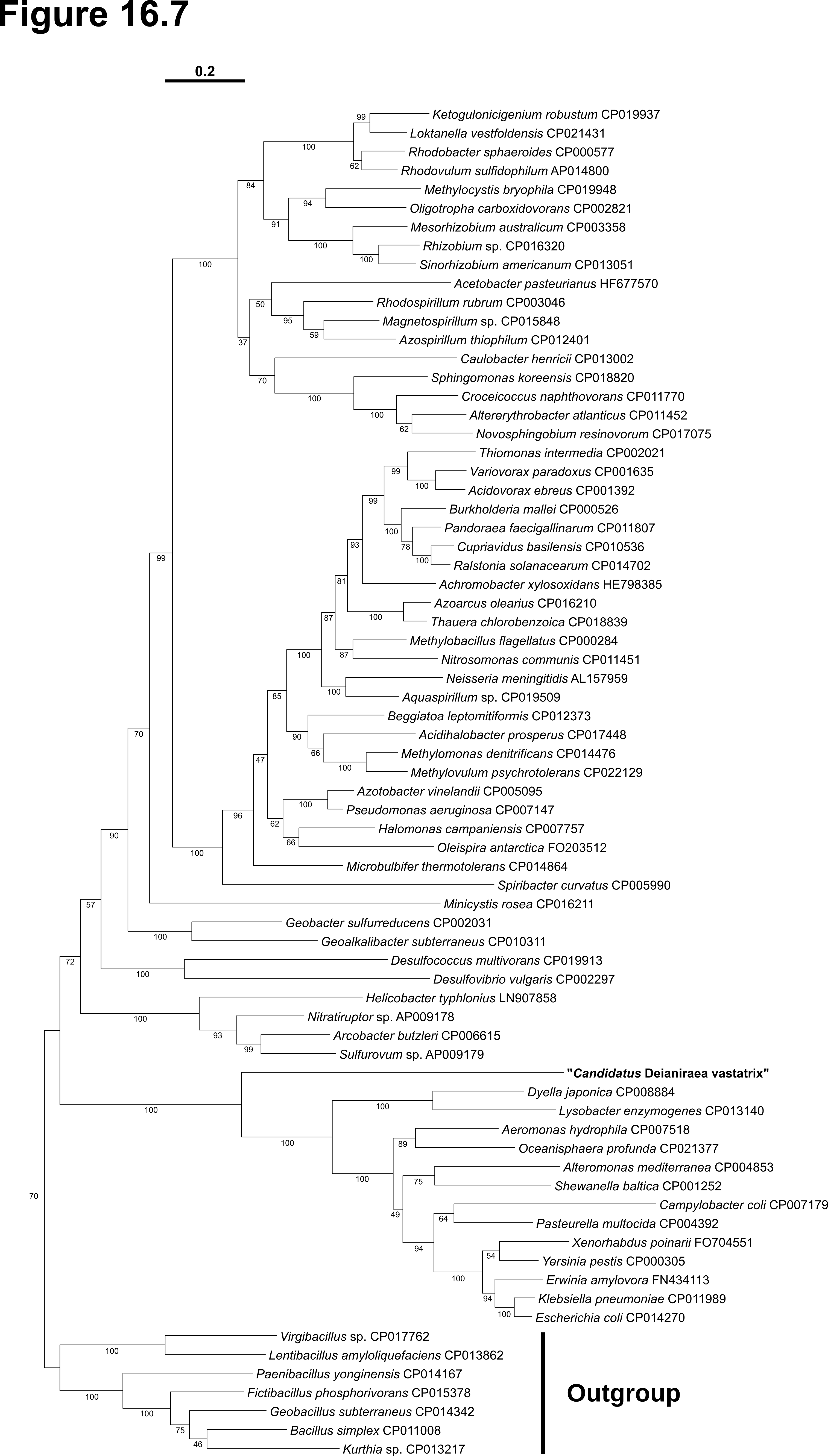

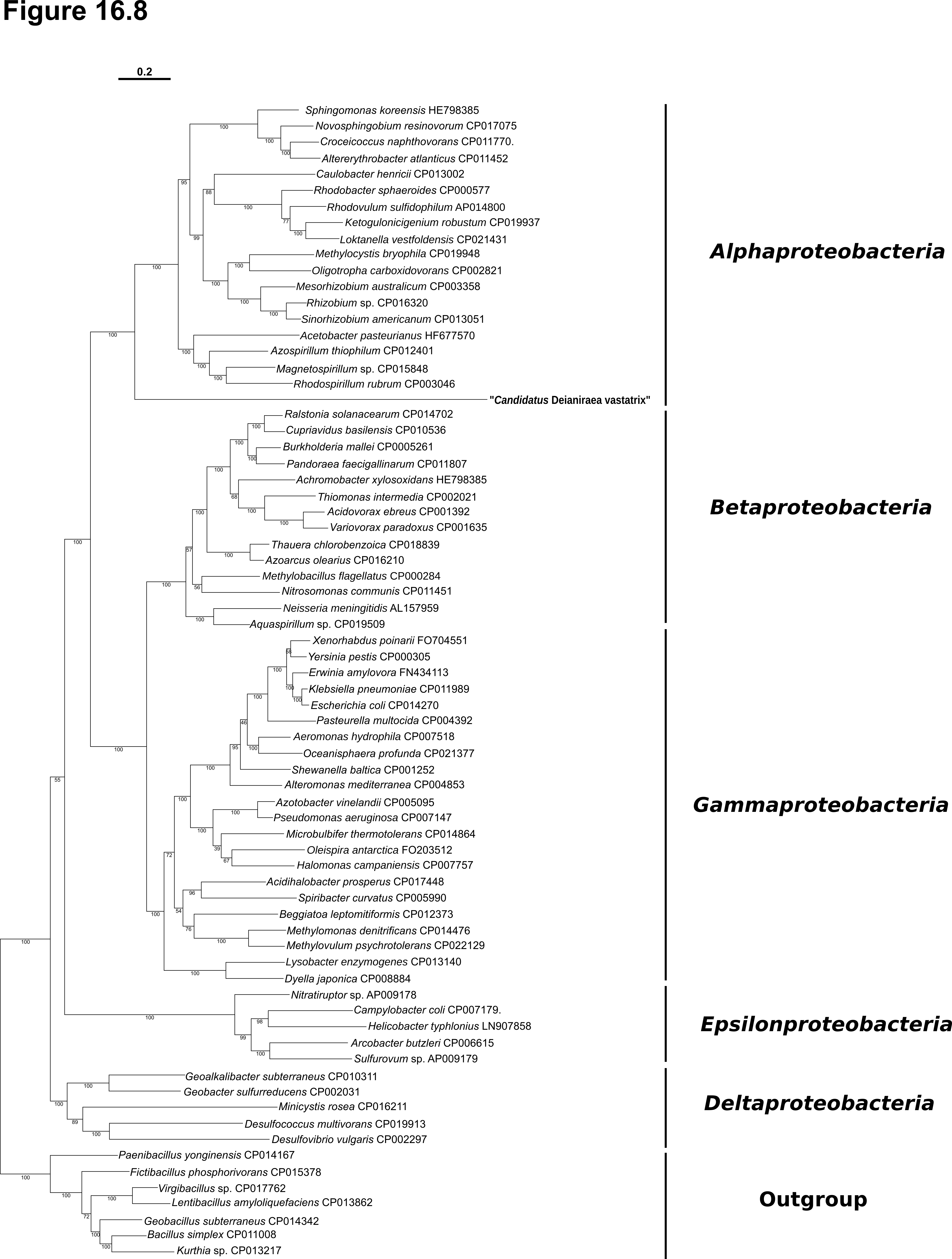

**Methionine synthesis and metabolism**

In bacteria (including *Deianiraea*), three enzymatic steps preliminary to the synthesis of methionine lead from aspartate to homoserine (Bender 2012). The first difference of *Deianiraea* respect to other *Rickettsiales* resides in the last step of this pathway, catalysed by the homoserine dehydrogenase. This enzymatic activity is indeed absent in all other *Rickettsiales* except “*Ca*. Arcanobacter lacustris” (Martijn et al. 2015), in which homoserine is likely the precursor of threonine, which cannot be synthesised by any other *Rickettsiales* (including *Deianiraea*). In *Deianiraea*, the homoserine dehydrogenase activity is fused to aspartate kinase (catalysing the first step of homoserine synthesis), similarly to many *Gammaproteobacteria* and *Deltaproteobacteria*, and differently respect to most *Alphaproteobacteria*, possessing separate genes for the two enzymatic activities.

The bacterial pathway specific for methionine biosynthesis from homoserine involves three to four steps (Ferla and Patrick 2014; Bender 2012). The first step is always the activation of homoserine, mostly through acylation from a CoA carrier. This reaction can be catalysed alternatively catalysed by distinct enzymes (MetA or MetX), depending on the phylogenetic lineage. While among *Alphaproteobacteria* the two genes are almost equally abundant (Ferla and Patrick 2014), *Deianiraea* lacks both. Considering that it possesses the enzymatic potential for the following steps, most likely it is able to circumvent such apparent deficiency either by direct import of activated homoserine, or by producing it autonomously through a different reaction. Interestingly, according to blastp search, another ORF (Deia_00649) finds some homology to proteins annotated as “Methionine biosynthesis protein MetW”, mostly belonging to other *Alphaproteobacteria* (Supplementary material 19). This protein was suggested to be involved in the acylation of homoserine, possibly in cooperation with MetX (Andersen et al. 1998; Alaminos and Ramos 2001). Thus, ORF Deia_00649 was provisionally included in the *Deianiraea* gene set for methionine biosynthesis, while further analyses would be obviously necessary to clarify this point.

Further synthetic steps involve the sulfurylation of the activated homoserine, to produce homocysteine. Two alternative strategies are employed by bacteria, either a two-step trans-sulfurylation route, catalysed by cystathionine gamma-synthase (MetB) and cystathionine beta-lyase (MetC) using cysteine a sulfhydryl group donor, or a direct sulfurylation from free hydrogen sulfide, catalysed by one among two possible O-acetylhomoserine thiolases (MetY or MetZ). The four proteins (MetB, MetC, MetY, MetZ) are all homologous, and the apparently redundant presence of multiple alternative pathways in the same bacterium is frequent. Two complete genes of belonging to this homology group were identified in *Deianiraea* (Deia_00032, Deia_00704), plus an additional much shorter ORF (Deia_00017), which was thus not considered in the analysis. According to sequence identity, the complete genes were putatively assigned to bacterial MetC and MetY, respectively. Both of these forms are common in *Alphaproteobacteria*. The presence of MetC gene in absence of MetB might be explained by hypothesising a possible bifunctional activity, involving also the gamma-synthase activity (Ferla and Patrick 2014). Among *Rickettsiales*, putative MetC homologues were found also in some *Anaplasmataceae*, in particular several *Wolbachia* (e.g. Foster et al. 2005), “*Ca*. Neoehrlichia lotoris” (Daugherty, S.C et al. unpublished: [LANX01000001]), and few *Anaplasma* species (*A. marginale* and *A. centrale*) (Dark et al. 2009; Herndon et al. 2010).

The last biosynthetic step is the methylation of homocysteine, to obtain methionine. Two distinct homocysteine transmethylases can catalyse this reaction, and the two forms might coexist in the same organism. *Deianiraea* has two different ORFs (Deia_00662, Deia_00705) belonging to the same homology group, the cobalamin-independent homocysteine transmethylase MetE (methyl group donor is 5-methyltetrahydropteroyl-tri-L-glutamate), differently than several other *Alphaproteobacteria*, which more frequently display the cobalamin-dependent form.

Another ORF (Deia_00702), encoding the 5,10-methylenetetrahydrofolate reductase MetF, was taken into account in the analysis. Though not directly involved in methionine biosynthesis, it is as well related to methionine metabolism, and it is absent in all other *Rickettsiales*, thus it was considered putatively relevant, and was included in the analysis.

**Phylogenetic analyses**: Used three genes and 70 organisms, 1,071 sites. For the phylogenomics, used 53 ortholog genes and 9,246 sites.

**Figure 16.9** Maximum likelihood tree of the three concatenated genes involved in the methionine biosynthesis with the LG+I+G+F substitution model with 100 bootstrap pseudo-replicates. The position of *Deianiraea* genes is unsupported. Scale bar stands for estimated sequence divergence. Number on branches stand for bootstrap values.

**Figure 16.10** Reference maximum likelihood phylogenomic tree of the organisms employed for phylogenesis in **16.9** with the LG+I+G substitution model with 100 bootstrap pseudo-replicates. The five proteobacterial classes (as indicated in the figure) are all monophyletic and highly supported, including the positioning of *Deianiraea* within *Alphaproteobacteria*, which is consistent with the analyses shown **Supplementary material 11**. Scale bar stands for estimated sequence divergence. Number on branches stand for bootstrap values.

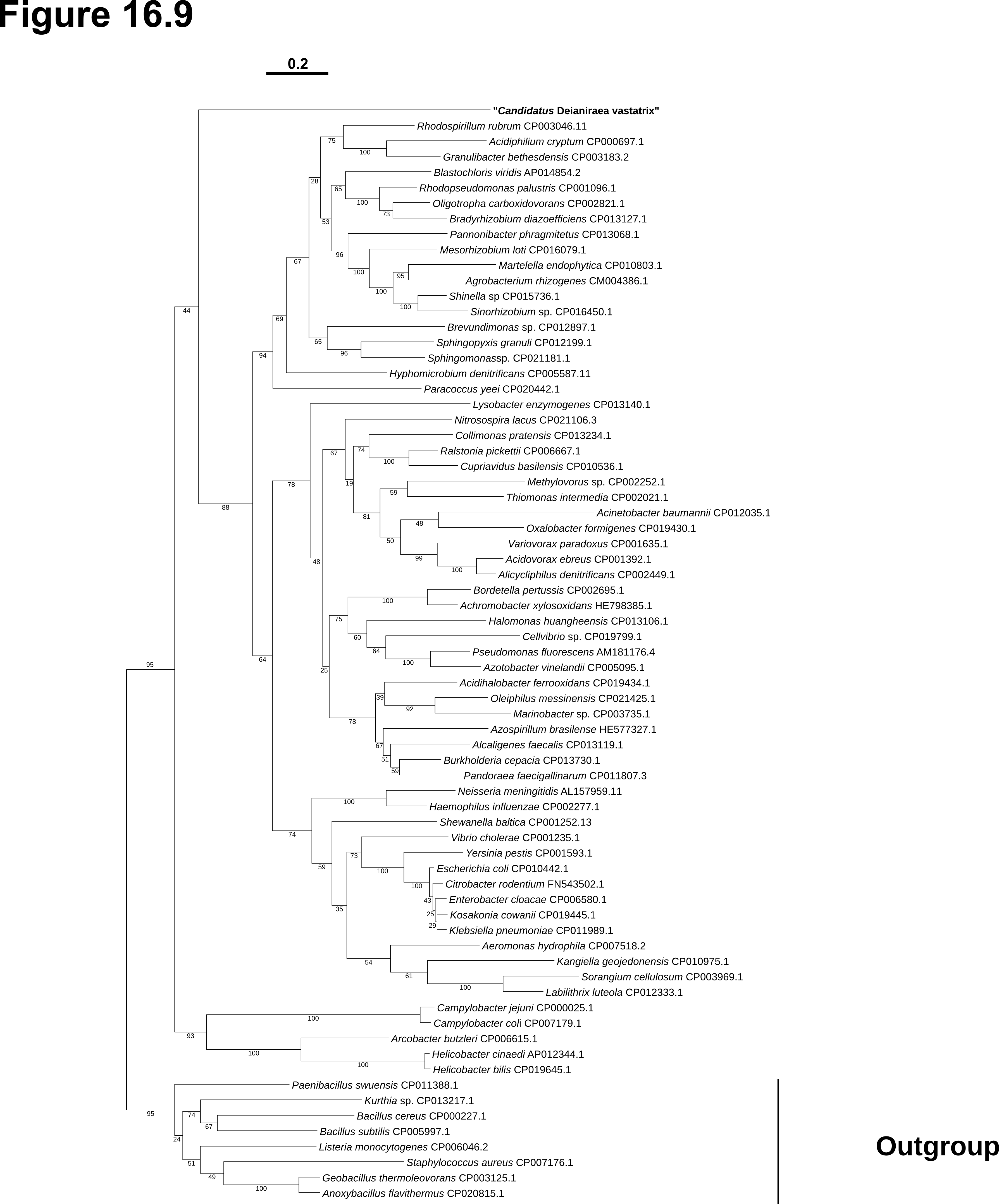

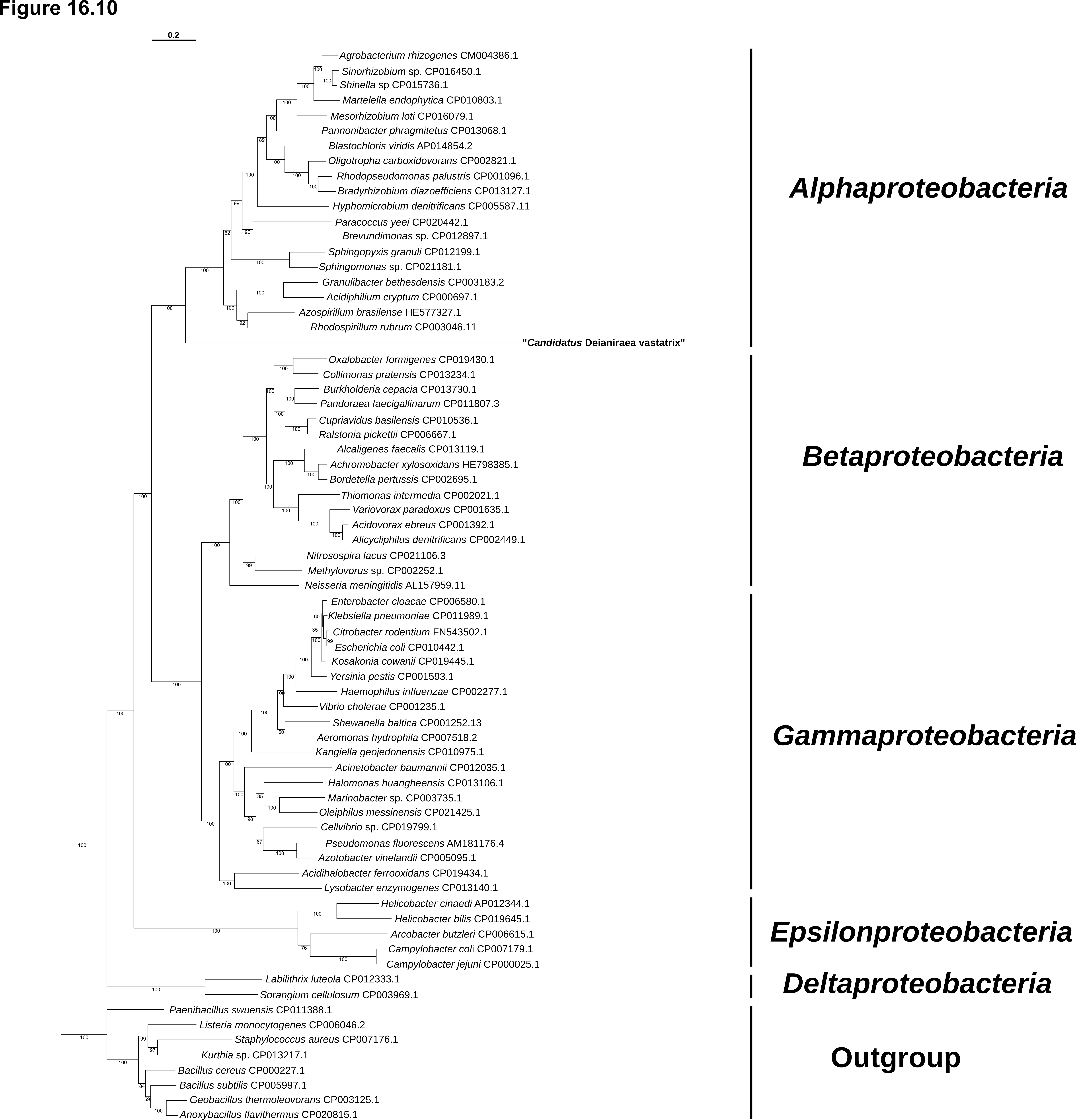

**Conclusions**

Summing up, *Deianiraea* possesses the distinctive ability to synthesise eight amino acids more than any other known *Rickettsiales* bacterium. The respective biosynthetic enzymes can be grouped into five pathways. For each pathway, no evidence of recent HGT was found. Indeed, all the genes are highly divergent from any other organism (mostly <50% identity), and no relevant trend of higher similarity to very distantly related organisms was observed, since the identity values respect to best hits on the entire nr protein database, on *Proteobacteria*, and *Alphaproteobacteria* (which would be the closest relatives in the absence of HGT) are all comparable (Supplementary material 19). Moreover, for no pathway the combined result of phylogenetic analyses and GC content and CAI deviation tests showed a clear evidence of recent HGT. For what concerns phylogenetic analyses, in most cases, the whole pathway-specific dataset*,* lacked a sufficient phylogenetic signal, as confirmed by the highly significant deviations respect to organismal phylogenies in SH tests. Thus, it was not possible to reconstruct confidently the position of *Deianiraea* genes. Nevertheless, given that they were always found with a very long branch (consistent with sequence divergence), and that they were never found associated to a recent (family level or lower) group of bacteria, the conclusion that those genes did not undergo any recent HGT was indirectly confirmed. Therefore, the hypothesis that they could be inherited directly from *Rickettsiales* ancestor resulted fortified. Based on these lines of reasoning, further hypotheses on the evolutionary trends of *Deianiraea* and *Rickettsiales* in general were fittingly drawn.

Daugherty, S. C. *et al.* unpublished: [LANX01000001]

Emms, D. M. & Kelly, S. OrthoFinder: solving fundamental biases in whole genome comparisons dramatically improves orthogroup inference accuracy. *Genome Biol.* 16(1): 157.

Felsheim, R. F. *et al.* unpublished. [NZ_CP009217])

Ferla, M. P. & Patrick, W. M. Bacterial methionine biosynthesis. *Microbiology* **60**, 1571-1584 (2014)

Foster, J., *et al.* The *Wolbachia* genome of *Brugia* *malayi*: endosymbiont evolution within a human pathogenic nematode *PLoS Biol.* **3**, e121 (2005)

Herndon, D. R., Palmer, G. H., Shkap, V., Knowles, D. P. Jr. & Brayton,K.A. Complete genome sequence of *Anaplasma* *marginale* subsp. *centrale.* *Bacteriol*. **192**, 379-380 (2010)

Jima, D. D. *et al.* Whole-genome sequence of "*Candidatus* Rickettsia asemboensis" strain NMRCii, isolated from fleas of western Kenya. *Genome Announc.* **3**, pii: e00018-15 (2015)

Kwan, J.C. & Schmidt, E.W. Bacterial endosymbiosis in a chordate host: long-term co-evolution and conservation of secondary metabolism. *PLoS One* **8**, e80822 (2013)

Lawson, C. E. *et al.* Metabolic network analysis reveals microbial community interactions in anammox granules. *Nat. Comm.* **8**, 15416 (2017)

Makino, Y. *et al.* An archaeal ADP-dependent serine kinase involved in cysteine biosynthesis and serine metabolism *Nat. Comm.* **7**, 13446 (2016)

Marchler-Bauer, A. *et al.* CDD: NCBI's conserved domain database. *Nucleic Acids Res.* **43**, D222-D226 (2015)

Martijn, J. *et al.* Single-cell genomics of a rare environmental alphaproteobacterium provides unique insights into *Rickettsiaceae* evolution. *ISME J.* **9**, 2373-2385 (2015)

McKinlay, J. B. & Harwood, C. S. Carbon dioxide fixation as a central redox cofactor recycling mechanism in bacteria. *Proc. Nat. Acad. Sci. USA* **107**, 11669–11675 (2010)

Mediannikov, O. *et al.* High quality draft genome sequence and description of *Occidentia* *massiliensis* gen. nov., sp. nov., a new member of the family *Rickettsiaceae*. *Stand. Genomic Sci.* **9**, 9 (2014)

Pittard, J. & Yang, J. Biosynthesis of the aromatic amino acids. *EcoSal Plus* **3**, (2008) doi:10.1128/ecosalplus.3.6.1.8.

Probst, A. J. *et al.* Depth-based differentiation of microbial function through sediment-hosted aquifers and enrichment of novel symbionts in the deep terrestrial subsurface. Nat. Microbiol. **3**, 328-336 (2018)

Risso, C., Van Dien, S. J., Orloff, A., Lovley, D. R. & Coppi, M. V. Elucidation of an alternate isoleucine biosynthesis pathway in *Geobacter sulfurreducens*. *J. Bacteriol.* **190**, 2266–2274 (2008)

Salmon, K. A., Yang, C. R. & Hatfield, G. W. Biosynthesis and regulation of the branched-chain amino acids. *EcoSal Plus* **2** (2006); doi:10.1128/ecosalplus.3.6.1.5.

Tang, K. H., Feng, X., Tang, Y.J. &Blankenship, R. E. Carbohydrate metabolism and carbon fixation in *Roseobacter denitrificans* och114. *PLoS One* **4**, e7233 (2009)

Tully, B. J., Sachdeva, R., Graham, E. D. & Heidelberg, J. F. 290 metagenome-assembled genomes from the Mediterranean Sea: a resource for marine microbiology. *PeerJ.* **5**, e3558 (2017)

Winkler, M.E. & Ramos-Montañez, S. Biosynthesis of histidine. *EcoSal Plus* **3** (2009) doi: 10.1128/ecosalplus.3.6.1.9.

Xu, H. *et al.* Isoleucine biosynthesis in *Leptospira interrogans* serotype lai strain 56601 proceeds via a threonine-independent pathway. *J. Bact.* **186**, 5400-5409 (2004)
