## Supplementary material for "The extracellular association of the bacterium “*Candidatus* Deianiraea vastatrix” with the ciliate *Paramecium* suggests an alternative scenario for the evolution of *Rickettsiales*": Suppl_26_assembly_procedures.docx

**Genome sequencing and assembly of *Deianiraea vastatrix***

Initially, total genomic DNA was extracted with Nucleospin Plant II kit from a sample of ~100 *Paramecium* cells covered by *Deianiraea* and fixed in ethanol, using the protocol for mycelium. Thus, in order to reach an appropriate DNA quantity for sequencing, the extract was subjected to whole-genome amplification (WGA) with the REPLI-g Single Cell Kit (Qiagen), following the instructions for DNA extracts.

The WGA product was processed through a Nextera XT library, and sequenced on an Illumina HiSeq X by Admera Health LC (South Plainfield, NJ, USA), producing 37,100,964 pairs of 150 bp reads. The read quality was preliminarily assessed with FastQC (Andrews 2010).

The total reads were processed using SPAdes 3.6 (Bankevich et al. 2012) with default settings, obtaining a “preliminary assembly” (50,557,437 bp; 76,183 contigs; N50= 1,281 bp; L50= 8,830). Then, a multi-step procedure was applied, in order to select only those contigs belonging to *Deianiraea* and discard those belonging to *Paramecium* and to free-living bacteria present in traces in the sample. These bacteria included *Enterobacter aerogenes*, provided as food for *Paramecium*, and other bacteria associated to the culture and presumably derived from the original sample.

For this purpose, the blobology pipeline was applied (Kumar et al. 2013). Basically, at first contigs were classified according to their length, GC% content, sequencing coverage (according to reads mapping with Bowtie2 (Langmead and Salzberg 2012)), and NCBI taxonomy of the best megablast hit on NCBI nucleotide.

Aftewards, SSU rRNA genes in the “preliminary assembly” were predicted with barrnap (Seemann 2013), identifying 78 partial genes, and no full-length gene (Supplementary material 27)

- 39 partial SSU rRNA genes had best blast hits on nuclear or mitochondrial *Paramecium* spp. SSU rRNA genes. Considering that members of the *Paramecium aurelia* species complex are almost identical in the SSU rRNA genes and can be discriminated only by more variable genetic markers (e.g. Catania et al. 2009), all the respective contigs were assigned to the *P. primaurelia* host
- 30 partial SSU rRNA genes were almost identical (>98%) to the 16S rRNA gene of *Deianiraea* *vastatrix* as previously determined through PCR and Sanger sequencing. Taking into account minor sequencing/assembly errors, we assigned the respective contigs to *Deianiraea*. Those contigs ranged from 181.35 to 7583.17 in sequencing coverage and they all had best megablast hits with *Bacteria*
- 8 partial SSU rRNA gene sequences blasted on other bacteria. Most of the respective contigs (7), displayed comparably lower coverage (≤86.92)
- 1 partial SSU sequence had best-hit with a fungus and its contig coverage was 11.95

Taking into account the SSU rRNA features, the “initial selection from preliminary assembly” for *Deianiraea* was designed, including all the contigs with coverage higher than 100, best blast hit with *Bacteria* or no hit (to avoid false negatives, accounting the evolutionary distance respect to previously sequenced bacteria), minimal length of 250 bp, and no restriction on GC content (7,362 contigs, 4,551,029 bp; Supplementary material 28)

The reads mapped to “initial selection from preliminary assembly” were extracted and reassembled separately with SPAdes default settings: “first re-assembly – Bacteria and not annotated” (1,441,196 bp, 325 contigs, N50=192630 bp; L50=3; Figure A). Reciprocal and self blastn of the contig selection from the preliminary assembly and the respective reassembly were performed, to investigate the apparently high difference in total length. This was verified to be due to the high degree of fragmentation of the preliminary assembly, containing many short, partially overlapping contigs, which were thus redundant and resulting in a large over-estimate of the lengths of the actual genome sequences (data not shown). The first reassembly contained, according to barrnap, the full-length SSU rRNA gene of *Deianiraea* on a large contig (37706 bp), plus only three partial SSU rRNA genes from other bacteria and the host mitochondrion, all in short contigs (<1000bp) (Supplementary material 29).

After such identification of a first putative *Deianiraea* contig, other putative contigs were added using the assembly-derived connections between contig ends, according to the contigs.fastg SPAdes output, inspected with the Bandage software (Wick et al. 2015). Each assignment was verified by careful manual inspection of all the contigs of this assembly, examining their blastn results, the automatic annotation by prokka (Seemann 2014), and blastp results of representative predicted CDSs for each contig. This procedure allowed to discard a total of 299 contigs (238,190 bp), in detail: 14 assigned to *Paramecium* mitochondrion, 34 to *Paramecium* macronucleus; 68 to other bacteria, and 183 unassigned. These last 183 were short (<4000 bp) and unconnected in the assembly graph, and likely represented minor sequencing/assembly errors and/or negligible traces of other genomes (Figure A). The remaining 26 contigs (1,203,006 bp: 83.5% of this assembly) were robustly assigned to *Deianiraea* and represented the trusted set for following analyses.

**Figure A.** Scheme of the contigs (coloured bars) of the “initial selection from preliminary assembly” and their putative end-connections in the assembly graph (gray lines), obtained with Bandage. Coloured rectangular shapes indicate the organism(s) to which the internal contigs were assigned, their number, total length, and length percentage on the total assembly. The *Deianiraea* selection represented the trusted contigs for successive assembly refineniments. Due to space limitation, only for this selection contig names (numbering is from the longest to the shortest in the total assembly) and length are shown.
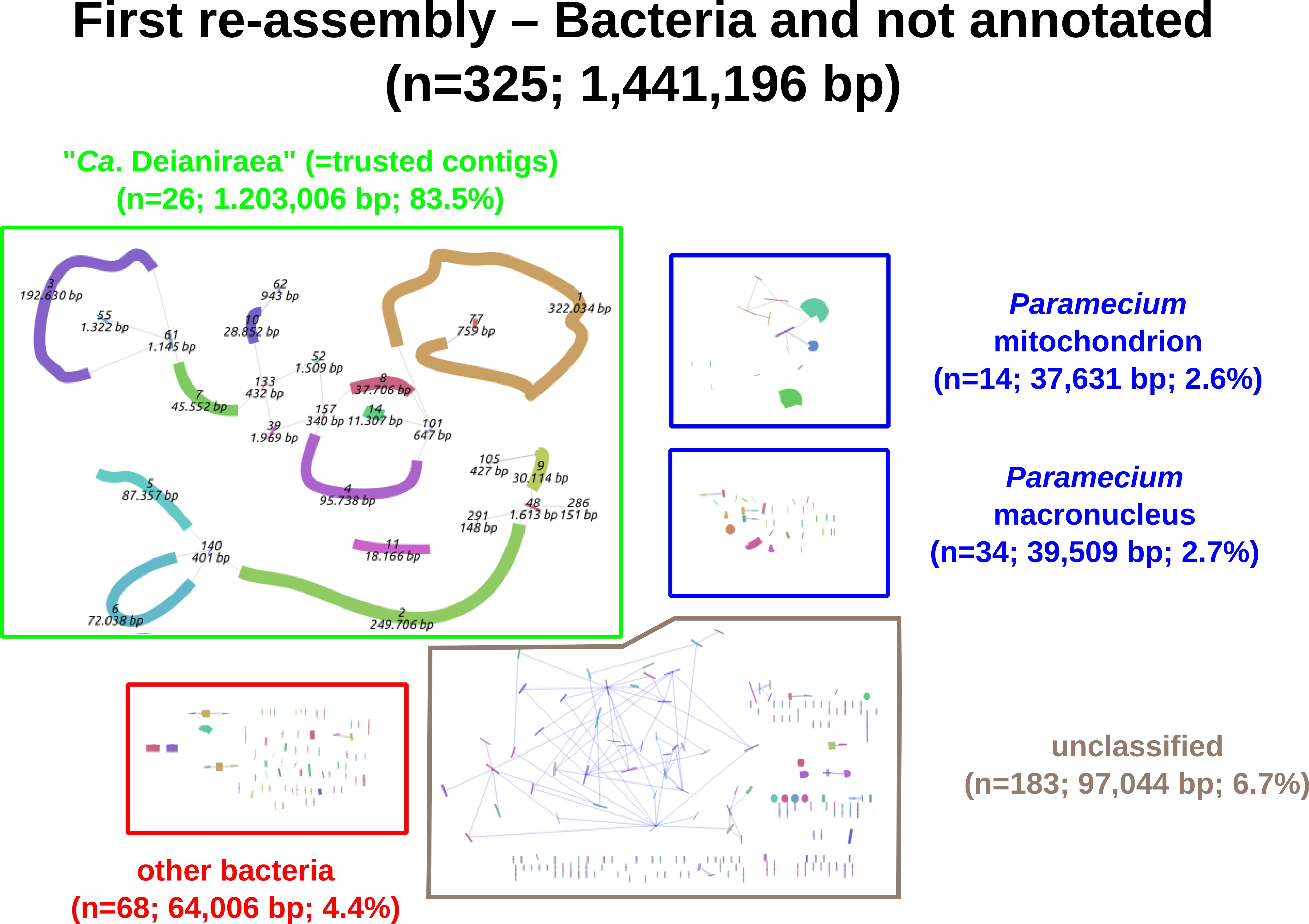

Thereafter, a step-wise revision and improvement of the assembly was performed by fishing false-negative contigs of the preliminary assembly, by the above-described blastn- and annotation-based procedure, as well as also by repiprocal-blast with trusted contigs, seeking for overlapping ends. In this way, two putative “*Ca*. Deianiraea”-contigs from the preliminary assembly were identified (NODE_492, NODE_7016), and corresponding reads were reassembled in an intermediate reassembly attempt (data not shown). At this stage, primer design for PCR started (see end of the present supplementary file). A single walking PCR reaction (Supplementary material 30; Pilhofer et al. 2007) allowed to identify another contig from the preliminary assembly (NODE_697), which resulted to be neighboring a contig of the intermediate reassembly. Thus, these combined approaches allowed to recover in total 3 further contigs from the preliminary assembly (all with coverage>100), which were added to the “initial selection from preliminary assembly” (= “final selection from preliminary assembly”: 7,365 contigs, 4,551,029 bp, Supplementary material 28). Interestingly, these three had best-BLAST-hit to eukaryotes (low scores), and for this reason had been initially wrongly discarded.

The reads mapped to this “final selection from preliminary assembly” were assembled with SPAdes. To optimize the result, multiple values of “cov_cutoff” at 100 intervals in the range 100-2000 were tested (using the trusted contigs as reference), finally selecting 1500 as optimal, which allowed to discard automatically many lower coverage non-“*Ca*. Deianiraea” sequences. Thus, the “final SPAdes assembly” was obtained (60 contigs, 1,250,290 bp; N50=323,702 bp, L50=2; Figure B), containing a single full length SSU rRNA, belonging to “*Ca*. Deianiraea” (100% identity with PCR sequence). The manual inspection of “final SPAdes assembly” allowed to further discard 48 contigs (4 assigned to *Paramecium* mitochondrion, 7 to other bacteria, and 37 short (≤1928 bp) unassigned ones). Hence, a “final bioinformatic assembly” was obtained, constituting ~96.2% of the length of “final SPAdes assembly” (1,202,853 bp; 12 contigs; N50=323,702 bp; L50=2).

**Figure B.** Scheme of the contigs (coloured bars) of the “final SPAdes assembly” and their putative end-connections in the assembly graph (gray lines), obtained with Bandage. Contig names (numbering is from the longest to the shortest), and length are shown. Coloured rectangular shapes indicate the organism(s) to which the internal contigs were assigned, and the length percentage on the total assembly.
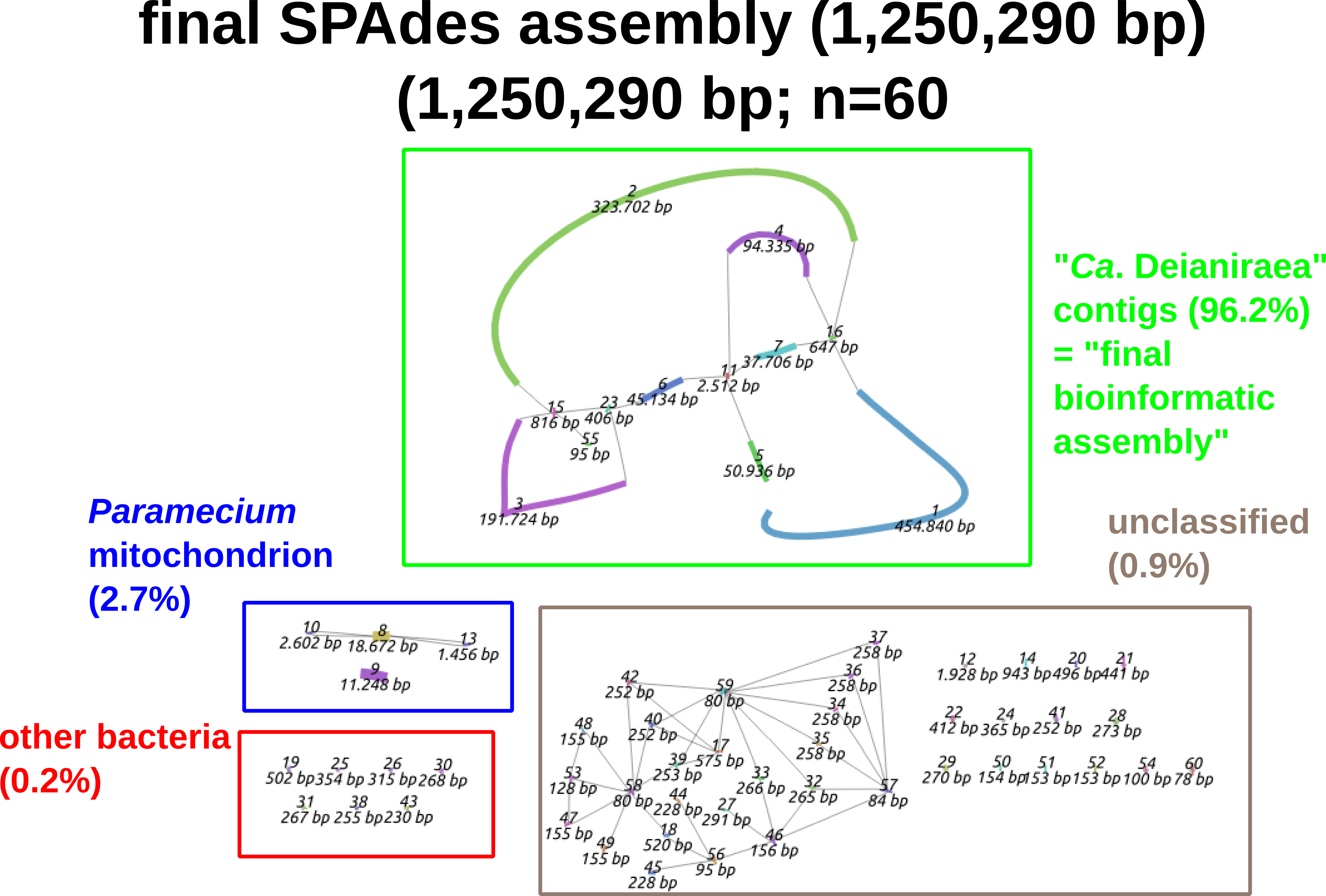

In order to fully join the contigs of the “final bioinformatic assembly”, a PCR-based approach was exploited (Details on primers and PCR cycles in Supplementary material 30). Outwardly directed primer were designed in proximity to the ends of the 7 major contigs. The combinations of contig ends, and thus primers, to be tested for PCR reactions were selected based on the assembly-derived connections, according to the contigs.fastg output, inspected with Bandage (Figure B). Thus, 7 contig junctions were assessed and confirmed by conventional PCR. In addition, during the refinement of the selection of contigs from the preliminary assembly, a two-step walking PCR (Pilhofer et al. 2007) allowed to recover a preliminary contig (Supplementary material 30), for which the sequence resulted to be neighboring the one of a contig from an intermediate reassembly step (this junction resulted internal to NODE_1 of the “Final SPAdes assembly”).

Thus, all contigs were manually joined into a single circular chromosome (sized 1,205,153 bp), representing the final assembly of *Deianiraea vastatrix*.
