## Supplementary figures and images for "The extracellular association of the bacterium “*Candidatus* Deianiraea vastatrix” with the ciliate *Paramecium* suggests an alternative scenario for the evolution of *Rickettsiales*"

### Suppl_1_healthy_DIC_AFM_svg.png

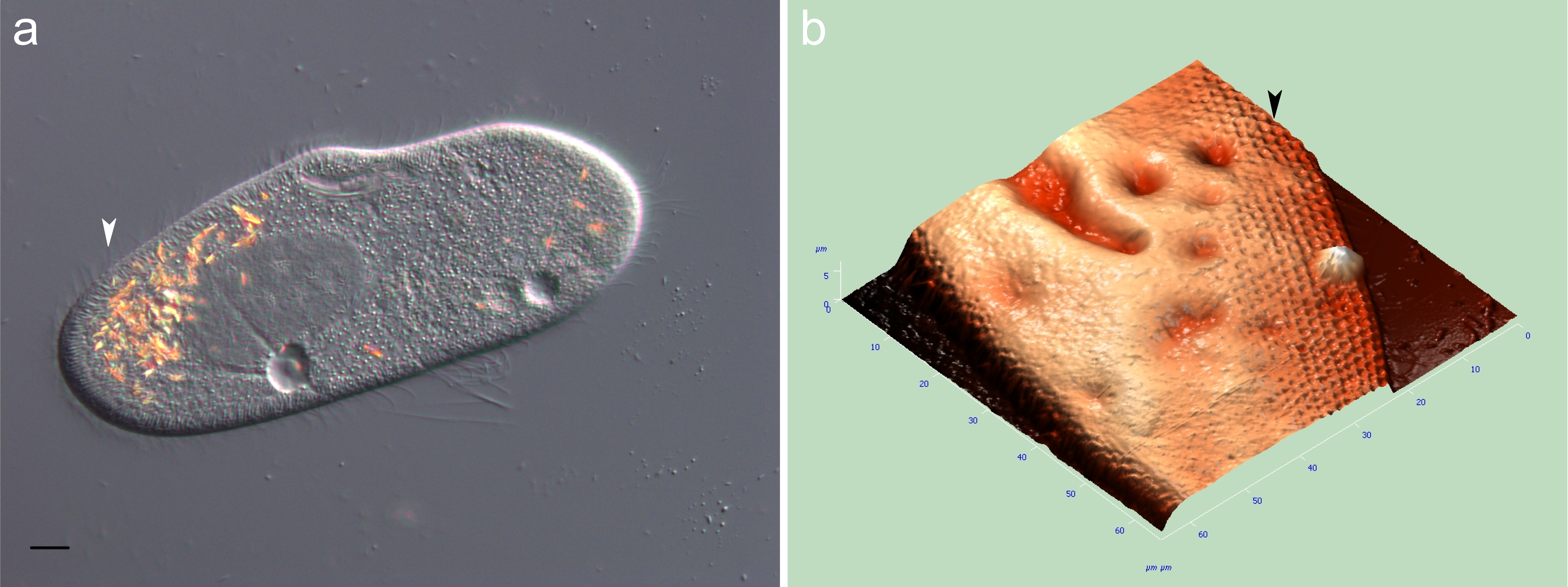

### Suppl_2_TEM_and_AFM.tif2.png

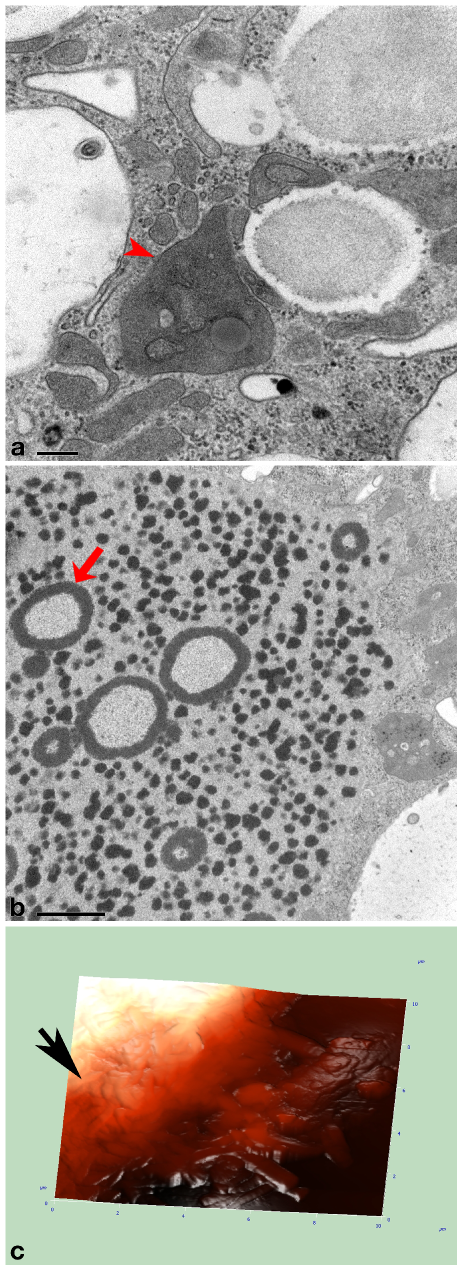

### Suppl_4_FISH.tif

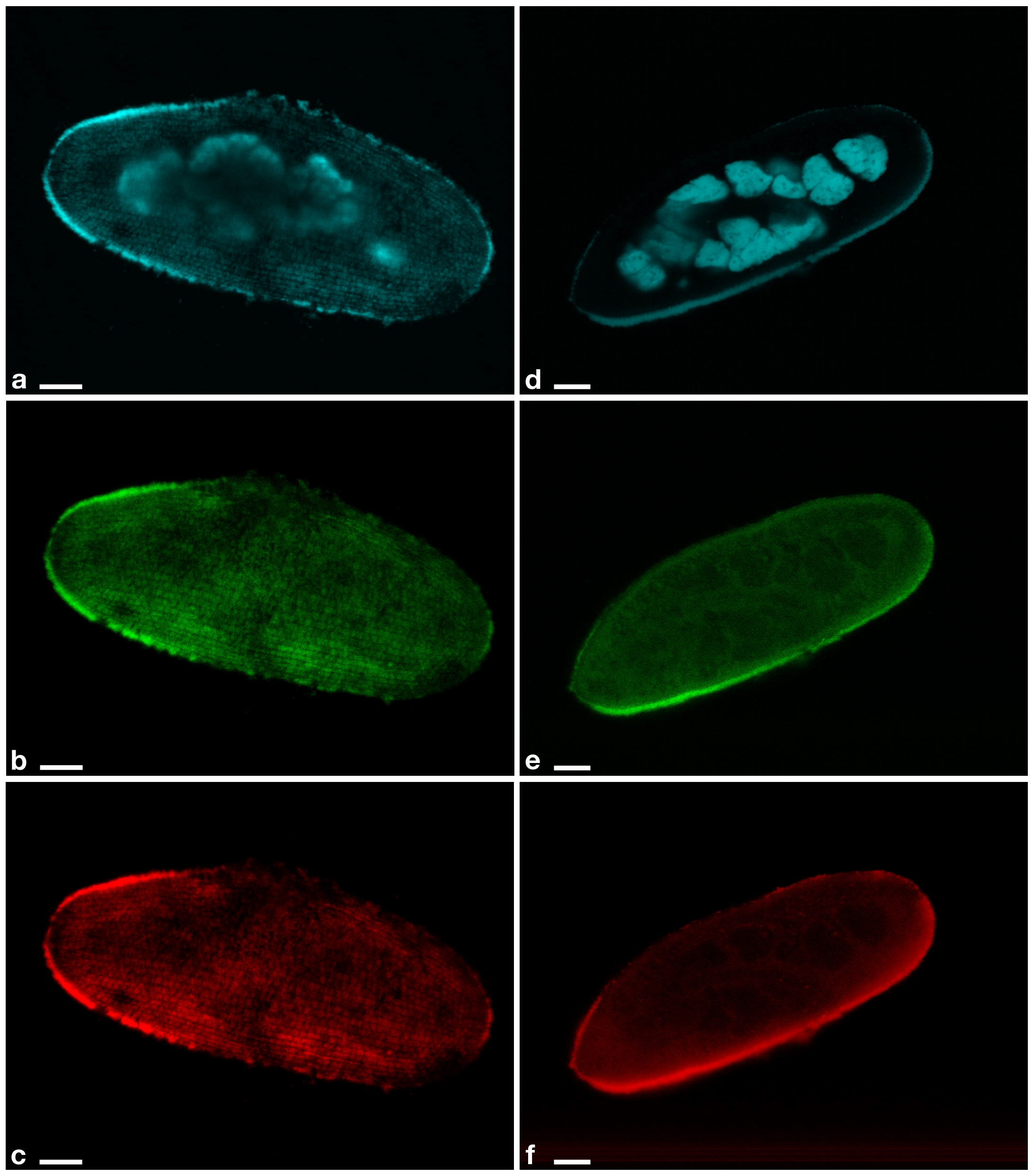

### Suppl_5_ML_and_BI_16S_merged.pdf

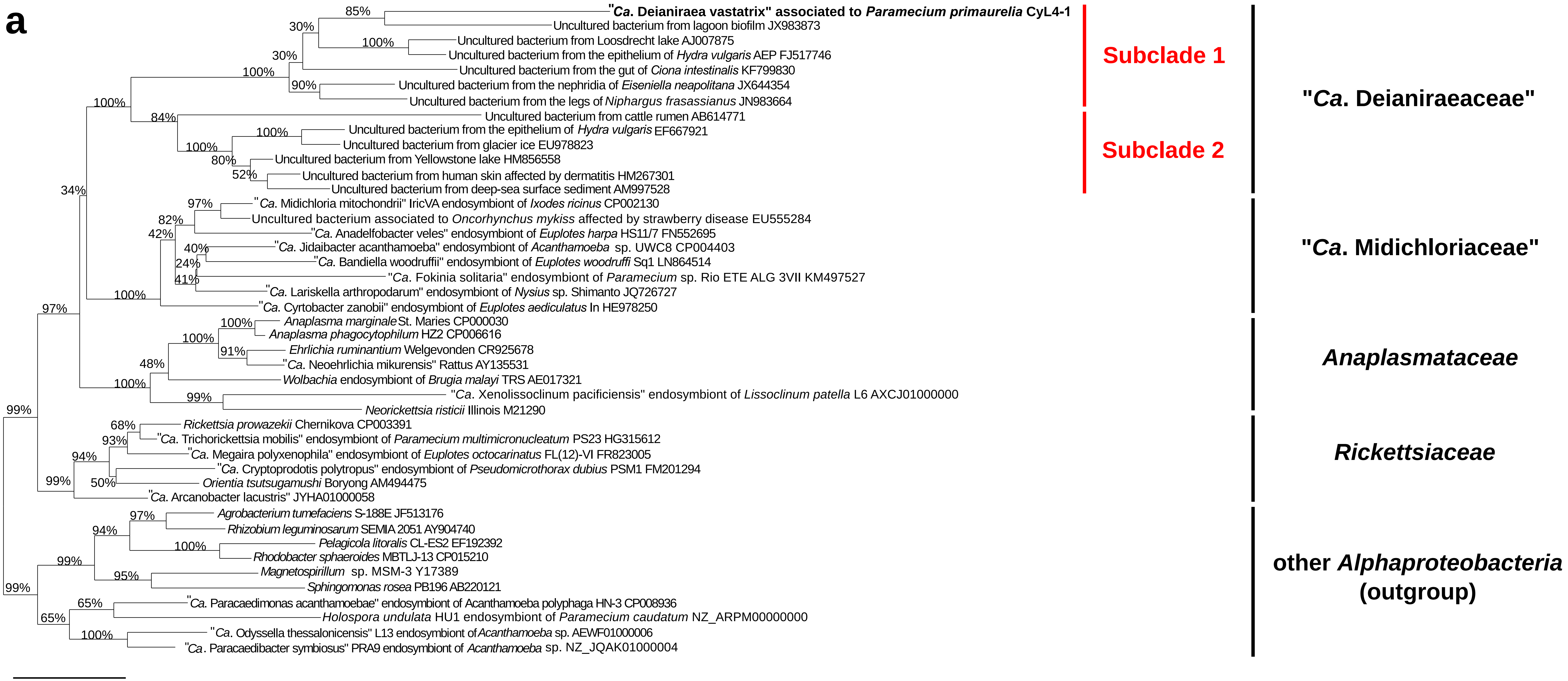

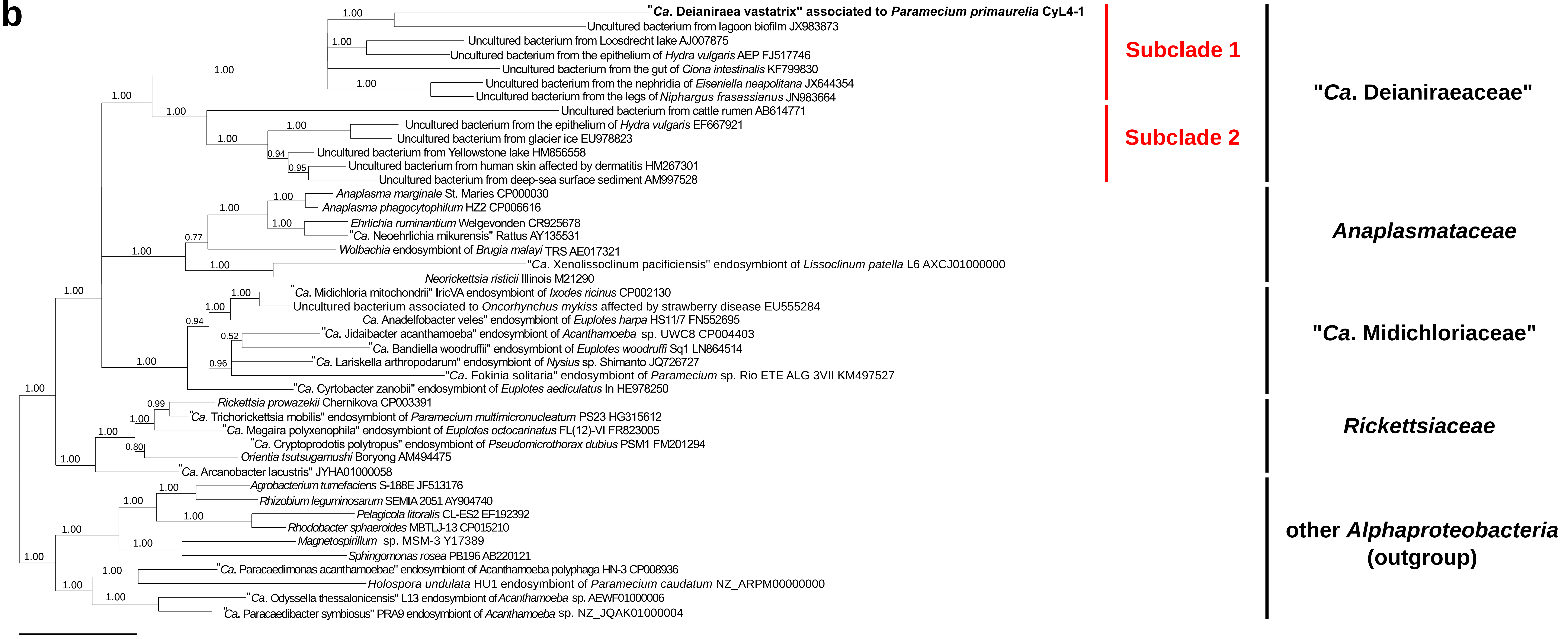

### Suppl_11_Phylogenomics_gene_set1.pdf

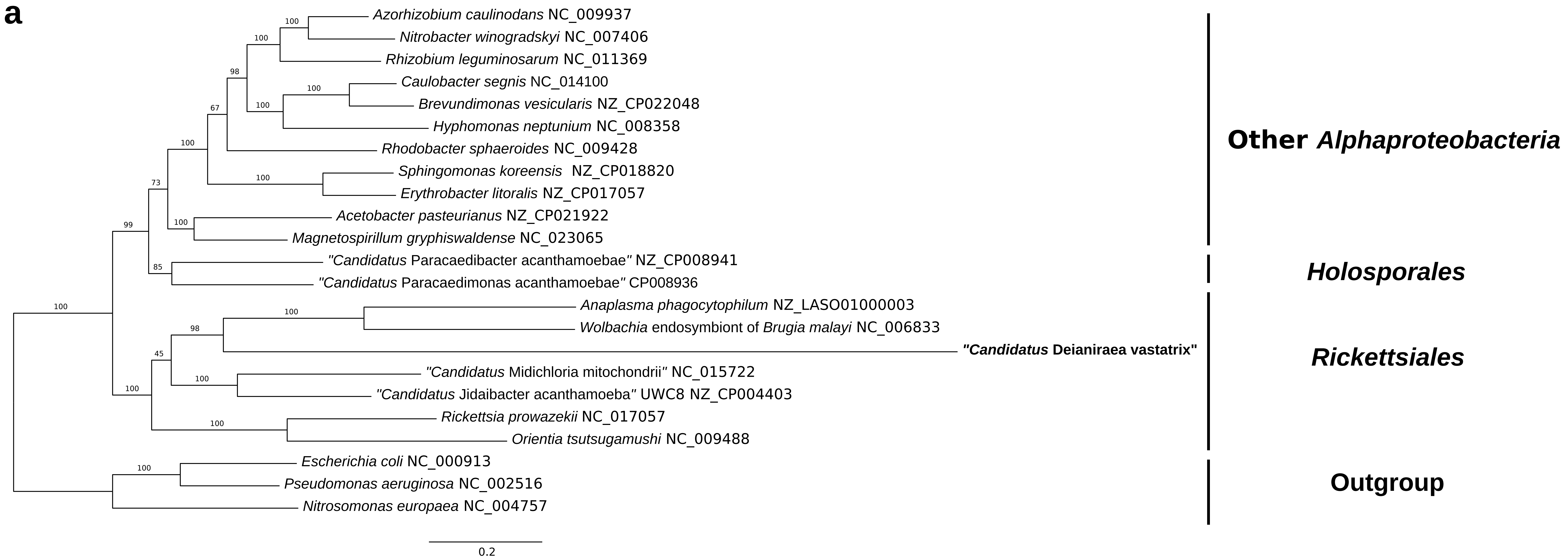

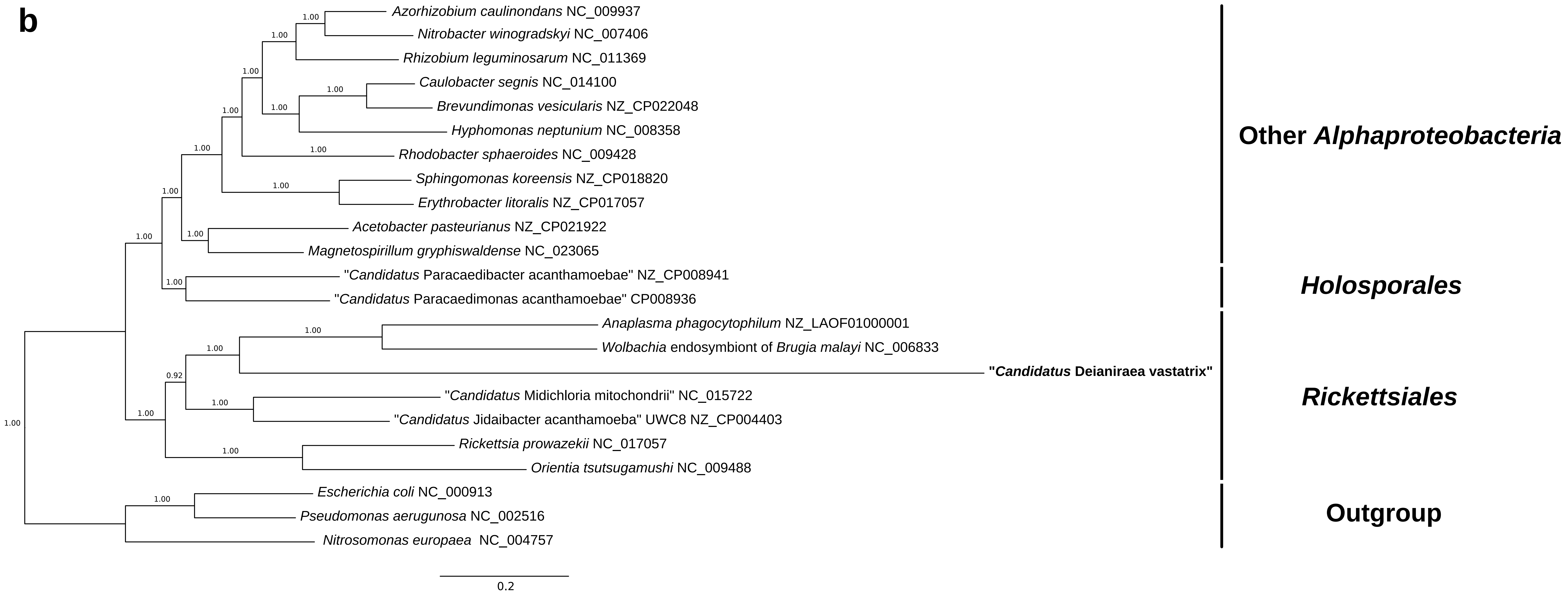

### Suppl_12_Phylogenomics_gene_set2.pdf

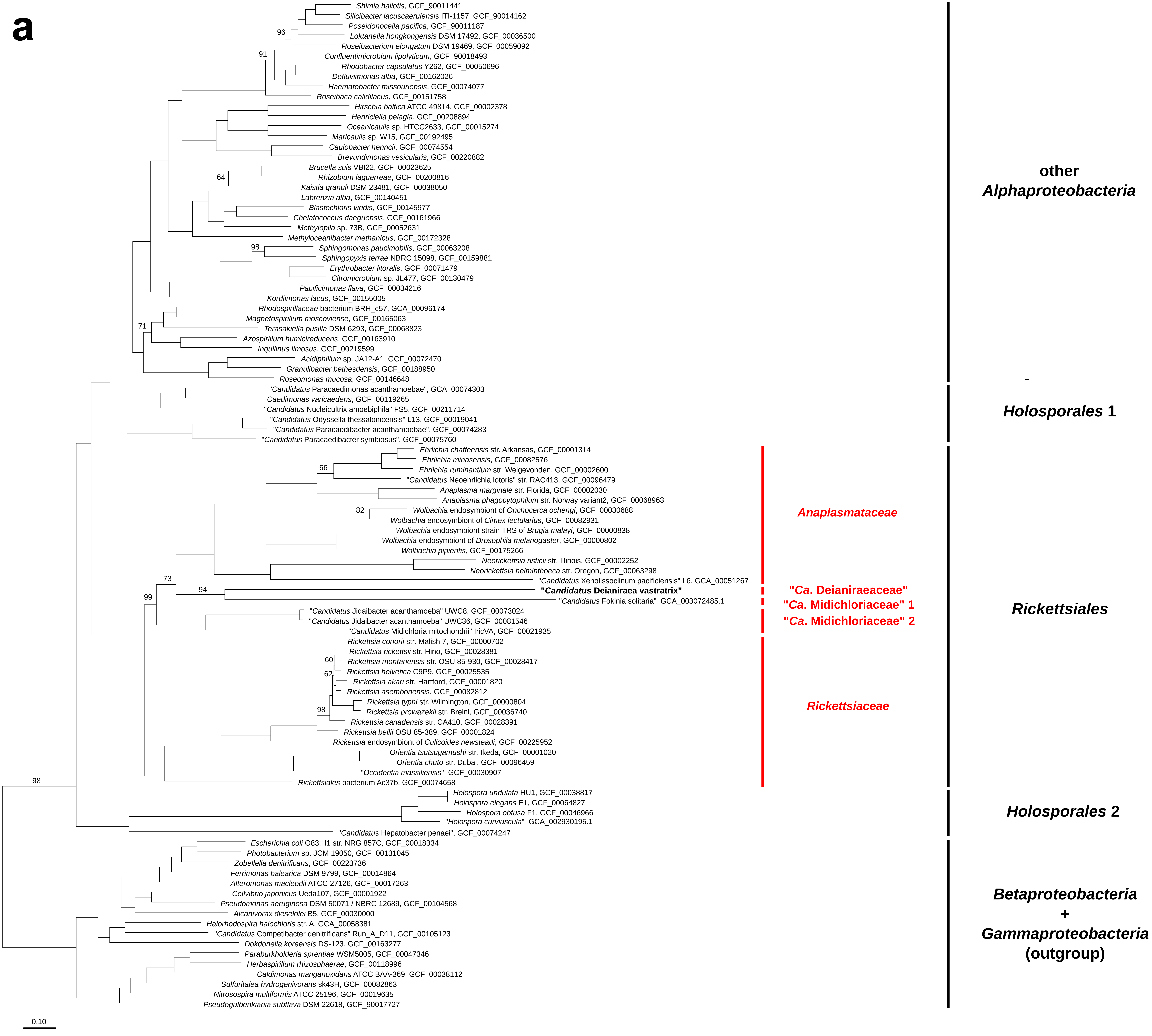

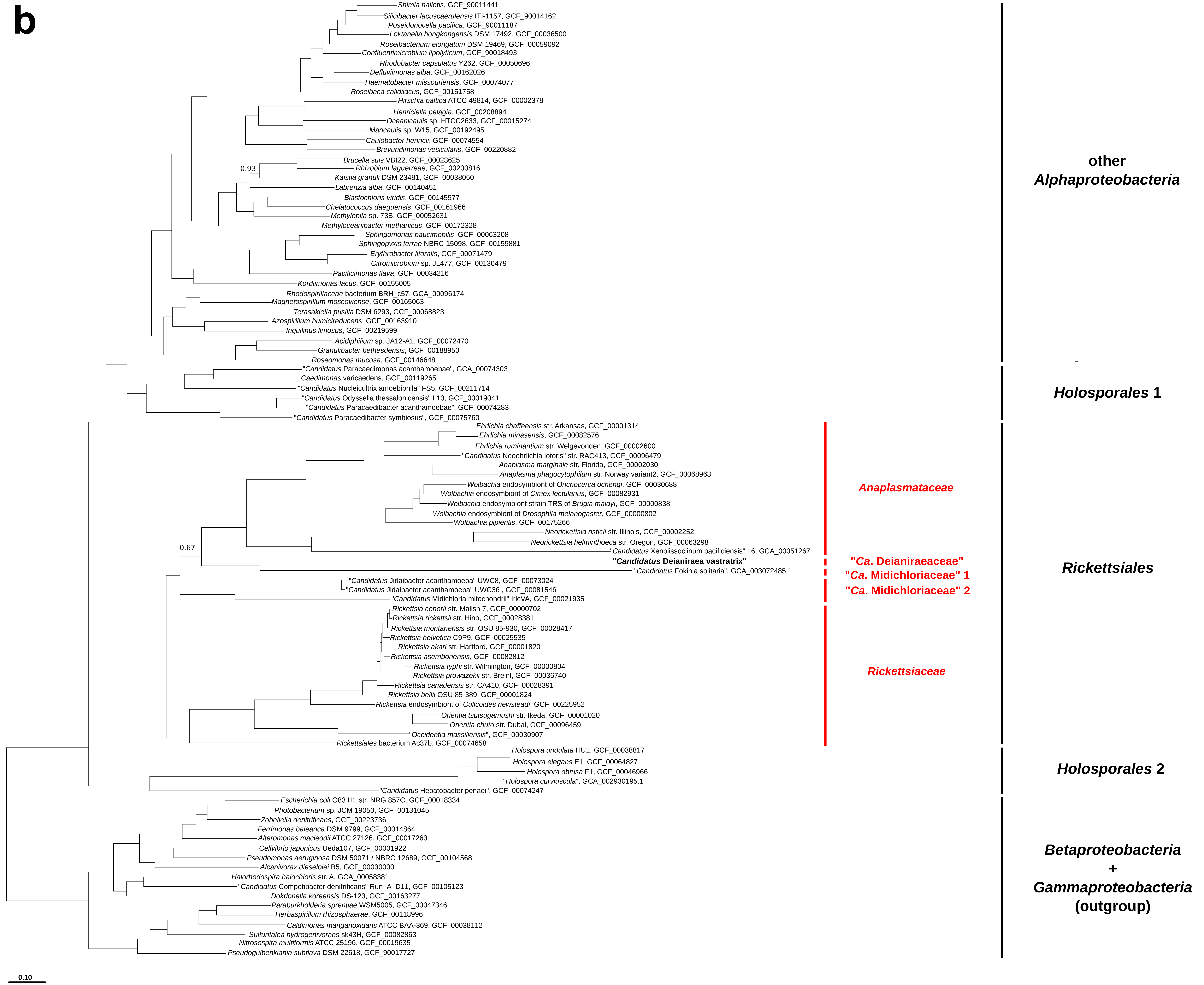

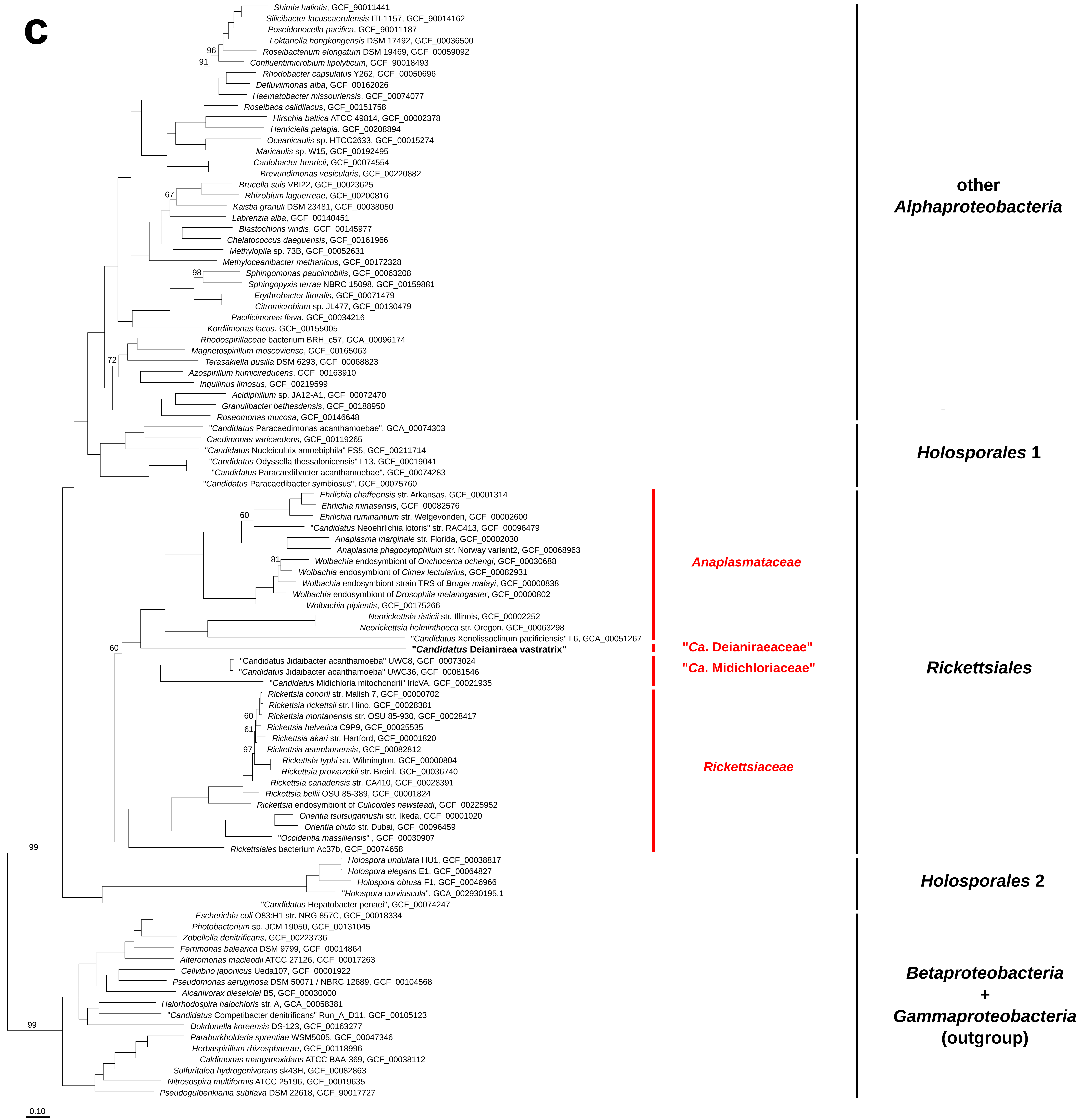

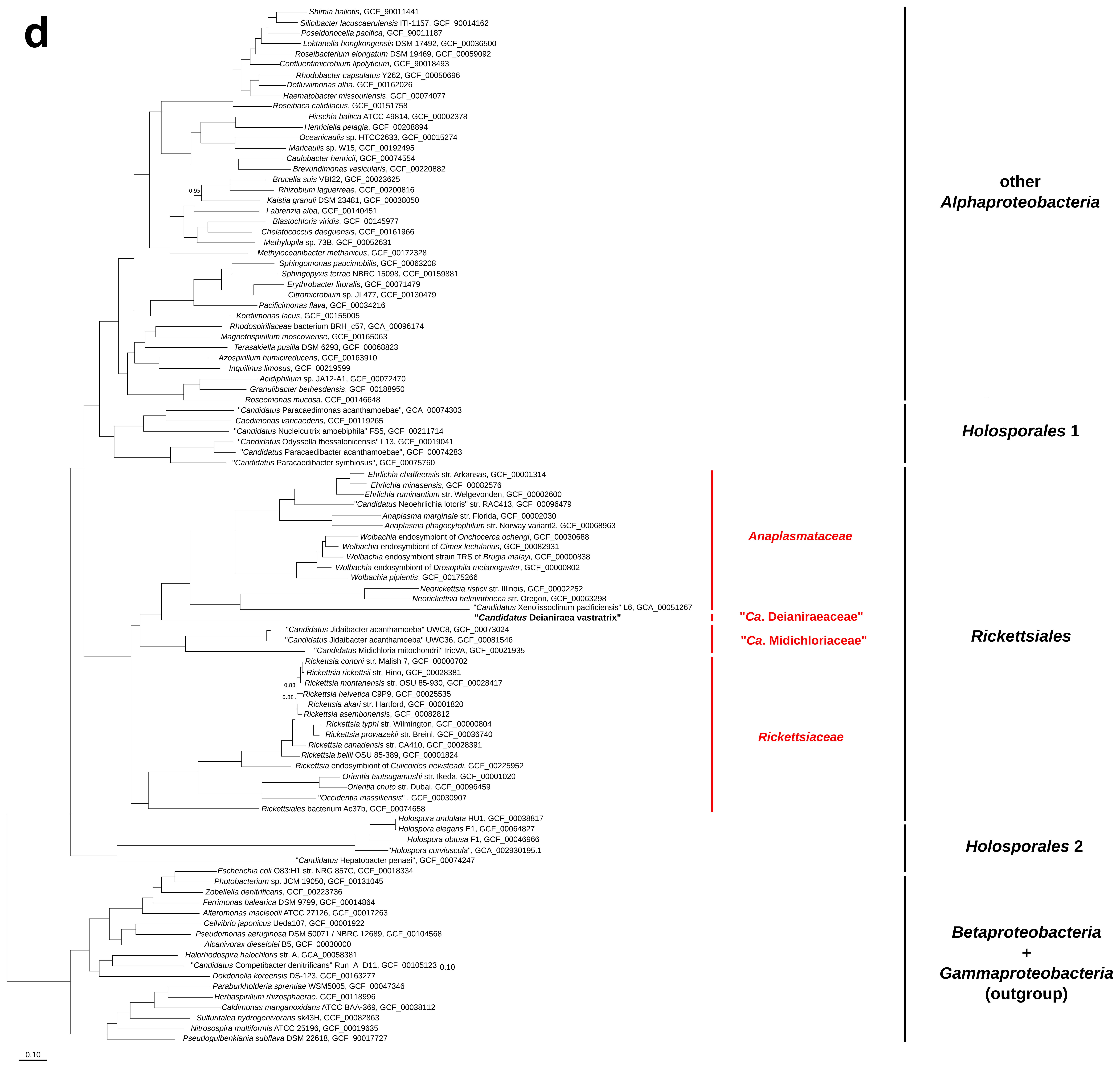

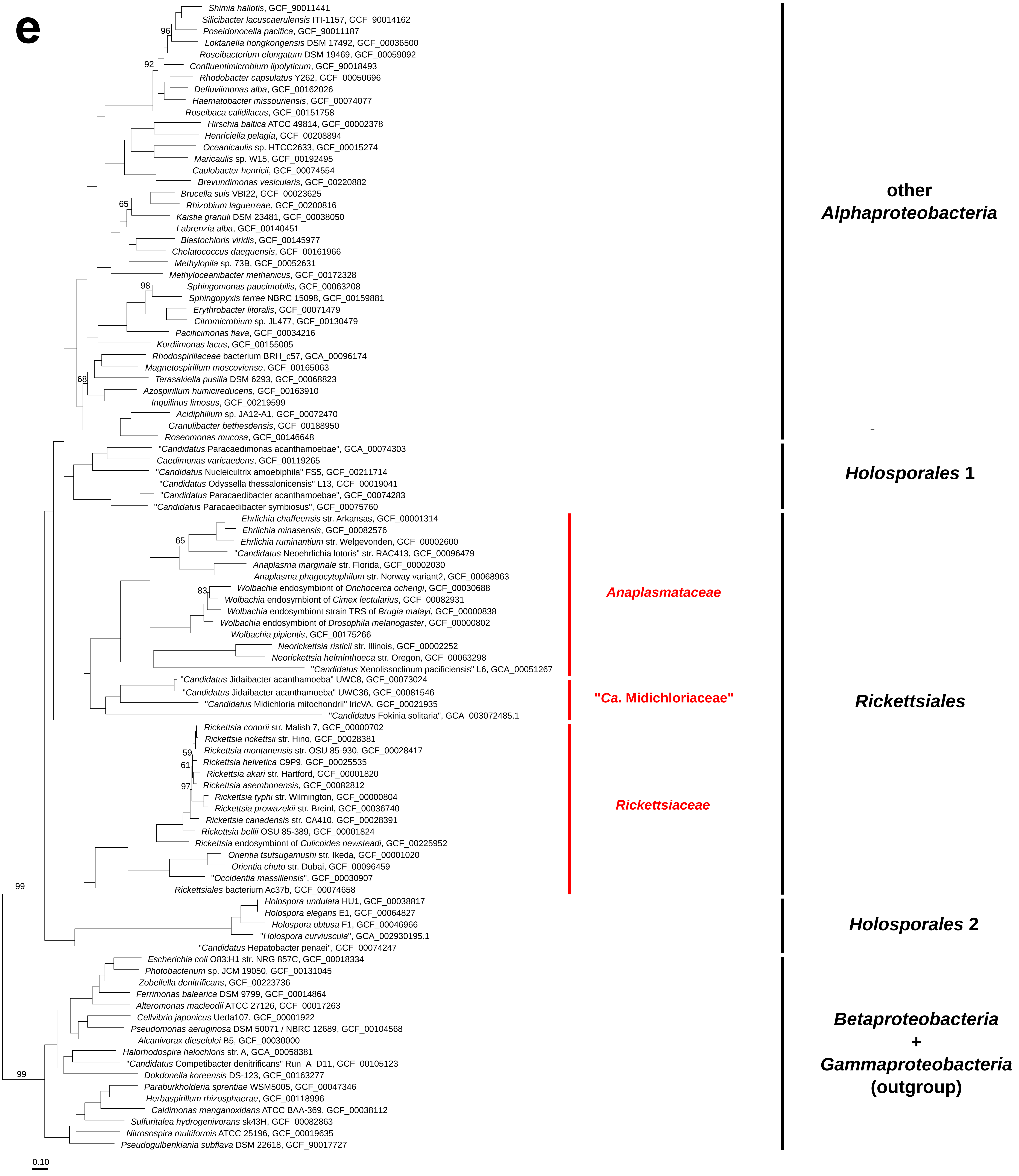

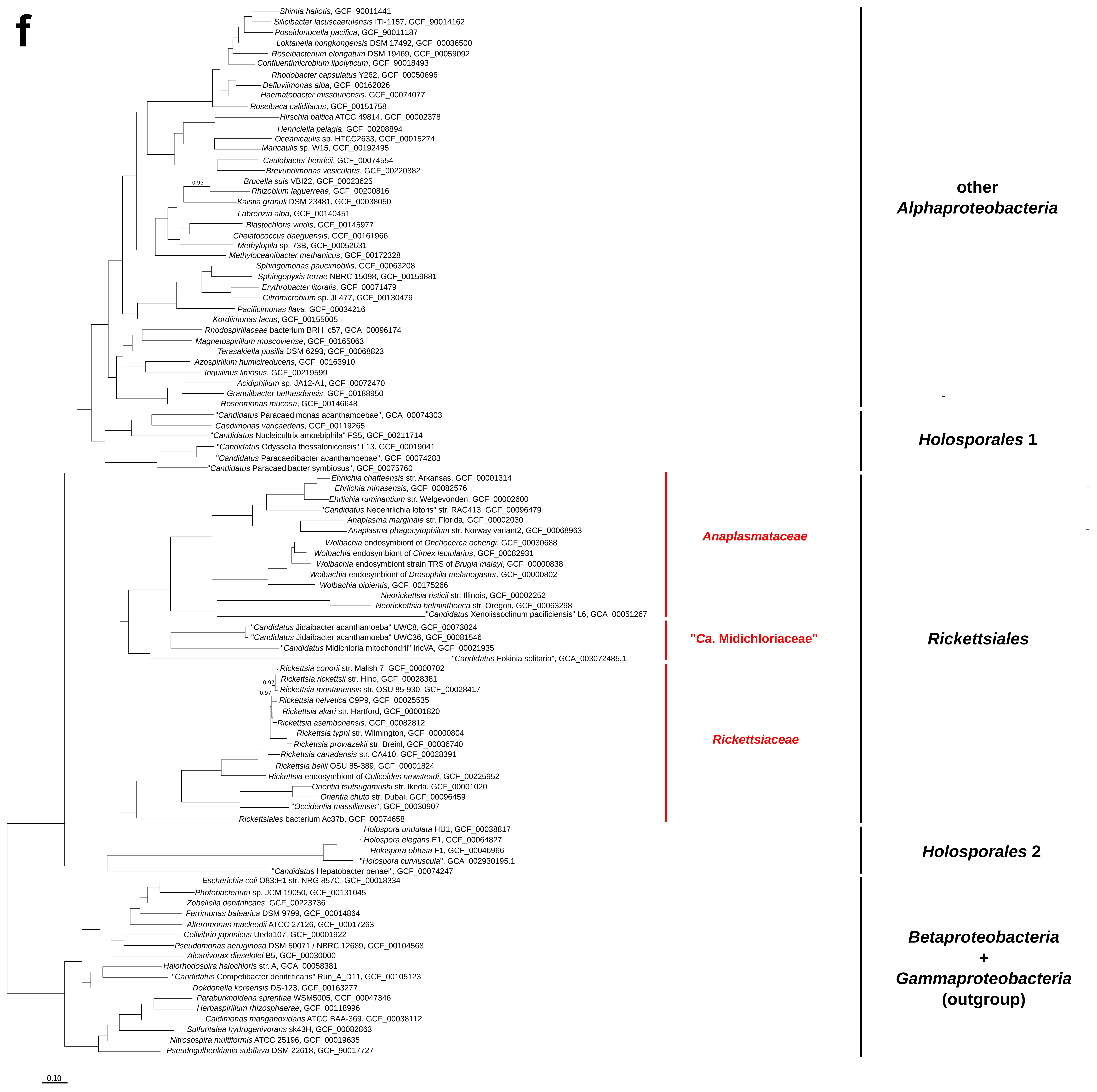

### Suppl_18_Amino_acid_pathways.pdf

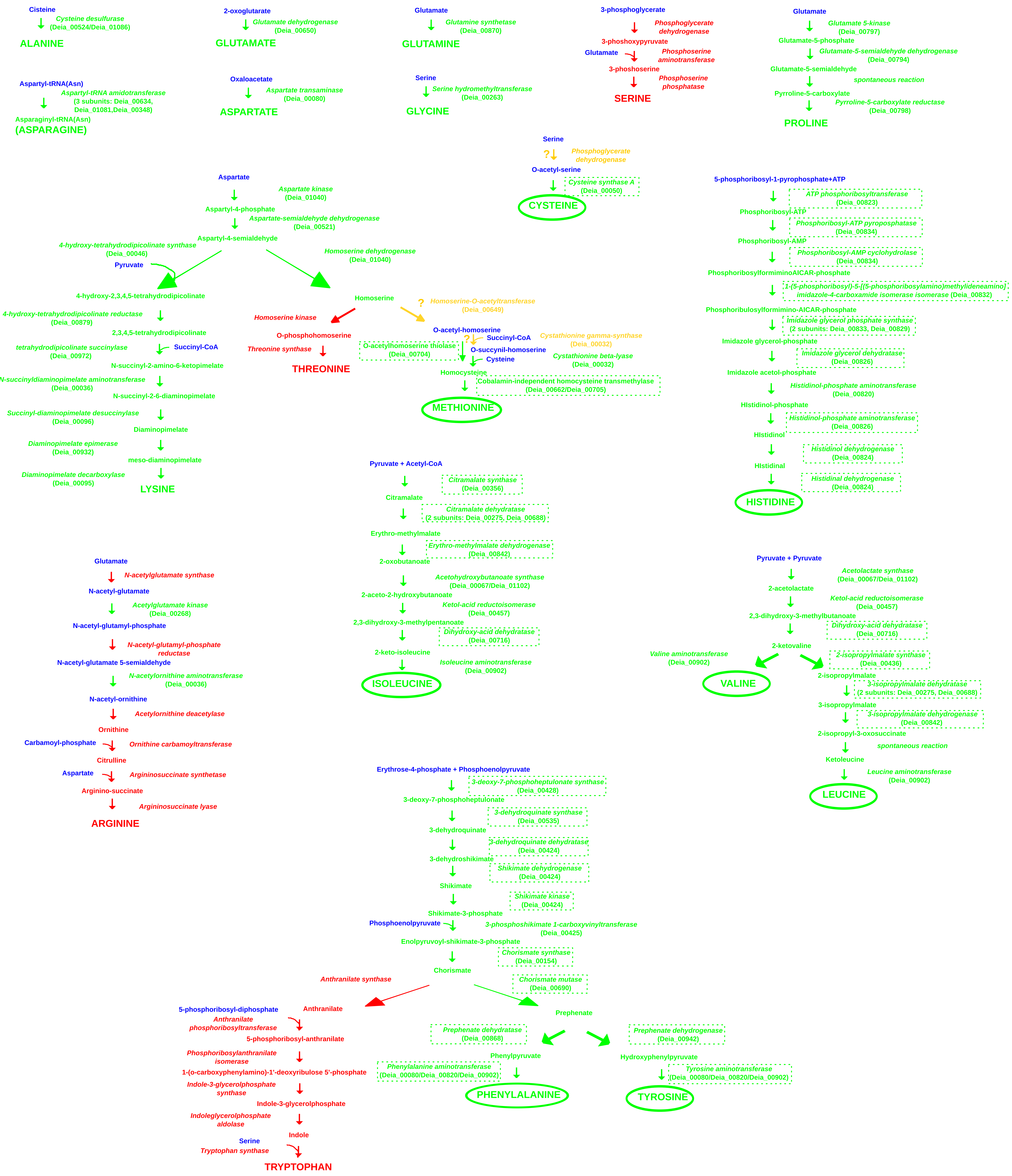

### Suppl_21_Phylogeny_selected_genes.pdf

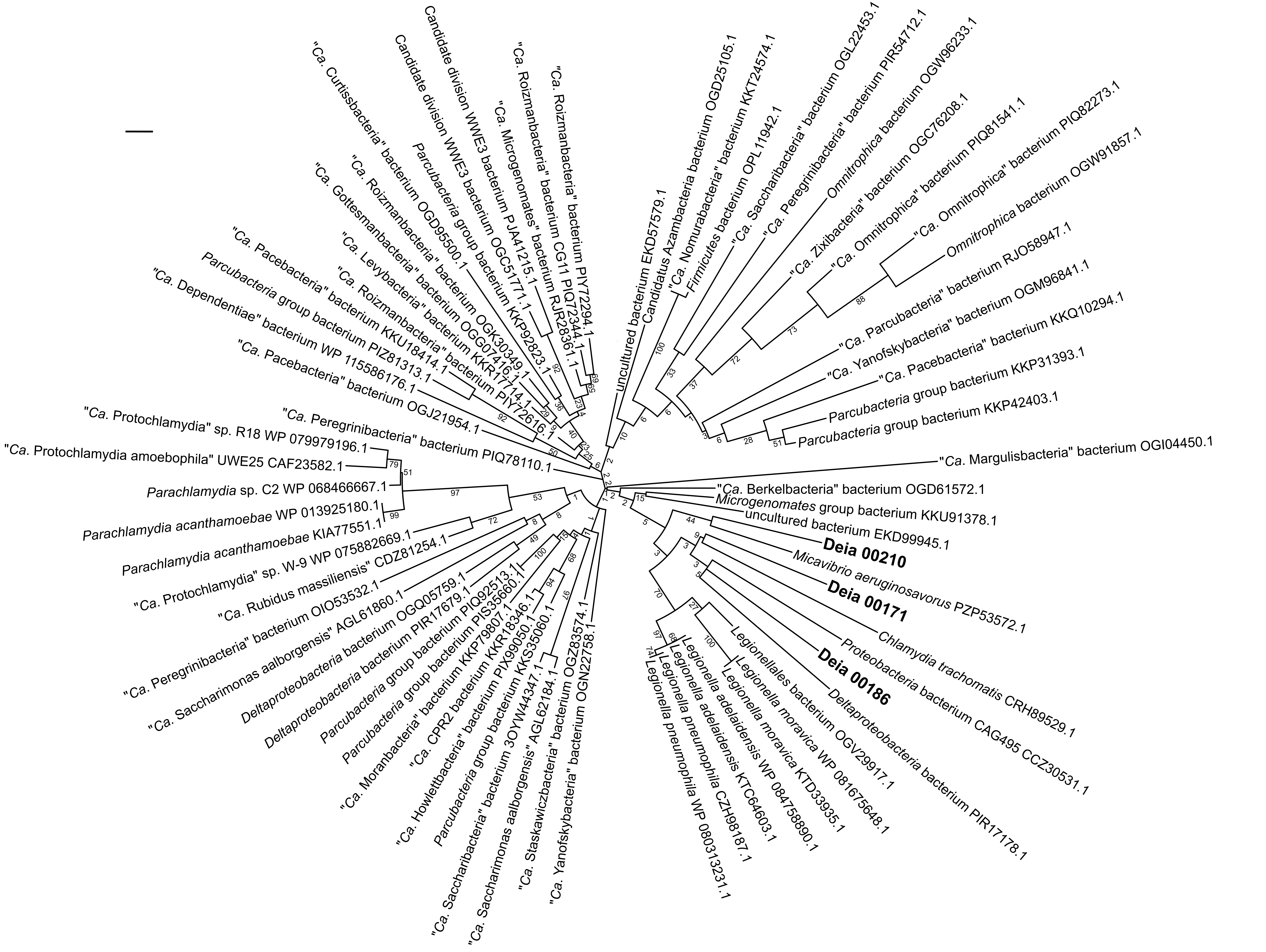

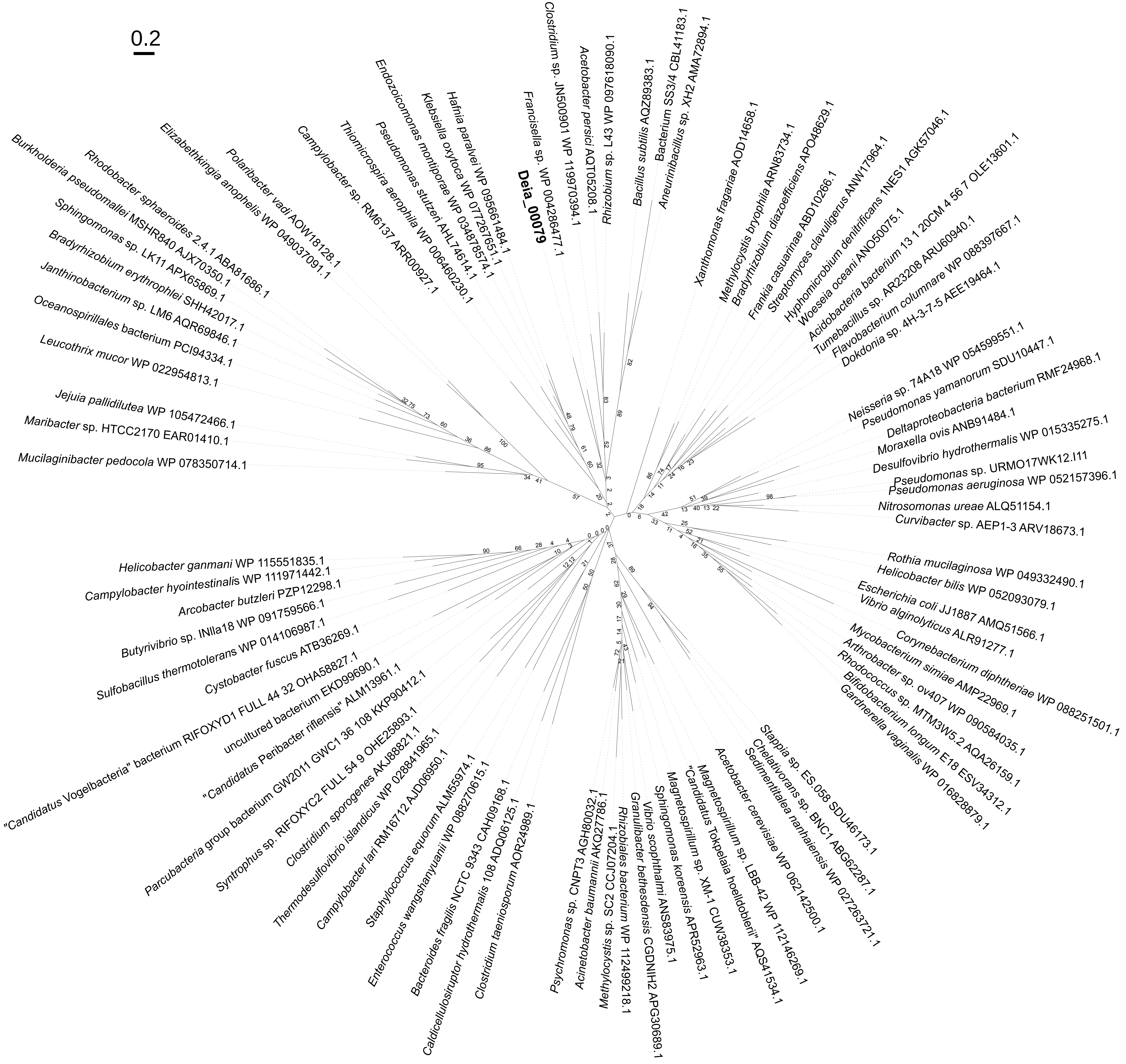

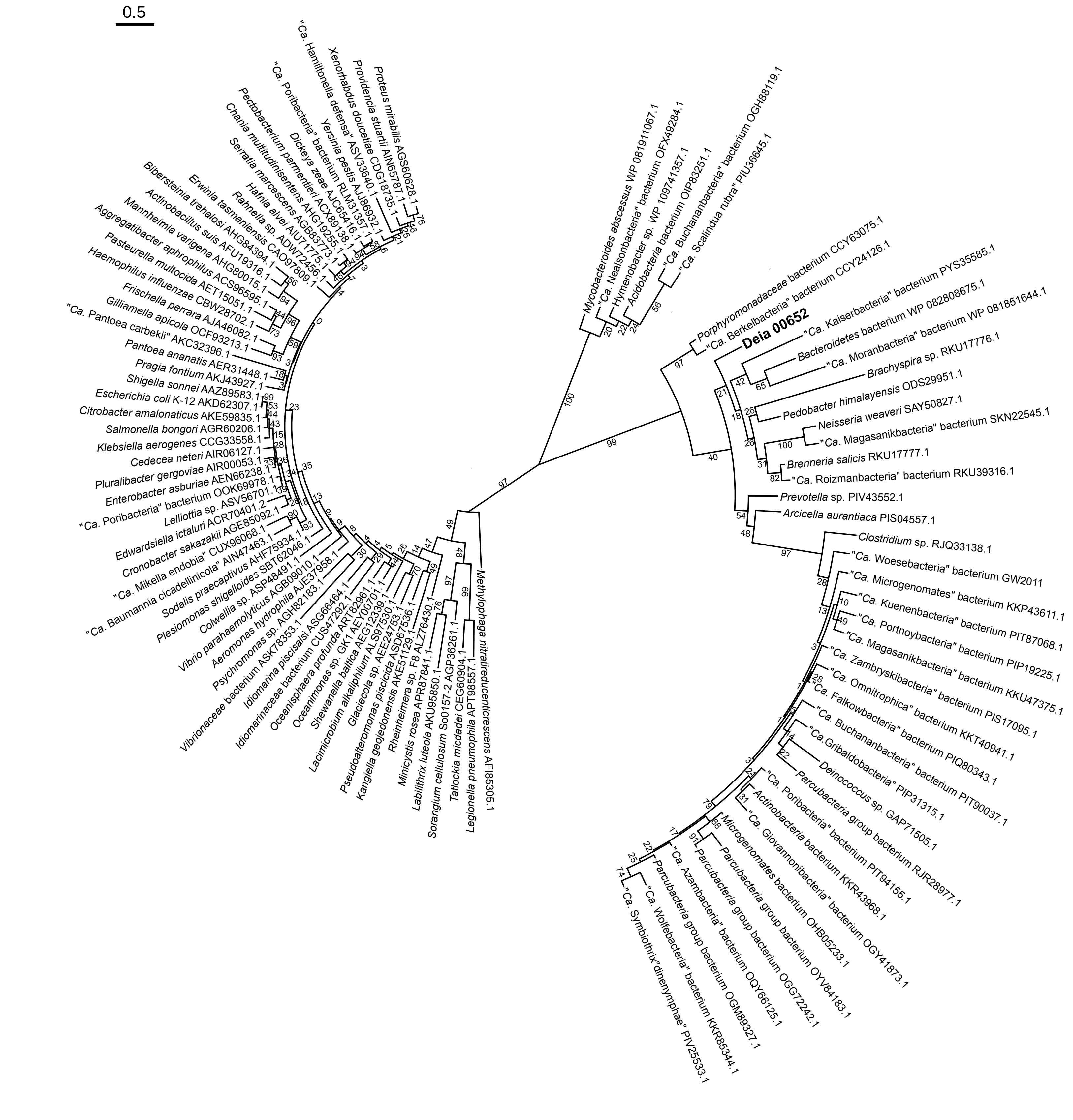

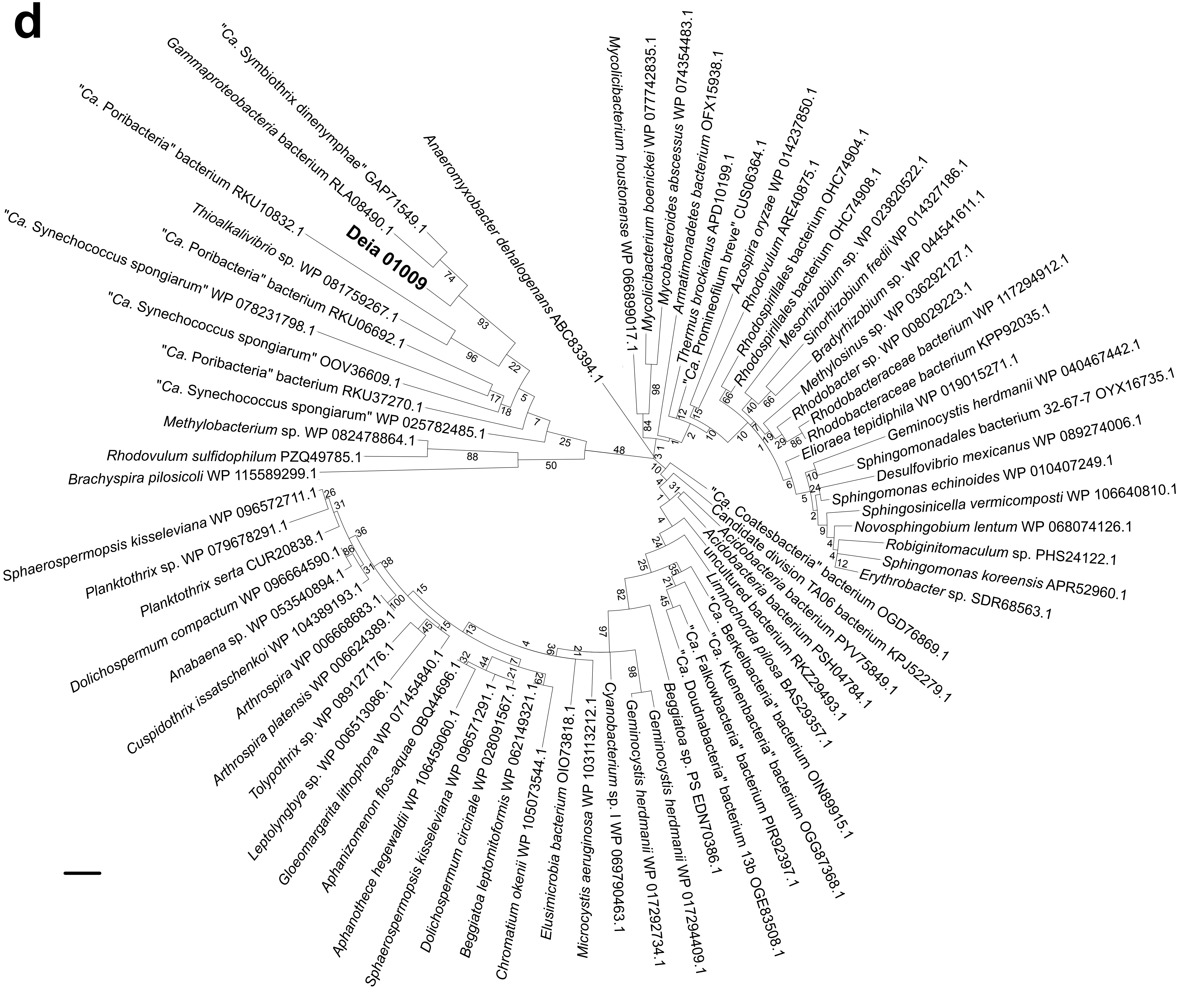
